# Estimating indirect parental genetic effects on offspring phenotypes using virtual parental genotypes derived from sibling and half sibling pairs

**DOI:** 10.1101/2020.02.21.959114

**Authors:** Liang-Dar Hwang, Justin D Tubbs, Justin Luong, Mischa Lundberg, Gunn-Helen Moen, Pak C Sham, Gabriel Cuellar-Partida, David M Evans

## Abstract

Indirect parental genetic effects may be defined as the influence of parental genotypes on offspring phenotypes over and above that which results from the transmission of genes from parents to children. However, given the relative paucity of large-scale family-based cohorts around the world, it is difficult to demonstrate parental genetic effects on human traits, particularly at individual loci. In this manuscript, we illustrate how parental genetic effects on offspring phenotypes, including late onset diseases, can be estimated at individual loci in principle using large-scale genome-wide association study (GWAS) data, even in the absence of parental genotypes. Our strategy involves creating “virtual” mothers and fathers by estimating the genotypic dosages of parental genotypes using physically genotyped data from relative pairs. We then utilize the expected dosages of the parents, and the actual genotypes of the offspring relative pairs, to perform conditional genetic association analyses to obtain asymptotically unbiased estimates of maternal, paternal and offspring genetic effects. We develop a freely available web application that quantifies the power of our approach using closed form asymptotic solutions. We implement our methods in a user-friendly software package **IMPISH** (**IM**puting **P**arental genotypes **I**n **S**iblings and **H**alf-Siblings) which allows users to quickly and efficiently impute parental genotypes across the genome in large genome-wide datasets, and then use these estimated dosages in downstream linear mixed model association analyses. We conclude that imputing parental genotypes from relative pairs may provide a useful adjunct to existing large-scale genetic studies of parents and their offspring.

## Introduction

There is increasing interest in estimating the indirect effect of parental genotypes on the phenotypes of their offspring (Bates et al. 2018; Evans et al. 2019; Kong et al. 2018; Lawlor et al. 2017). We and others have shown in human populations that the maternal and paternal genomes can indirectly affect a range of offspring traits including perinatal (Beaumont et al. 2018; Evans et al. 2019; Tyrrell et al. 2016; Warrington et al. 2019; Warrington et al. 2018; Yang et al. 2019; Zhang et al. 2015; Zhang et al. 2017; Zhang et al. 2018) and later life phenotypes (Kong et al. 2018; Warrington et al. 2019). However, these sorts of analyses typically require large numbers of genotyped parent-offspring duos and trios in order to partition genetic effects into parental and offspring mediated components (Evans et al. 2019; Moen et al. 2019). Unfortunately, there are only a few cohorts around the world with large numbers of genotyped parents and children (Boyd et al. 2013; Fraser et al. 2013; Krokstad et al. 2013; Magnus et al. 2006; Sudlow et al. 2015) implying that even if investigators were able to combine all the large-scale family-based cohorts in the world together, the statistical power to resolve parental genetic effects on offspring phenotypes may be low (Moen et al. 2019). The problem of low statistical power is exacerbated further if the interest is on identifying parental genetic effects on late onset diseases, since many of the cohorts that contain genotypic information on parents and their children are birth cohorts that were established less than thirty years ago (Boyd et al. 2013; Fraser et al. 2013; Magnus et al. 2006). This means that offspring from these cohorts are not old enough to have developed many late onset diseases of interest. There is therefore a considerable need to develop statistical genetics methods and software that can maximize the amount of data available to detect indirect parental genetic effects (Evans et al. 2019).

In the following manuscript, we describe a simple strategy for estimating indirect parental genetic effects on offspring phenotypes which is capable of leveraging the considerable information contained within large publicly available cohorts and the tens of thousands of individuals contained within twin registries and family studies from around the world (Silventoinen et al. 2015). Briefly, our strategy involves creating “virtual” mothers and fathers by estimating the genotypic dosages of parental genotypes using physically genotyped data from sibling and half sibling relative pairs. We then use the expected dosages of the parents, and the actual genotypes of the siblings/half sibling pairs to perform conditional genetic association analyses and estimate maternal, paternal and offspring genetic effects on the offspring phenotype.

We derive formulae to impute the expected dosage of maternal and paternal genotypes given sibling or half-sib genotypes at both autosomal and X-linked loci. We implement our calculations in a user-friendly software package, **IMPISH** (**IM**puting **P**arental genotypes **I**n **S**iblings and **H**alf siblings) that allows users to quickly and efficiently impute parental genotypes across the genome in large genome-wide datasets, and then use these estimated dosages in downstream genome-wide association analyses (http://evansgroup.di.uq.edu.au/software.html). We investigate the statistical power, type 1 error and bias associated with estimating parental and offspring genetic effects via simulation and using closed form asymptotic solutions. Finally, we develop a series of freely available web applications (http://evansgroup.di.uq.edu.au/power-calculators.html) that researchers can use to estimate power to detect parental and offspring genetic effects in studies of sibling or half sibling pairs, with or without parental genotypes.

## Methods

### Imputing Expected Gene Dosages for Parents Given Observed Offspring Genotypes

The intuition for why relative pairs enable imputation of parental genotypes is illustrated in Figure 1. Essentially, an individual’s sibling/half sibling provides additional information on the likely genotype of their parents- so that some parental genotypes are more probable than others given the observed genotype data. For example in Figure 1, it is possible to conclude that both parents of siblings who have genotypes “AA” and “aa” at an autosomal locus must be heterozygous. Likewise, maternal half siblings (i.e. half siblings who share a common mother) whom have genotype “AA” and “aa” at an autosomal locus, imply that their shared mother must be genotype “Aa” and their fathers “AA” or “Aa” and “Aa” or “aa” respectively (the exact probabilities depending on the allele frequencies at the locus under consideration). We calculated the probability of maternal and paternal biallelic SNP genotypes given data from sibling pairs or half sibling pairs at the same locus. We did this for autosomal and non-pseudoautosomal X chromosome loci for bi-allelic SNP markers using Bayes Theorem e.g. for an autosomal locus:

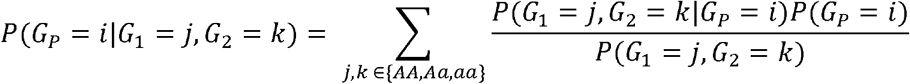

where *G*_*P*_ ∈ {“AA”, “Aa”, “aa”} refers to the genotype of the parent, and *G*_*1*_ and *G*_*2*_ the genotypes of offspring one and two. In the case of full sibling pairs, separate maternal and paternal genotypes can be resolved for X-linked loci. However, in the case of autosomal loci, the expected dosage for maternal and paternal genotypes is the same, meaning that it is impossible to distinguish maternal from paternal genotypes.

**Figure 1.**
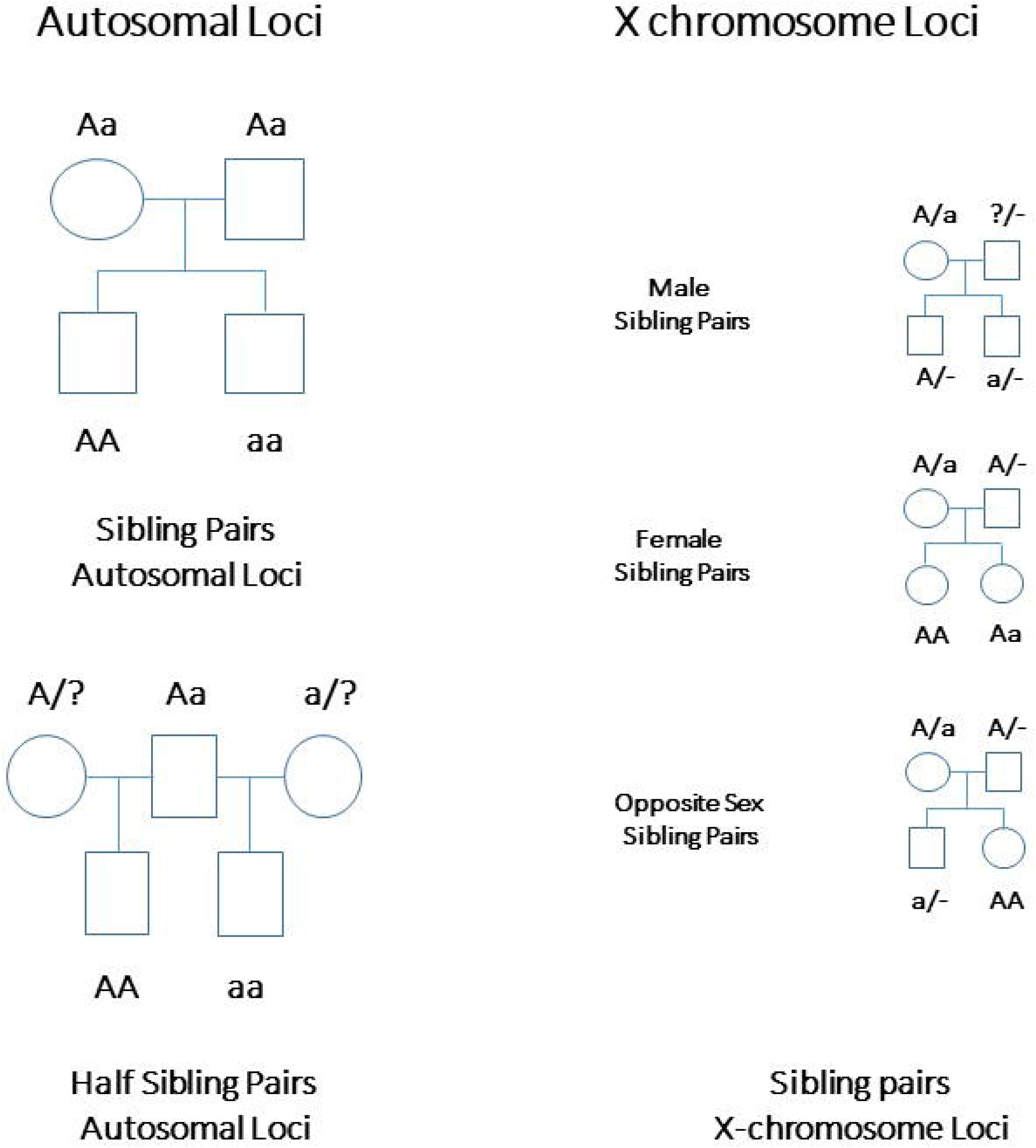
Illustration showing the intuition behind why the genotypes of relative pairs such as siblings and half siblings provide information on parental genotypes. In the case of sibling pairs at autosomal loci, sibling genotypes provide information on parental genotypes. However, mothers and fathers have the same expected genotypes and so separate maternal and paternal genotypes cannot be imputed given only genotype information from sibling pairs. However, mothers and fathers have different expected genotypes given sibling pair genotypes at non-autosomal X chromosome loci, and so different maternal and paternal genotypes can be imputed at these loci. Likewise, in the case of half sibling pairs, mothers and fathers have different expectations for their genotypes given half sibling genotypes, and so maternal and paternal genotypes can be imputed at loci. Male individuals are uninformative for paternal genotypes at (non-pseudoautosomal) X chromosome loci.

Given conditional genotype probabilities, it is then a simple matter to calculate the expected genotype dosages (*X*) of the parents for a given pair of offspring genotypes:

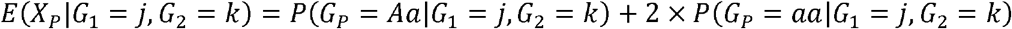

These expected maternal and/or paternal dosages can then be included as terms in the fixed effects part of a linear mixed model together with the observed dosages of the offspring genotypes and then estimates of maternal, paternal and fetal genetic effects on the offspring phenotype can be obtained. Derivations of expected parental genotype dosages given sibling/half sibling genotypes for autosomes and the X chromosome are given in Supplementary Tables 1-4.

### Exploring Parameter Bias, Power and Type 1 Error of Tests of Genetic Association via Simulation

We investigated parameter bias, power and type 1 error rate of tests of genetic association via simulation. Genotypes were simulated for nuclear families (mother, father and two siblings) and maternal half sibling families (common mother, two fathers and two half siblings). In the case of sibling pairs at autosomal loci, trait values were simulated according to the following model:

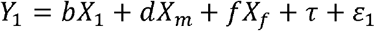

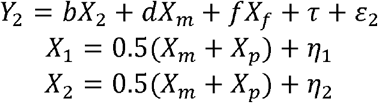

where *Y*_*1*_ and *Y*_*2*_ are the phenotypes of siblings one and two, *X*_*1*_, *X*_*2*_, *X*_*m*_ and *X*_*f*_ ∈ {0, 1, 2} are the genotype dosages of siblings one and two and their mother and father respectively, *b*, *d* and *f* are the effect of the offspring, maternal and paternal genotypes on the offspring phenotype, *τ* is a random effect shared by the siblings, *ε*_1_ and *ε*_2_ are uncorrelated error terms for the two phenotypes, and *η*_1_ and *η*_2_ are random effects due to the segregation of alleles. Without loss of generality, effects were scaled so that the variance of the genotype dosages and phenotype terms was one. The variances of the random effects are:

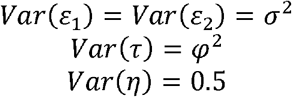

In the case of X chromosome loci for sibling pairs, we assume that the effect of genotypes on offspring phenotype are equal in males and females (i.e. the regression coefficients *b*, *d* and *f* are equal regardless of whether the sibling is male or female). Unstandardized female X ∈ {0, 1, 2} whilst unstandardized male genotypes are coded {0, 2}. This means that male genotypes have twice the variance of female genotypes, and explain double the variance in the offspring phenotype. We simulated sibling phenotypes at X chromosomal loci under the following model:

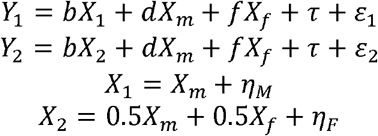

where the terms are defined similar to the sibling model above where sibling one is male and sibling two is female, and *η*_M_ and *η*_F_ are random effects due to segregation in male and female offspring. The variances of the random effects are:

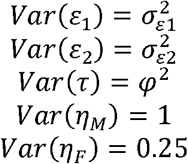

At X chromosome loci, the covariances between genotypes of relative pairs are sex-dependent:

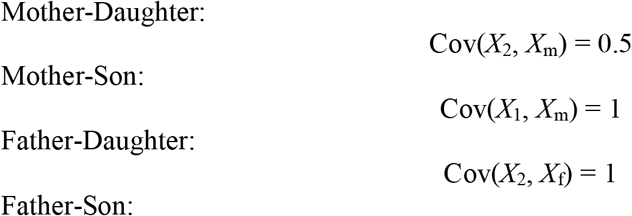

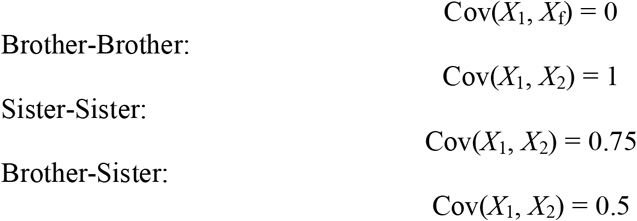

and so we simulate under three separate models for female-female, male-male, and opposite sex sibling pairs (see Supplementary Materials for more details).

Finally, in the case of (maternal) half sibling pairs at autosomal loci we simulated data according to the model:

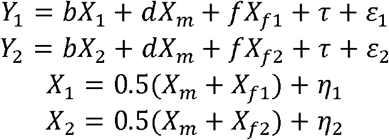

where the subscripts *f1* and *f2* denote the fathers of half sibling one and half sibling two respectively. The variances of the random effects are

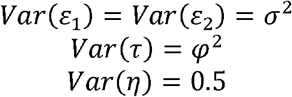

Paternal half sibling pairs can be parameterized analogously. The reason we don’t show this explicitly is that for autosomal loci, the power to detect maternal effects using paternal half sibling pairs is the same as the power to detect paternal effects using maternal half sibling pairs, and the power to detect paternal effects using paternal half sibling pairs is the same as the power to detect maternal effects using maternal half sibling pairs.

For each simulated pair, we calculated the expected genotype dosages of the parents based on the formulae from the preceding section (Supplementary Tables 1 to 4). Offspring phenotype was regressed on offspring genotype, and imputed (or physically genotyped) parental dosages using the *lmer* package in R. Tests were conducted using full information maximum likelihood. In the case of sibling pairs at autosomal SNPs, we investigated the properties of the following tests of association:

1. Omnibus test: We compared the full model where free terms for the offspring and parental genetic effect(s) were estimated versus a model where the offspring and parental regression coefficient(s) were fixed to zero (i.e. either a two degrees of freedom test if parental genotype was imputed; or a three degree of freedom test if genotypes for both parents were available).
2. Test using offspring genotypes only: We compared a model where there was a free term for the offspring genetic effect only, against a model where this term was set to zero (i.e. a one degree of freedom test). In other words, the effect of parental genotypes was not modelled in this analysis, even though parental genetic effects may influence the offspring phenotypes and parental genotypes may or may not be present.
3. Test of the offspring genetic effect: We compared the full model where free terms for the offspring and parental genetic effect were estimated versus a model where the offspring regression coefficient was fixed to zero (a one degree of freedom test).
4. Test of the parental genetic effect: We compared the full model where free terms for the offspring and parental genetic effect are estimated versus a model where the imputed parental regression coefficient was fixed to zero (a one degree of freedom test).

In the case of half sibling pairs, as well as sibling pairs at X chromosome SNPs, we investigated the properties of the following tests of association:

1. Omnibus test: We compared the full model where free terms for offspring, maternal and paternal genetic effects are estimated versus a model where the offspring, maternal and paternal regression coefficients were fixed to zero (a three degrees of freedom test).
2. Test using offspring genotypes only: We compared a model where there was a free term for the offspring genetic effect only, against a model where this term was set to zero (a one degree of freedom test). In other words, the effect of parental genotypes were not modelled in this analysis, even though maternal and/or paternal genetic effects may influence the offspring phenotypes and parental genotypes may or may not be present.
3. Test of the offspring genetic effect: We compared the full model where free terms for offspring, maternal and imputed paternal genetic effects are estimated versus a model where the offspring regression coefficient was fixed to zero (a one degree of freedom test).
4. Test of the maternal genetic effect: We compared the full model where free terms for offspring, maternal and paternal genetic effects are estimated versus a model where the maternal regression coefficient was fixed to zero (a one degree of freedom test).
5. Test of the paternal genetic effect: We compared the full model where free terms for offspring, maternal and paternal genetic effects are estimated versus a model where the paternal regression coefficient was fixed to zero (a one degree of freedom test).

In the case of the Omnibus test (Model 1) and the test using the offspring genotypes only (Model 2), the focus is on locus detection (i.e. whether there is a genetic effect at the locus, regardless of whether it is mediated through the offspring or parental genomes). In contrast, in the case of the tests for the parental, maternal, paternal or offspring genetic effects (Models 3 to 5), the focus is on partitioning a known locus into its indirect parental genetic and/or offspring genetic components. These tests are more relevant if the goal is to determine which genome mediates a known genetic effect on the offspring phenotype, or if the objective is on deriving unbiased effect estimates of genetic effects e.g. for Mendelian randomization analyses.

For our simulations, we varied the size of genetic effects (two conditions: *b*^*2*^ = *d*^*2*^ = *f*^*2*^ = 0; *b*^*2*^ = *d*^*2*^ = *f*^*2*^ = 0.1%), frequency of the trait decreasing allele (three conditions: *p* = 0.1, *p* = 0.5, *p* = 0.9), and shared residual variance (two conditions: *φ*^2^ = 0; *φ*^2^ = 0.2). For all simulations we used N = 2000 sibling pairs/half sibling pairs, a type 1 error rate of α = 0.05, and 1000 replications. R code implementing the simulations are provided in the Supplementary Materials.

### Calculating Power Analytically Using the Non-Centrality Parameter

We derived closed form expressions for the non-centrality parameter of the statistical tests described above for actual and imputed parental genotypes and confirmed the results of these against our simulations. We have implemented these asymptotic power calculations in a series of applications which are freely available on our website (http://evansgroup.di.uq.edu.au/power-calculators.html). In the results section, we use our utilities to compare the statistical power to detect genetic effects when parental genotypes are available and when they need to be imputed for both sibling and half sibling pairs.

### Software to impute Parental Genotypes

We have coded the parental imputation routines described above in a C++ software package called **IMPISH** (**IM**puting **P**arental genotypes in **S**iblings and **H**alf siblings) which is freely available on our website (http://evansgroup.di.uq.edu.au/software.html). IMPISH uses source code adapted from the GCTA software package (version 1.26.0) that has been modified to impute parental genotype data given genotypes from sibling or half sibling pairs (Yang et al. 2010). IMPISH accepts data in the form of PLINK style binary .bed, .bim and .fam file formats (Purcell et al. 2007). Users can elect to output expected parental genotype dosages or have the software compute these internally and utilize them in genome-wide association testing. IMPISH fits a genetic mixed linear model with fixed effects for offspring genotype and (imputed) maternal and paternal genotypes and allows users to compute these statistics across the genome in a computationally efficient fashion. A genome-wide genetic relationship matrix is used in the random effects part of the model just as in the original GCTA software, allowing users to account for population stratification and cryptic relatedness in their analyses.

To quantify the computational requirements of the **IMPISH** software, we simulated datasets that ranged in size from *N* = 1,000 to 20,000 sibling pairs and *M* = 500,000 autosomal SNP markers. The datasets were simulated using an approach similar to that described above. We benchmarked the running time and memory use of the **IMPISH** software by running simulations on these datasets. Reported runtimes are the medians of five identical runs in a computing environment with 256 GB memory and 16 CPU cores with solid-state disk in one compute node.

## Results

### Simulation Results

A summary of the results of our data simulations is presented in Supplementary Figure 1. Estimates of paternal, maternal and offspring genetic effects from the full omnibus models were unbiased, even when imputed parental genotypes were used in the place of real genotypes. Type 1 error rates were also maintained at expected levels (Supplementary Tables 5-9). Estimates of statistical power, closely matched those from asymptotic calculations (Supplementary Tables 5-9 and see below).

### Derivation of non-centrality parameters and asymptotic power for tests of association in sibling pairs (autosomal loci)

Under full information maximum likelihood, all the tests of association considered in this manuscript are distributed as non-central chi-square distributions under the alternative hypothesis of genetic association, with degrees of freedom equal to the difference in the number of free parameters between full and reduced models. The non-centrality parameter (*ζ*) of these distributions is equal to twice the difference in expected log-likelihoods between the full and reduced models. Given the non-centrality parameter (*ζ*) of the statistical test, the power to detect association (P) can be obtained by the formula:

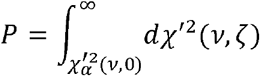

where 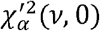 i the 100(1 − α) percentage point of the central *χ*^2^ distribution with ν degrees of freedom, and 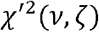, denotes a non-central chi-square distribution with non-centrality parameter *ζ* and degrees of freedom ν. In the section below, we derive the expected covariance matrix of the residuals for each statistical model and its associated expected minus two log-likelihood. From these values the non-centrality parameter and statistical power of the relevant test of association can be calculated.

To illustrate our derivations, we consider the case of sibling pairs with phenotype data *Y*_1_ and *Y*_2_, and corresponding genotype data *X*_1_ and *X*_2_, at an autosomal single nucleotide polymorphism (SNP). Similar derivations for sibling pairs at X chromosome loci and for half sibling pairs on the autosomes are provided in the Supplementary Materials. The coding of genotype assumes additivity (i.e. no dominance), and without loss of generality, all genotypes and phenotypes are standardized to have mean 0 and variance 1. In situations where the genotype data of parents (i.e. paternal genotype *X*_f_ and maternal genotype *X*_m_) are unavailable, the maternal and paternal genotypes are imputed from the genotypes of the sibling pairs as:

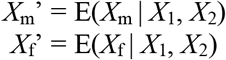

We assume the model above for sibling pairs (see Methods) and random mating so that Cov(*X*_*m*_, *X*_*f*_) = 0. The covariances between genotypes are:

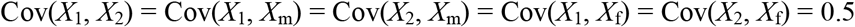

the covariance between phenotypes and genotypes are:

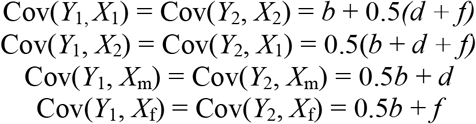

and the covariance between the two phenotypes is:

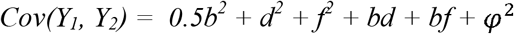

The phenotypic variance can be decomposed as follows:

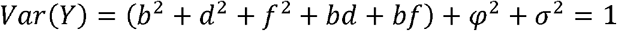

The variance of the true maternal and paternal genotypes prior to standardization is equivalent to the expected heterozygosity, given the allele frequencies *p* and *q*:

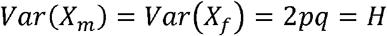

Full sibling relationships do not provide adequate information to distinguish between alleles of maternal versus paternal origin, therefore the imputed maternal genotype 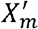 will be equivalent to the imputed paternal genotype 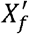. Thus, models using the imputed parental genotype 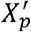, estimate the parameter *c*, the combined effects of *d* and *f* such that *c* = *d* + *f*.

The variance of the imputed parental genotype, 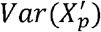, imputed from full sibling pairs is:

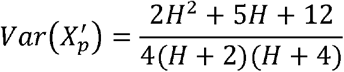

The covariance between actual and imputed genotype is equal to the variance of the imputed genotype:

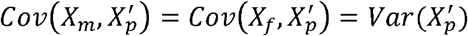

and the covariance between the imputed parental genotype and sib genotypes and phenotypes are:

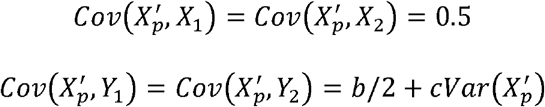

When actual maternal and paternal genotypes are available, the linear mixed model is

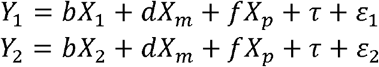

The fixed effects *b*, *d*, and *f* may be estimated by generalised least squares (GLS), where the covariance matrix of random effects is:

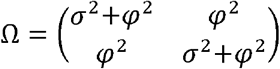

The inverse of the covariance matrix of random effects is:

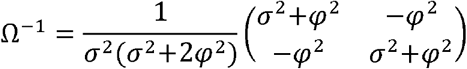

The asymptotic GLS estimates of a vector of parameters *β* are given by We consider the following models:

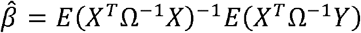

We consider the following models:

#### Null model of no association

The residual covariance matrix is simply the covariance matrix of Y:

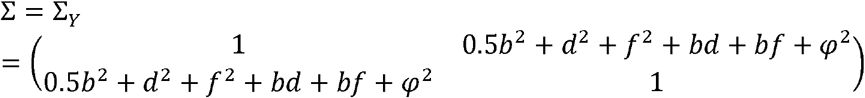

The expected minus two log-likelihood (−2lnL) of the model per sibling pair is therefore:

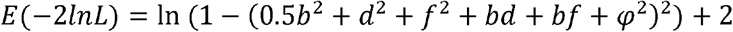

#### One parental genotype (maternal genotype) in the model

The *X* matrix contains one column with elements *X*_*m*_. The asymptotic GLS estimate of the regression coefficient of *X*_*m*_ is:

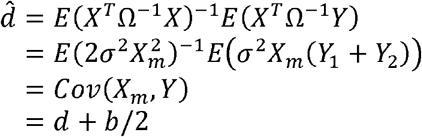

The residual covariance matrix is:

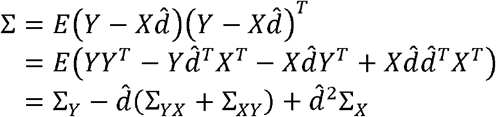

where

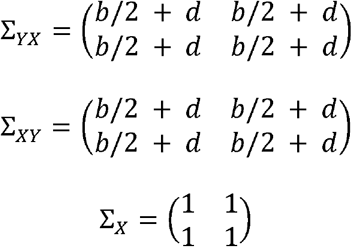

Therefore:

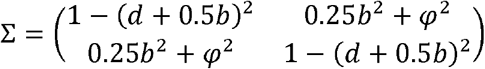

The expected −2lnL of the model per sibpair is therefore:

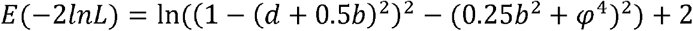

#### Both parental genotypes only in model (terms for *X*_*m*_ and *X*_*f*_ only)

The *X* matrix contains two columns with elements *X*_*m*_ and *X*_*f*_. The asymptotic GLS estimates for the regression coefficients of *X*_*m*_ and *X*_*f*_ are:

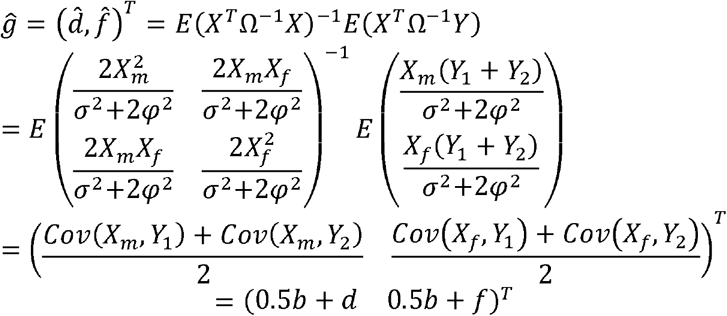

The residual covariance matrix is:

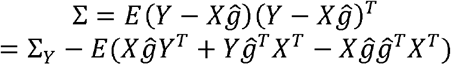

where

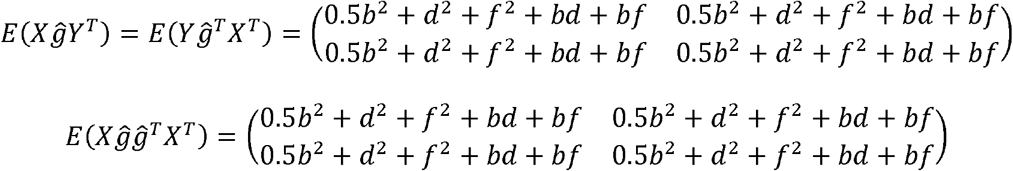

Therefore:

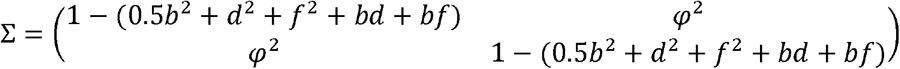

The expected −2lnL of the model per sibling pair is therefore:

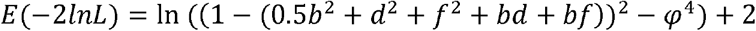

#### Offspring genotypes in model only (terms for *X*_1_ and *X*_2_ only)

The *X* matrix contains one column with elements *X*_1_ and *X*_2_. The asymptotic GLS estimate of the regression coefficient of *X*_1_ and *X*_2_ is:

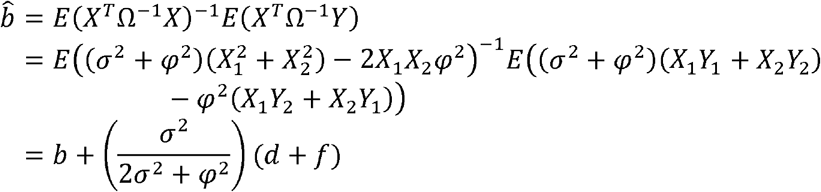

The residual covariance matrix is:

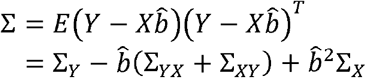

where:

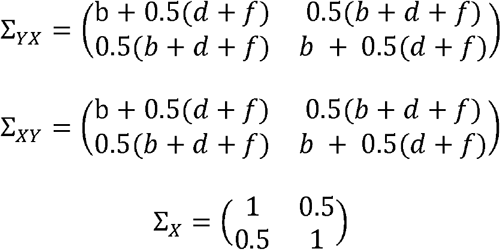

Therefore:

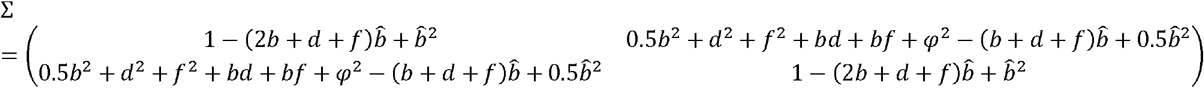

The expected −2lnL of the model per sibling pair is:

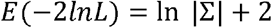

#### Full Omnibus Model (terms for *X*_*m*_, *X*_*f*_, *X*_1_ and *X*_2_)

The *X* matrix contains three columns; column 1 with elements *X*_1_ and *X*_2_, column 2 with elements *X*_*m*_ and *X*_*m*_ and column 3 with elements *X*_*f*_ and *X*_*f*_. The asymptotic GLS estimate of the regression coefficients of columns 1, 2 and 3 are:

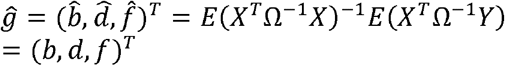

The residual covariance matrix is:

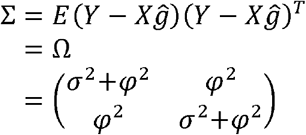

The expected −2lnL of the model per sibling pair is:

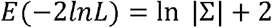

#### Imputed parental genotypes only in model (terms for 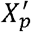 only)

When only imputed parental genotypes are available, the linear mixed model becomes:

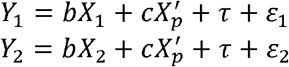

The *X* matrix contains one column with elements 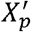. The asymptotic GLS estimate of the regression coefficient of 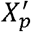 is:

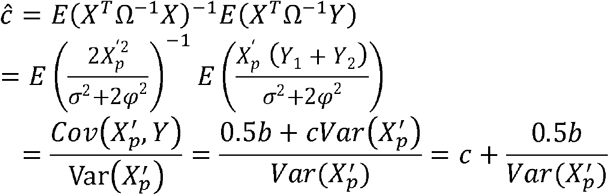

The residual covariance matrix is:

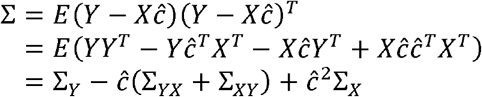

where:

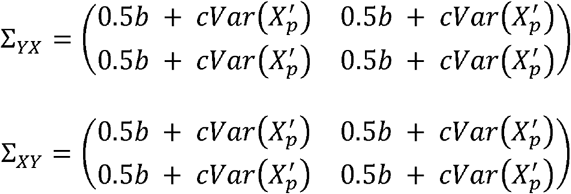

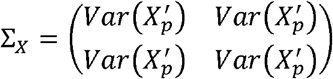

The expected −2lnL of the model per sibling pair is:

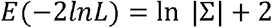

#### Full omnibus model with imputed parental genotypes (terms for 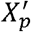, *X*_1_ and *X*_2_

The *X* matrix contains two columns; column 1 with elements *X*_1_ and *X*_2_, and column 2 with elements 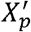 and 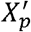. The asymptotic GLS estimate of the regression coefficients of columns 1 and 2 are:

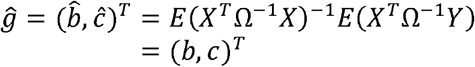

The residual covariance matrix is:

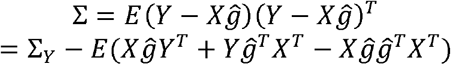

where:

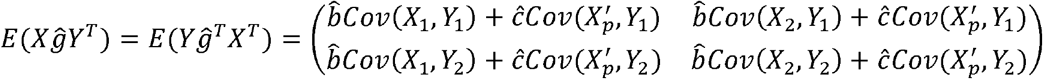

and:

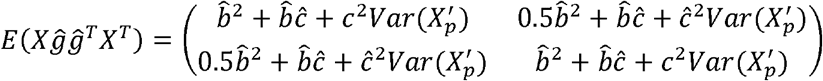

The expected −2lnL of the model per sibling pair is:

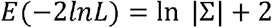

### Results of Asymptotic Power Calculations

We used our asymptotic formulae to investigate the statistical power to detect association across a range of different parameters, study designs and statistical tests (Supplementary Tables 10-14). We highlight some general results from our power calculations that we hope investigators may find useful in terms of planning genetic association studies, particularly those aimed at identifying and/or estimating the contribution of indirect parental genetic effects on offspring phenotypes.

A key question for researchers is, what is the optimal analysis strategy if the primary focus is on locus detection? According to our power calculations, the answer to this question, perhaps unsurprisingly, depends on the genetic architecture of the trait, in particular on the existence of indirect maternal/paternal genetic effects and whether these are in the same or opposing directions. Figure 2 displays power to detect a locus using sibling pairs when a locus is influenced by maternal and/or offspring genetic effects (e.g. a perinatal trait like birth weight). When the locus under study involves an offspring genetic effect only (green lines in Figure 2), which is probably the case for the majority of loci in the genome for most traits, then the most powerful strategy appears to be simply testing for an offspring genetic effect against the null model of no association (i.e. performing a one degree of freedom test just using the sibling pairs with no parental imputation). This includes situations where parents have been genotyped. This is because fitting the full omnibus model and testing against the null model requires extra degrees of freedom to model parental genetic effects (which in this case are not present) which adversely affects power. We note that this decrement does not appear to be great in the case of sibling pairs if only the mother is genotyped and paternal genetic effects are not modelled and do not contribute to the trait of interest, (Figure 2, Supplementary Table 10)- which is perhaps a reasonable assumption for many perinatal phenotypes.

**Figure 2.**
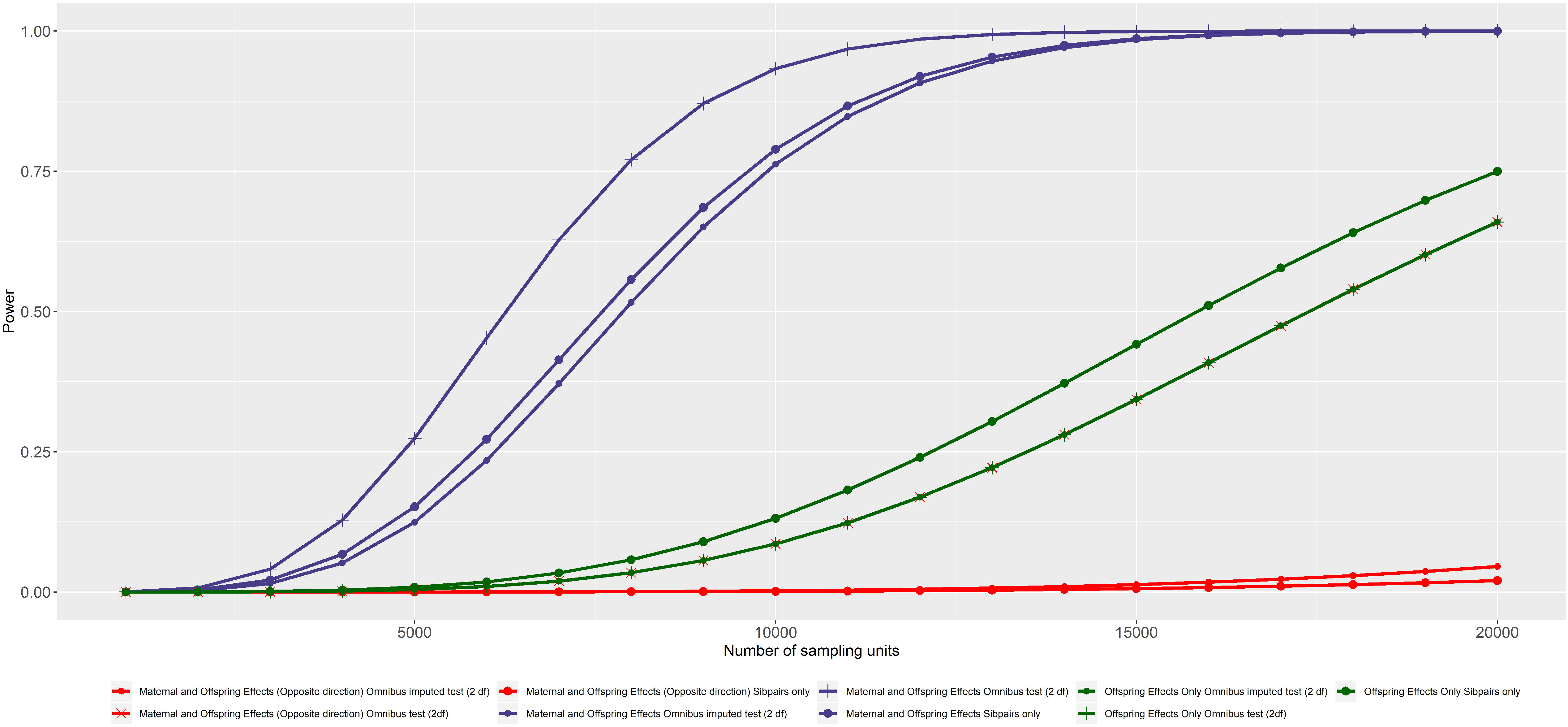
Power of locus detection in sibling pairs assuming directionally concordant maternal and offspring genetic effects (blue lines: *d*^*2*^ = 0.1%; *b^2^* = 0.1%), directionally discordant maternal and offspring genetic effects (red lines: *d*^*2*^ = 0.1%; *b*^*2*^ = 0.1%), or offspring genetic effects only (green lines: *d*^*2*^ = 0%; *b*^*2*^ = 0.1%). Shown are results of a one degree of freedom test using sibling genotypes only (lines with large circles), an omnibus two degree of freedom test using observed maternal and sibling genotypes (lines with crosses), and an omnibus two degree of freedom test of association when parental genotypes need to be imputed from sibling genotypes (lines with small circles). We note that the power of three conditions are equivalent (i.e. offspring effects only omnibus imputed and omnibus tests; discordant maternal and offspring effects omnibus test). To understand this result intuitively, the variance in the phenotype explained under the offspring genetic effects only model (*b*^*2*^ = 0.1%; d^*2*^ = 0%) is the same as that explained under the directionally discordant model because of the negative covariance between maternal and offspring genotypes (i.e. variance explained = *b*^*2*^ + *d*^*2*^ – *bd* = 0.1% + 0.1% − 0.1% = 0.1%). Likewise, the power is also identical under the omnibus test when parental genotypes are imputed because the effect in this condition is completely driven by the offspring genotype (i.e. the presence/absence of maternal genotypes does not contribute to the model fit). For all calculations we assume an autosomal locus, shared variance *φ*^2^ = 0.2, a type 1 error rate α = 5×10^−8^, and where relevant, a decreasing allele frequency of p = 0.1.

In contrast, when indirect maternal (or paternal) genetic effects substantially influence the offspring phenotype (blue and red lines Figure 2), and parental genotypes are present, the full omnibus model (lines with crosses) often performs comparably or better than a simple one degree of freedom test using the sibling genotypes alone (lines with small circles). This is especially the case when offspring and/or parental genetic effects are directionally discordant (red lines), as is frequently observed for some trait-locus combinations like fasting glucose associated loci and offspring birth weight (Warrington et al. 2019; Warrington et al. 2018). Here the power of a simple one degree of freedom test involving the sibling genotypes only can be vastly diminished, because the discordant parental and offspring genetic effects tend to cancel each other out. In contrast, an omnibus test which models both offspring and indirect parental genetic effects performs much better in these situations. Importantly, when parental genotypes are unavailable, for many situations there appears to be little gained by imputing parental genotypes and including these in an omnibus test if the focus is solely on locus detection (Figure 2; Supplementary Tables 10-11).

Another goal investigators might be interested in is partitioning effects at known genetic loci into direct offspring and indirect parental genetic components. This may be of relevance if investigators want to prove the existence of indirect maternal genetic effects on offspring phenotypes for example. Figure 3 displays the power to partition a genetic effect into maternal (or equivalently paternal) genetic sources of variation in the case of half sibling or sibling pairs, with and without parental genotypes at autosomal loci. The graph highlights the clear advantage in power of including actual as opposed to imputed parental genotypes in the statistical model when the focus is on resolving indirect genetic effects on offspring phenotypes. Figure 3 shows that if parental genotypes are unavailable, then a considerable number of sibling pairs (>40,000) and maternal half sibling pairs (>60,000) will be required to achieve high power (>80%, α = 0.05) to partition genetic effects at a locus- even for those of relatively large effect (*d* ^2^ = 0.1%). Interestingly, paternal half sibling pairs who have not had their parents genotyped, provide much less power to estimate maternal genetic effects and require even larger numbers (a similar decrement in power is also observed in the case for maternal half sibling pairs if the interest is in estimating paternal genetic effects). The lower power of the imputed half sibling analyses compared to the imputed sibling analyses partially reflects the fact that only two sources of variation are modelled in the imputed sibling models (i.e. offspring and parental genetic sources of variation), whereas in the half sibling models, three different sources of variation are modelled (offspring, parental, maternal genetic sources of variation). If investigators believe that paternal genetic effects do not contribute to offspring trait variation (a reasonable assumption for perinatal traits), then one option to increase power is to fix this path to zero in models involving half siblings. Interestingly, the presence/absence of other genetic effects has little effect on power of the conditional tests of association for realistic effect sizes when correctly modelled.

**Figure 3.**
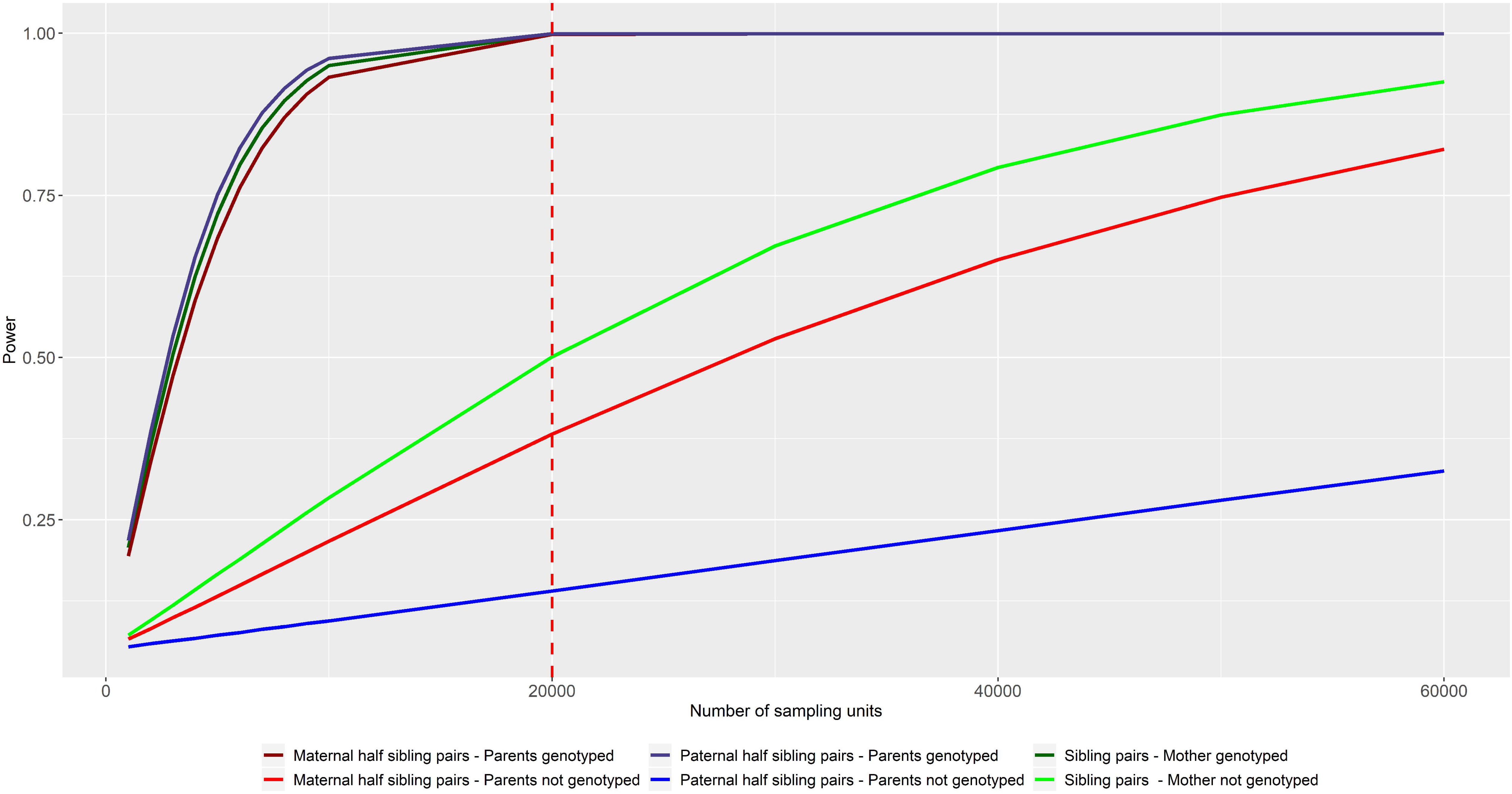
Power to resolve an autosomal maternal genetic effect (*d*^*2*^ = 0.1%; *f*^*2*^ = 0%; *b*^*2*^ = 0%;) at a known genetic locus, using a conditional one degree of freedom test of association in sibling pairs (green lines), maternal half sibling pairs who share the same mother (red lines) and paternal half sibling pairs who share the same father (blue lines). All calculations assume p = 0.3 frequency of the trait decreasing allele; shared variance *φ*^2^ = 0.2; type 1 error rate α = 0.05). The red dashed vertical line in the figure indicates the approximate number of sibling pairs in the UK Biobank (N = 20,000). This figure highlights the advantage of having actual parental genotypes in the statistical model.

In order to put the above numbers in context, in the publicly available UK Biobank dataset (which roughly consists of ~20,000 sibling pairs), we estimate that an autosomal parental genotype would need to explain ~0.2% of the variance in the offspring phenotype in order to have 80% power to resolve an indirect genetic effect if parental genotypes need to be imputed (assuming the same parameters as in Figure 3). An indirect effect size this large is probably unrealistic for most traits, implying that larger samples will be needed to resolve genetic effects at known loci into indirect and direct genetic effects if parental genotypes need to be imputed. We also note that the power of the conditional tests are typically lower than omnibus tests, implying that omnibus tests of association should be used for locus discovery purposes whilst conditional tests of association should be reserved for partitioning effects/estimating effect sizes at known loci (Supplementary Tables 10-14).

We found that the effect of the other parameters we investigated (allele frequency, shared residual variance) on statistical power was complicated and often interacted with the level of other factors in the calculation (Supplementary Tables 10-11). Allele frequency exerted a modest effect on the power of most of the statistical tests examined, and its effect on power appeared to be symmetric around p = 0.5. The effect of the shared residual variance on statistical power was complex and depended on the statistical test, the underlying genetic model, allele frequency etc (Supplementary Tables 10-11).

Imputing parental genotypes on the X chromosome has the advantage that separate maternal, paternal and offspring genetic effects can be resolved for sibling pairs (although at X linked loci, male siblings are uninformative for paternal transmissions, and so contribute nothing in terms of identifying paternal genetic effects when fathers have not been genotyped). We coded *G*_*1*_ ∈{0, 2} and female genotypes are coded *G*_*2*_ ∈{0, 1, 2}. We also assume that the regression coefficient of offspring phenotype on (maternal/paternal/offspring) genotype is the same in male and female offspring. This means that male loci explain double the amount of variance in the phenotype compared to females (see Supplementary Materials). We have coded the web utilities (http://evansgroup.di.uq.edu.au/power-calculators.html) so that users enter the variance in the offspring phenotype explained by maternal, paternal and/or offspring genotypes at the locus. For offspring genetic effects in opposite sex siblings, users enter the variance explained by male loci.

The results of the power analyses for sibling pairs on the X chromosome are displayed in Supplementary Tables 12-14. The general pattern of results for loci on the X chromosome was similar to that described for the autosomes, and consequently we make similar recommendations regarding the appropriate analyses for locus detection and partitioning genetic effects at X linked loci. Comparing the power across the different study designs however revealed a few interesting results which we highlight. First, male sibling pairs offer increased power to resolve indirect paternal genetic effects on the X chromosome compared to the autosomes- so long as fathers have been genotyped. This is because (under random mating) father’s genotype is uncorrelated with maternal and male offspring genotype at X linked loci. Correlation between paternal X linked loci and male offspring phenotype can therefore not be explained by maternal or offspring genotype. The corollary is that male sibling pairs cannot be used to impute paternal genotypes at X linked loci and so are uninformative for paternal genetic effects unless the father has been genotyped (opposite sex sibling pairs also provide slightly elevated power to detect paternal genetic effects when fathers have been genotyped for the same reason).

When parental genotypes are present, male siblings also provide lower power to detect X linked loci (3 degree of freedom tests) compared to many of the other study designs. The reason is the converse of the explanation above- paternal genetic effects do not contribute to the covariance between male sibling pairs. Opposite sex sibling pairs also provide reduced power to detect loci (3 degree of freedom tests), but this partly a consequence of how we parameterize the model of association on the X chromosome (i.e. we calculate the size of offspring genetic effects in reference to the variance explained by male offspring, meaning that the variance explained by the same locus in the sister will be half this amount). We choose not to compare results across the different study designs when parental genotypes are imputed because the different models and their tests are usually not equivalent (e.g. one can’t resolve paternal genotypes at X linked loci for male sibling pairs; opposite sex sibling pairs have their variances parameterized slightly differently to the other sibships etc). These results are tabulated in Supplementary Tables 12-14.

### IMPISH Software Performance

Supplementary Table 15 shows the performance of the **IMPISH** software in terms of CPU times and time to perform genome-wide association. Our results show that **IMPISH** can be used to impute parental genotypes from large numbers of relative pairs and perform tests of association across the genome in a reasonable time frame.

## Discussion

In this manuscript we have shown that it is possible to impute parental genotypes given genotype data on sibling and half sibling pairs and then subsequently use this information to derive unbiased estimates of parental genetic effects. We have derived asymptotic formulae to compute power to detect association when parental genotypes are observed or imputed, and implemented these calculations in a series of freely available online power calculators that investigators can use to guide study design and analysis strategy. Finally, we developed **IMPISH**, a freely available easy to use software package that imputes parental genotypes given genotype information on sibling or half sibling pairs, and then uses this information to perform genome-wide conditional tests of paternal, maternal and offspring genetic effects.

Our asymptotic calculations reveal that the power to partition known individual loci into parental and offspring genetic effects using imputed parental genotypes is low in general, and highlights the value in having parents genotyped if the interest is in resolving indirect parental genetic effects at known loci. In situations where parental genotypes are unavailable, we show that indirect parental genetic effects can still be estimated without bias, but very large numbers of sibling (or half sibling) pairs will be required (e.g. >40,000 sibling and >60,000 half sibling pairs). Whilst these sorts of numbers may be realistic in the case of siblings (e.g. UK biobank contains roughly 20,000 sibling pairs, and there are many twin cohorts around the world that contain large numbers of dizygotic twins), most cohorts contain very few half sibling pairs. For these reasons we suggest our method may currently be more suitable as a complement to existing large-scale genetic studies of parents and their children. For example, both the Norwegian MOBA and HUNT cohorts not only contain tens of thousands of parent-offspring trios and duos, but also large numbers of sibling pairs that could be combined with more traditional parent-offspring analyses to further increase power to detect parental genetic effects (Krokstad et al. 2013; Magnus et al. 2006).

A key motivation for developing our approach was the realization that estimates of parental genetic effects derived from imputed genotypes could also be used in two sample Mendelian randomization (MR) studies examining possible causal relationships between parental exposures and offspring outcomes (Evans et al. 2019). Whilst our method could be used to increase the power of existing MR analyses involving perinatal outcomes (Tyrrell et al. 2016), an exciting novel application would be the examination of the influence of parental exposures on later life offspring outcomes. The majority of the world’s large-scale cohorts with genotyped mother-offspring pairs are relatively new historically (Boyd et al. 2013; Fraser et al. 2013; Magnus et al. 2006; Wright et al. 2013). This means that the children in these cohorts are not old enough to have developed many late onset diseases of interest including adverse cardiometabolic phenotypes. Consequently, it is currently difficult, if not impossible, to perform maternal-offspring MR studies on late onset diseases. Our procedure of imputing parental genotypes means that in principle mother-offspring MR analyses are now possible utilizing cohorts of mature sibling and half sibling pairs. Such an approach would enable the investigation of hypotheses in life course epidemiology such as the Developmental Origins of Health and Disease which posits a link between intrauterine growth restriction and the development of disease in the offspring in later life (Barker 1990).

Besides low statistical power, there are a number of limitations with our approach. In the case of sibling pairs, separate maternal and paternal genotypes can be resolved for X linked (non-pseudoautosomal) loci. However, for autosomal loci, the expected dosage for maternal and paternal genotypes is the same. This means that it is impossible to distinguish maternal and paternal genotypes using data from sibling pairs alone. Thus, utilization of sibling pairs to detect indirect genetic effects requires the non-trivial assumption that either paternal (or maternal) genetic effects do not affect the offspring phenotype under study. Whilst this assumption may be justified for certain perinatal phenotypes where the contribution of the father’s phenotype to trait variation in the offspring may be minimal (like birth weight), it may not be justifiable for other phenotypes. Sensitivity analyses could be performed by testing whether estimates derived from using sibling pairs are consistent with those derived from e.g. parent-offspring trios or even half sibling pairs where estimates of maternal, paternal and offspring genetic effects can be estimated consistently.

We have shown that for half sibling pairs, different maternal and paternal genotype probabilities (and therefore expected dosages) can be resolved at genetic loci. This means that, in principle, the half sibling pairs within large publicly available biobanks could be leveraged to provide information on parental genotypes and consequently help obtain unbiased estimates of indirect parental genetic effects on offspring traits. This will be possible if there is explicit pedigree information that unequivocally identifies half sibling relationships. However, the task becomes more challenging if half siblings have to be identified on the basis of genetic information alone. This is because half siblings share the same expected number of alleles identical by descent as grandparent-grandchild pairs and avuncular relationships, making it difficult to distinguish between these relationships given only genetic data. The majority of grandparent-grandchild pairs can be differentiated from half sibling pairs on the basis of age (i.e. the age difference in most grandparent-grandchild relationships will be >30 years). However, it is much more difficult to resolve half sibling from avuncular pairs. Half siblings and avuncular pairs can be partially distinguished by the former’s longer haplotype sharing. Intuitively, this is because any chromosome segments that half siblings share have only gone through a total of two meioses since their common ancestor (i.e. transmission from the shared parent to half sibling one and transmission from the shared parent to half sibling two). In contrast, any shared haplotype segments have gone through a total of three meioses since the last common ancestor in the case of avuncular relationships (i.e. transmission from shared grandparent to uncle/aunt and transmission from shared grandparent to parent to child). However, classification is imperfect (Gusev et al. 2009; Hill and Weir 2011; Hill and White 2013), but could be improved further through the use of additional information including age difference of the pair and reported information on the parents (e.g. half siblings who share the same mother should produce consistent reports of maternal illnesses). Any half sibling pairs that are identified would need to be classified into maternal half siblings (who share a mother) and paternal half siblings (who share a father). Genetic data on the sex chromosomes and mitochondria could help facilitate this differentiation.

Finally, we note that there are several ways that our procedure could be improved/extended. First, we have only considered relative pairs in our derivations. Additional first degree relatives (i.e. additional siblings, the addition of one parent etc) would enable better genotype imputation and therefore increased power to detect parental genetic effects on offspring phenotypes. It is also possible that more distant relatives may also be informative for imputation, particularly if shared haplotypes could be identified within larger pedigree structures. Third, we have only considered one SNP at a time. It is possible that the inclusion of haplotype information may increase imputation fidelity. Fourth we note that it is likely that family dynamics will alter the strength of indirect parental genetic effects depending on the relationship of offspring to their parents. For example, the relationship between half siblings and their birth parents is likely to be qualitatively different to those of full siblings in nuclear families. Thus, for later-life phenotypes especially, parental genetic effect size estimates in half siblings may not be comparable to those estimated from full siblings. This may be perhaps less of an issue for maternal genetic effects on perinatal phenotypes. Finally, we note that the models that we have considered in this manuscript could be extended in a variety of ways including adding more relatives to help estimate sibling and/or parent of origin effects.

In conclusion, we have developed a suite of online genetic power calculators and software to assist researchers in detecting and partitioning loci that exhibit indirect parental genetic effects. We hope that our methods and utilities will form useful adjuncts to large ongoing genetic studies of parents and their offspring.

## Acknowledgements

We thank Peter Visscher, Loic Yengo and Matthew Keller for interesting discussions involving the challenging task of identifying half siblings from avuncular relative pairs using genetic data alone. This research was carried out at the Translational Research Institute, Woolloongabba, QLD 4102, Australia. The Translational Research Institute is supported by a grant from the Australian Government. J.L is supported by a University of Queensland and University of Exeter Accelerator grant (to D.M.E). G.H.M is supported by the Norwegian Research Council (Post doctorial mobility research grant 287198). D.M.E. is supported by an Australian National Health and Medical Research Council Senior Research Fellowship (1137714) and this work was supported by project grants (GNT1125200, GNT1183074, GNT1157714).

