## Supplementary figures and images for "Estimating indirect parental genetic effects on offspring phenotypes using virtual parental genotypes derived from sibling and half sibling pairs"

### Supplementary Figure 1

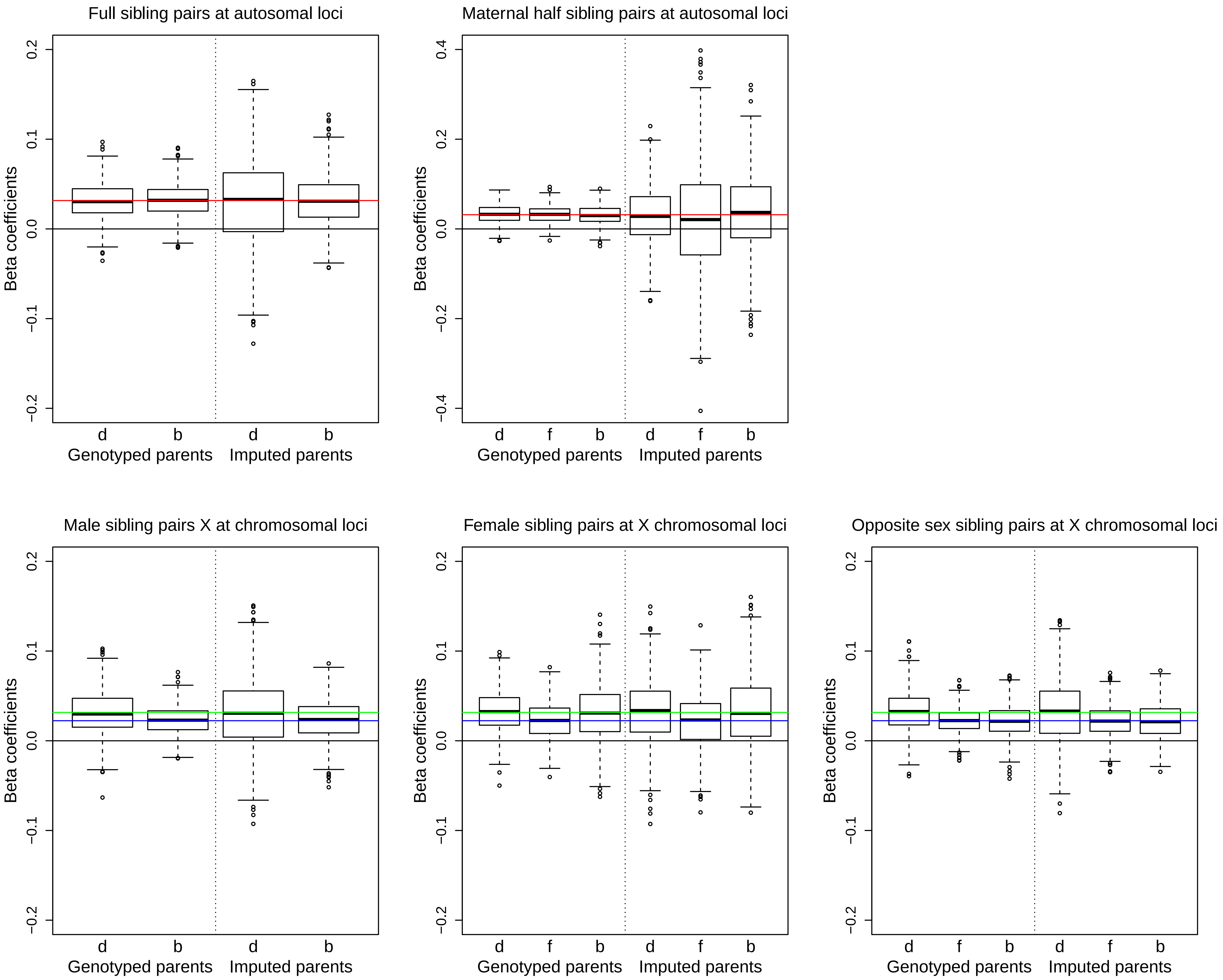
