## Supplementary Materials for "Estimating indirect parental genetic effects on offspring phenotypes using virtual parental genotypes derived from sibling and half sibling pairs"

Supplementary Methods	p.2

Derivation of non-centrality parameters	p. 2

Maternal half sibling pairs autosomes	p. 2

Siblings X chromosome loci	p. 9

Supplementary Table 1	p. 18

Supplementary Table 2	p. 19

Supplementary Table 3	p. 21

Supplementary Table 4	p. 22

Supplementary Table 15	p. 24

Legend to Supplementary Figure 1	p. 25

Simulation R Code	p. 26

**SUPPLEMENTARY METHODS**

**Derivation of non-centrality parameters**

We derive the expected minus two log-likelihoods for maternal half siblings (autosomes) and for male, female and opposite sex siblings (X chromosome).

**Maternal Half Siblings**

The model for maternal half sibling pair is

$$Y_{1}=bX_{1}+dX_{m}+fX_{f1}+\tau+\varepsilon_{1}$$

$$Y_{2}=bX_{2}+dX_{m}+fX_{f2}+\tau+\varepsilon_{2}$$

$$X_{1}=0.5(X_{m}+X_{f1})+\eta_{1}$$

$$X_{2}=0.5(X_{m}+X_{f2})+\eta_{2}$$

We assume random mating, so that Cov(X_m_,X_f_) = Cov(X_m_,X_f1_) = Cov(X_m_,X_f2_) = 0

So that the covariances between genotypes are

Cov(*X*_1_,*X*_2_) = 0.25

Cov(*X*_1_,*X*_m_) = Cov(*X*_2_,*X*_m_) = Cov(*X*_1_,*X*_f1_) = Cov(*X*_2_,*X*_f2_) = 0.5

Cov(*X*_1_,*X*_f2_) = Cov(*X*_2_,*X*_f1_) = 0

Cov(*X*_f1_,*X*_f2_) = 0

The covariances between phenotypes and genotypes are then

Cov(*Y*_1,_*X*_1_) = Cov(*Y*_2_,*X*_2_) = *b* + *0.5(d+f)*

Cov(*Y*_1_,*X*_2_) = Cov(*Y*_2_,*X*_1_) = 0.25*b +* 0.5*d*

Cov(*Y*_1_,*X*_m_) = Cov(*Y*_2_,*X*_m_) = 0.5*b* + *d*

Cov(*Y*_1_,*X*_f1_) = Cov(*Y*_2_,*X*_f2_) = 0.5*b + f*

Cov(*Y*_1_,*X*_f2_) = Cov(*Y*_2_,*X*_f1_) = 0

The covariance of the two phenotypes is

*Cov(Y_1_,Y_2_) = 0.25b^2^ + d^2^ + bd +* $\varphi^{2}$

The phenotypic variance can be decomposed as follows

$Var\left( Y \right)=\left( b^{2}+d^{2}+f^{2}+bd+bf \right)+\varphi^{2}+\sigma^{2}=1$

The variance of the true parental genotypes prior to standardization is equivalent to the expected heterozygosity, given the allele frequencies *p* and *q*

$$Var\left( X_{m} \right)=Var\left( X_{f} \right)=2pq=H$$

The variance of the imputed maternal and paternal genotypes are functions of the expected heterozygosity such that

$$Var\left( X_{m}^{'} \right)=\frac{5(H+1)}{(H+4)(2H+3)}$$

$$Var\left( X_{f}^{'} \right)=\frac{{(H+1)}^{2}}{(2H+3)(2H+1)}$$

The covariance between respective actual and imputed genotypes is equal to the variance of the imputed genotype:

$$Cov\left( X_{m},X_{m}^{'} \right)=Var(X_{m}^{'})$$

$$Cov\left( X_{f},X_{f}^{'} \right)=Var(X_{f}^{'})$$

The covariance between the two imputed paternal genotypes is a function of H:

$$Cov\left( X_{f1}^{'},X_{f2}^{'} \right)= \frac{-H}{8H+4}$$

as is the covariance between the imputed paternal and imputed maternal genotypes:

$$Cov\left( X_{m}^{'},X_{f1}^{'} \right)=Cov\left( X_{m}^{'},X_{f2}^{'} \right)= \frac{H+1}{4H+6}$$

The covariance between the imputed parental and sib genotypes and phenotypes are:

$$Cov\left( X_{m}^{'},X_{1} \right)=Cov\left( X_{m}^{'},X_{2} \right)=Cov\left( X_{f1}^{'},X_{1} \right)=Cov\left( X_{f2}^{'},X_{2} \right)=0.5$$

$$Cov\left( X_{m}^{'},Y_{1} \right)=Cov\left( X_{m}^{'},Y_{2} \right)=0.5b+dVar\left( X_{m}^{'} \right)+fCov\left( X_{m}^{'},X_{f}^{'} \right)$$

$$Cov\left( X_{f1}^{'},Y_{1} \right)=Cov\left( X_{f2}^{'},Y_{2} \right)=0.5b+dCov\left( X_{m}^{'},X_{f}^{'} \right)+fVar\left( X_{f}^{'} \right)$$

$$Cov\left( X_{f1}^{'},Y_{2} \right)=Cov\left( X_{f2}^{'},Y_{1} \right)=dCov\left( X_{m}^{'},X_{f}^{'} \right)+fCov\left( X_{f1}^{'},X_{f2}^{'} \right)$$

$$Cov\left( X_{f1}^{'},X_{2} \right)=Cov\left( X_{f2}^{'},X_{1} \right)=0$$

**Power calculation under linear mixed model analysis for autosomal loci for maternal half-siblings**

When actual maternal and paternal genotypes are available, the linear mixed model is

$$Y_{1}=bX_{1}+dX_{m}+fX_{f1}+\tau+\varepsilon_{1}$$

$$Y_{2}=bX_{2}+dX_{m}+fX_{f2}+\tau+\varepsilon_{2}$$

The fixed effects *b* and *d* are estimated by generalised least squares (GLS), where the covariance matrix of random effects is:

$$\Omega=\left( \begin{matrix} \sigma^{2}{+\varphi}^{2} & \varphi^{2} \\ \varphi^{2} & \sigma^{2}{+\varphi}^{2} \end{matrix} \right)$$

The inverse of the covariance matrix of random effects is

$$\Omega^{-1}=\frac{1}{\sigma^{2}(\sigma^{2}{+2\varphi}^{2})}\left( \begin{matrix} \sigma^{2}{+\varphi}^{2} & {-\varphi}^{2} \\ {-\varphi}^{2} & \sigma^{2}{+\varphi}^{2} \end{matrix} \right)$$

The asymptotic GLS estimates of a vector of parameters $\beta$ are given by

$$\hat{\beta}={E\left( X^{T}\Omega^{-1}X \right)}^{-1}E\left( X^{T}\Omega^{-1}Y \right)$$

We consider the following models:

Null model of no association

The residual covariance matrix is simply the covariance matrix of Y:

$$\Sigma=\Sigma_{Y}$$

$$=\left( \begin{matrix} 1 & 0.25b^{2} + d^{2} + bd + \varphi^{2} \\ 0.25b^{2} + d^{2} + bd + \varphi^{2} & 1 \end{matrix} \right)$$

The expected -2lnL of the model per sibpair is therefore

$$E(-2lnL)={ln(1-\left( 0.25b^{2} + d^{2} + bd + \varphi^{2} \right)}^{2})+2$$

Both parental genotypes in model (terms for $X_{m}$ and $X_{f}$ only):

The *X* matrix contains one column with elements $X_{m}$ and another with elements $X_{f1}$ and $X_{f2}$. The asymptotic GLS estimates of the regression coefficients of $X_{m}$ and $X_{f}$ are

$$\hat{g}=\left( \hat{d},\hat{f} \right)^{T}=E\left( X^{T}\Omega^{-1}X \right)^{-1}E\left( X^{T}\Omega^{-1}Y \right)$$

$$=E\left( \begin{matrix} \frac{2X_{m}^{2}}{\sigma^{2}{+2\varphi}^{2}} & \frac{X_{m}X_{f1}+X_{m}X_{f2}}{\sigma^{2}{+2\varphi}^{2}} \\ \frac{X_{m}X_{f1}+X_{m}X_{f2}}{\sigma^{2}{+2\varphi}^{2}} & \frac{{(X}_{f1}^{2}+X_{f2}^{2})\left( \sigma^{2}{+\varphi}^{2} \right)-2\varphi^{2}X_{f1}X_{f2}}{\sigma^{2}(\sigma^{2}{+2\varphi}^{2})} \end{matrix} \right)^{-1}E\left( \begin{matrix} \frac{X_{m}\left( Y_{1}+Y_{2} \right)}{\sigma^{2}{+2\varphi}^{2}} \\ \frac{{(\sigma}^{2}{+\varphi}^{2})\left( Y_{1}X_{f1}+Y_{2}X_{f2} \right)-\varphi^{2}\left( Y_{1}X_{f2}+Y_{2}X_{f1} \right)}{\sigma^{2}(\sigma^{2}{+2\varphi}^{2})} \end{matrix} \right)$$

$$=\left( \begin{matrix} \frac{Cov\left( X_{m},Y_{1} \right)+Cov\left( X_{m},Y_{2} \right)}{2} & \frac{Cov\left( X_{f1},Y_{1} \right)+Cov\left( X_{f2},Y_{2} \right)}{2} \end{matrix} \right)^{T}$$

$$=\left( \begin{matrix} 0.5b+d & 0.5b+f \end{matrix} \right)^{T}$$

The residual covariance matrix is

$$\Sigma=E\left( Y-X\hat{g} \right)\left( Y-X\hat{g} \right)^{T}$$

$$=\Sigma_{Y}-E(X\hat{g}Y^{T}+Y\hat{g}^{T}X^{T}-X\hat{g}\hat{g}^{T}X^{T})$$

where

$$E(X\hat{g}Y^{T})=E(Y\hat{g}^{T}X^{T})=\left( \begin{matrix} \hat{d}Cov\left( X_{m},Y_{1} \right)+\hat{f}Cov(X_{f1},Y_{1}) & \hat{d}Cov\left( X_{m},Y_{2} \right) \\ \hat{d}Cov\left( X_{m},Y_{1} \right) & \hat{d}Cov\left( X_{m},Y_{2} \right)+\hat{f}Cov(X_{f2},Y_{2}) \end{matrix} \right)$$

$$E\left( X\hat{g}\hat{g}^{T}X^{T} \right)=\left( \begin{matrix} \hat{d}^{2}+\hat{f}^{2} & \hat{d}^{2} \\ \hat{d}^{2} & \hat{d}^{2}+\hat{f}^{2} \end{matrix} \right)$$

Therefore

$$\Sigma=\left( \begin{matrix} 1-{(0.5b+d)}^{2} & \varphi^{2} \\ \varphi^{2} & 1-{(0.5b+d)}^{2} \end{matrix} \right)$$

The expected -2lnL of the model per half sibling pair is:

$$E(-2lnL)=\ln|\Sigma|+2$$

Offspring genotypes in model only (terms $X_{1}$ and $X_{2}$ only):

The *X* matrix contains one column with elements $X_{1}$ and $X_{2}$. The asymptotic GLS estimate of the regression coefficient of $X_{1}$ and $X_{2}$ is

$$\hat{b}=E\left( X^{T}\Omega^{-1}X \right)^{-1}E\left( X^{T}\Omega^{-1}Y \right)$$

$$=E\left( \left( \sigma^{2}+\varphi^{2} \right)\left( X_{1}^{2}+X_{2}^{2} \right)-2X_{1}X_{2}\varphi^{2} \right)^{-1}E\left( \left( \sigma^{2}+\varphi^{2} \right)\left( X_{1}Y_{1}+X_{2}Y_{2} \right)-\varphi^{2}\left( X_{1}Y_{2}+X_{2}Y_{1} \right) \right)$$

$$=\frac{\left( \sigma^{2}+\varphi^{2} \right)\left( Cov\left( X_{1},Y_{1} \right)+Cov\left( X_{2},Y_{2} \right) \right)-\varphi^{2}\left( Cov\left( X_{1},Y_{2} \right)+Cov\left( X_{2},Y_{1} \right) \right)}{\left( \sigma^{2}+\varphi^{2} \right)\left( Var\left( X_{1} \right)+Var\left( X_{2} \right) \right)-2\varphi^{2}Cov(X_{1},X_{2})}$$

$$=b+\frac{\left( {d\sigma}^{2}+f\sigma^{2}+f\varphi^{2} \right)}{\left( {2\sigma}^{2}+2\varphi^{2}-0.5\varphi^{2} \right)}$$

The residual covariance matrix is

$$\Sigma=E\left( Y-X\hat{b} \right)\left( Y-X\hat{b} \right)^{T}$$

$$=\Sigma_{Y}-\hat{b}\left( \Sigma_{YX}+\Sigma_{XY} \right)+\hat{b}^{2}\Sigma_{X}$$

Where

$$\Sigma_{YX}=\left( \begin{matrix} b + 0.5(d+f) & 0.25b + 0.5d \\ 0.25b + 0.5d & b + 0.5(d+f) \end{matrix} \right)$$

$$\Sigma_{XY}=\left( \begin{matrix} b + 0.5(d+f) & 0.25b + 0.5d \\ 0.25b + 0.5d & b + 0.5(d+f) \end{matrix} \right)$$

$$\Sigma_{X}=\left( \begin{matrix} 1 & 0.25 \\ 0.25 & 1 \end{matrix} \right)$$

Therefore

$$\Sigma=\left( \begin{matrix} 1-\left( 2b+d+f \right)\hat{b}+\hat{b}^{2} & 0.25b^{2} + d^{2} + bd + \varphi^{2}-\left( 0.5b+d \right)\hat{b}+0.25\hat{b}^{2} \\ 0.25b^{2} + d^{2} + bd + \varphi^{2}-\left( 0.5b+d \right)\hat{b}+0.25\hat{b}^{2} & 1-\left( 2b+d+f \right)\hat{b}+\hat{b}^{2} \end{matrix} \right)$$

The expected -2lnL of the model per half sibling pair is:

$$E(-2lnL)=ln|\Sigma|+2$$

Full Omnibus Model (terms for $X_{m}$, $X_{f}$, $X_{1}$ and $X_{2}$):

The *X* matrix contains three columns; column 1 with elements $X_{1}$ and $X_{2}$, column 2 with elements $X_{m}$ and $X_{m}$, and column 3 with $X_{f1}$ and $X_{f2}$. The asymptotic GLS estimate of the regression coefficients of columns 1 and 2 are

$$\hat{g}={(\hat{b},\hat{d},\hat{f})}^{T}=E\left( X^{T}\Omega^{-1}X \right)^{-1}E\left( X^{T}\Omega^{-1}Y \right)$$

$$={(b,d, f)}^{T}$$

The residual covariance matrix is

$$\Sigma=E\left( Y-X\hat{g} \right)\left( Y-X\hat{g} \right)^{T}$$

$$=\Omega$$

$$=\left( \begin{matrix} \sigma^{2}{+\varphi}^{2} & \varphi^{2} \\ \varphi^{2} & \sigma^{2}{+\varphi}^{2} \end{matrix} \right)$$

The expected -2lnL of the model per half sibling pair is:

$$E(-2lnL)=ln|\Sigma|+2$$

Imputed parental genotypes only in model (terms for $X_{m}^{'}$ and $X_{f}^{'}$):

When only imputed maternal genotypes are available, the linear mixed model becomes

$$Y_{1}=bX_{1}+dX_{m}^{'}+fX_{f1}^{'}+\tau+\varepsilon_{1}$$

$$Y_{2}=bX_{2}+dX_{m}^{'}+fX_{f2}^{'}+\tau+\varepsilon_{2}$$

The *X* matrix contains one column with elements $X_{m}^{'}$ and another with elements $X_{f1}^{'}$ and $X_{f2}^{'}$. The asymptotic GLS estimates of the regression coefficients of $X_{m}^{'}$ and $X_{f}^{'}$ are

$$\hat{g}=\left( \hat{d},\hat{f} \right)^{T}=E\left( X^{T}\Omega^{-1}X \right)^{-1}E\left( X^{T}\Omega^{-1}Y \right)$$

$$=E\left( \begin{matrix} \frac{2{X_{m}^{'}}^{2}}{\sigma^{2}{+2\varphi}^{2}} & \frac{X_{m}^{'}X_{f1}^{'}+X_{m}^{'}X_{f2}^{'}}{\sigma^{2}{+2\varphi}^{2}} \\ \frac{X_{m}^{'}X_{f1}^{'}+X_{m}^{'}X_{f2}^{'}}{\sigma^{2}{+2\varphi}^{2}} & \frac{({X_{f1}^{'}}^{2}+{X_{f2}^{'}}^{2})\left( \sigma^{2}{+\varphi}^{2} \right)-2\varphi^{2}X_{f1}^{'}X_{f2}^{'}}{\sigma^{2}(\sigma^{2}{+2\varphi}^{2})} \end{matrix} \right)^{-1}E\left( \begin{matrix} \frac{X_{m}^{'}\left( Y_{1}+Y_{2} \right)}{\sigma^{2}{+2\varphi}^{2}} \\ \frac{{(\sigma}^{2}{+\varphi}^{2})\left( Y_{1}X_{f1}^{'}+Y_{2}X_{f2}^{'} \right)-\varphi^{2}\left( Y_{1}X_{f2}^{'}+Y_{2}X_{f1}^{'} \right)}{\sigma^{2}(\sigma^{2}{+2\varphi}^{2})} \end{matrix} \right)$$

The residual covariance matrix is

$$\Sigma=E\left( Y-X\hat{g} \right)\left( Y-X\hat{g} \right)^{T}$$

$$=\Sigma_{Y}-E(X\hat{g}Y^{T}+Y\hat{g}^{T}X^{T}-X\hat{g}\hat{g}^{T}X^{T})$$

where

$$E(X\hat{g}Y^{T})=E(Y\hat{g}^{T}X^{T})=\left( \begin{matrix} \hat{d}Cov\left( X_{m}^{'},Y_{1} \right)+\hat{f}Cov(X_{f1}^{'},Y_{1}) & \hat{d}Cov\left( X_{m}^{'},Y_{2} \right)+\hat{f}Cov(X_{f1}^{'},Y_{2}) \\ \hat{d}Cov\left( X_{m}^{'},Y_{1} \right)+\hat{f}Cov(X_{f2}^{'},Y_{1}) & \hat{d}Cov\left( X_{m}^{'},Y_{2} \right)+\hat{f}Cov(X_{f2}^{'},Y_{2}) \end{matrix} \right)$$

$$E\left( X\hat{g}\hat{g}^{T}X^{T} \right)=\left( \begin{matrix} \hat{d}^{2}Var\left( X_{m}^{'} \right)+2\hat{d}\hat{f}Cov\left( X_{m}^{'},X_{f1}^{'} \right)+\hat{f}^{2}Var(X_{f1}^{'}) & \hat{d}^{2}Var\left( X_{m}^{'} \right)+2\hat{d}\hat{f}Cov\left( X_{m}^{'},X_{f1}^{'} \right)+\hat{f}^{2}Cov(X_{f1}^{'},X_{f2}^{'}) \\ \hat{d}^{2}Var\left( X_{m}^{'} \right)+2\hat{d}\hat{f}Cov\left( X_{m}^{'},X_{f2}^{'} \right)+\hat{f}^{2}Cov(X_{f1}^{'},X_{f2}^{'}) & \hat{d}^{2}Var\left( X_{m}^{'} \right)+2\hat{d}\hat{f}Cov\left( X_{m}^{'},X_{f2}^{'} \right)+\hat{f}^{2}Var(X_{f2}^{'}) \end{matrix} \right)$$

The expected -2lnL of the model per half sibling pair is:

$$E(-2lnL)=ln|\Sigma|+2$$

Full omnibus model with imputed parental genotypes (terms for $X_{m}^{'}$,$X_{f}^{'}$, $X_{1}$ and $X_{2}$):

The *X* matrix contains three columns; column 1 with elements $X_{1}$ and $X_{2}$, column 2 with elements $X_{m}^{'}$ and $X_{m}^{'}$, and column 3 with elements $X_{f1}^{'}$ and $X_{f2}^{'}$. The asymptotic GLS estimate of the regression coefficients of columns 1, 2, and 3 are

$${\hat{g}=(\hat{b},\hat{d}, \hat{f})}^{T}=E\left( X^{T}\Omega^{-1}X \right)^{-1}E\left( X^{T}\Omega^{-1}Y \right)$$

where

$$E\left( X^{T}\Omega^{-1}X \right)^{-1}=\left( \begin{matrix} \frac{\left( \sigma^{2}{+\varphi}^{2} \right)\left( Var\left( X_{1} \right)+Var\left( X_{2} \right) \right)-2\varphi^{2}\left( Cov(X_{1},X_{2}) \right)}{\sigma^{2}(\sigma^{2}{+2\varphi}^{2})} & \frac{Cov\left( X_{1},X_{m}^{'} \right)+Cov\left( X_{2},X_{m}^{'} \right)}{\sigma^{2}{+2\varphi}^{2}} & \frac{\left( \sigma^{2}{+\varphi}^{2} \right)\left( Cov(X_{f1}^{'},X_{1})+Cov(X_{f2}^{'},X_{2}) \right)-\varphi^{2}\left( Cov(X_{f1}^{'},X_{2})+Cov(X_{f2}^{'},X_{1}) \right)}{\sigma^{2}(\sigma^{2}{+2\varphi}^{2})} \\ \frac{Cov\left( X_{1},X_{m}^{'} \right)+Cov\left( X_{2},X_{m}^{'} \right)}{\sigma^{2}{+2\varphi}^{2}} & \frac{2Var(X_{m}^{'})}{\sigma^{2}{+2\varphi}^{2}} & \frac{2Cov(X_{m}^{'},X_{f}^{'})}{\sigma^{2}{+2\varphi}^{2}} \\ \frac{\left( \sigma^{2}{+\varphi}^{2} \right)\left( Cov(X_{f1}^{'},X_{1})+Cov(X_{f2}^{'},X_{2}) \right)-\varphi^{2}\left( Cov(X_{f1}^{'},X_{2})+Cov(X_{f2}^{'},X_{1}) \right)}{\sigma^{2}(\sigma^{2}{+2\varphi}^{2})} & \frac{2Cov(X_{m}^{'},X_{f}^{'})}{\sigma^{2}{+2\varphi}^{2}} & \frac{\left( \sigma^{2}{+\varphi}^{2} \right)\left( Var(X_{f1}^{'})+Var(X_{f2}^{'}) \right)-2\varphi^{2}Cov(X_{f1}^{'},X_{f2}^{'})}{\sigma^{2}(\sigma^{2}{+2\varphi}^{2})} \end{matrix} \right)^{-1}$$

$$E\left( X^{T}\Omega^{-1}Y \right)=\left( \begin{matrix} \frac{\left( \sigma^{2}{+\varphi}^{2} \right)\left( Cov\left( X_{1},Y_{1} \right)+Cov\left( X_{2},Y_{2} \right) \right)-\varphi^{2}\left( Cov\left( X_{1},Y_{2} \right)+Cov\left( X_{2},Y_{1} \right) \right)}{\sigma^{2}(\sigma^{2}{+2\varphi}^{2})} \\ \frac{Cov\left( Y_{1},X_{m}^{'} \right)+Cov\left( Y_{2},X_{m}^{'} \right)}{\sigma^{2}{+2\varphi}^{2}} \\ \frac{\left( \sigma^{2}{+\varphi}^{2} \right)\left( Cov(X_{f1}^{'},Y_{1})+Cov(X_{f2}^{'},Y_{2}) \right)-\varphi^{2}\left( Cov(X_{f1}^{'},Y_{2})+Cov(X_{f2}^{'},Y_{1}) \right)}{\sigma^{2}(\sigma^{2}{+2\varphi}^{2})} \end{matrix} \right)$$

The residual covariance matrix is

$$\Sigma=E\left( Y-X\hat{g} \right)\left( Y-X\hat{g} \right)^{T}$$

$$=\Sigma_{Y}-E(X\hat{g}Y^{T}+Y\hat{g}^{T}X^{T}-X\hat{g}\hat{g}^{T}X^{T})$$

where

$$E(X\hat{g}Y^{T})=E(Y\hat{g}^{T}X^{T})=\left( \begin{matrix} \hat{b}Cov\left( X_{1},Y_{1} \right)+\hat{d}Cov\left( X_{m}^{'},Y_{1} \right)+fCov(X_{f1}^{'},Y_{1}) & \hat{b}Cov\left( X_{1},Y_{2} \right)+\hat{d}Cov\left( X_{m}^{'},Y_{2} \right)+fCov(X_{f1}^{'},Y_{2}) \\ \hat{b}Cov\left( X_{2},Y_{1} \right)+\hat{d}Cov\left( X_{m}^{'},Y_{1} \right)+fCov(X_{f2}^{'},Y_{1}) & \hat{b}Cov\left( X_{2},Y_{2} \right)+\hat{d}Cov\left( X_{m}^{'},Y_{2} \right)+fCov(X_{f2}^{'},Y_{2}) \end{matrix} \right)$$

$$E\left( X\hat{g}\hat{g}^{T}X^{T} \right)=\left( \begin{matrix} \hat{b}^{2}+\hat{d}^{2}Var\left( X_{m}^{'} \right)+\hat{f}^{2}Var\left( X_{f}^{'} \right)+\hat{b}\hat{d}+\hat{b}\hat{f}+2\hat{d}\hat{f}Cov\left( X_{m}^{'},X_{f}^{'} \right) & 0.25\hat{b}^{2}+\hat{d}^{2}Var\left( X_{m}^{'} \right)+\hat{f}^{2}Cov\left( X_{f1}^{'},X_{f2}^{'} \right)+\hat{b}\hat{d}+2\hat{d}\hat{f}Cov\left( X_{m}^{'},X_{f}^{'} \right) \\ 0.25\hat{b}^{2}+\hat{d}^{2}Var\left( X_{m}^{'} \right)+\hat{f}^{2}Cov\left( X_{f1}^{'},X_{f2}^{'} \right)+\hat{b}\hat{d}+2\hat{d}\hat{f}Cov\left( X_{m}^{'},X_{f}^{'} \right) & \hat{b}^{2}+\hat{d}^{2}Var\left( X_{m}^{'} \right)+\hat{f}^{2}Var\left( X_{f}^{'} \right)+\hat{b}\hat{d}+\hat{b}\hat{f}+2\hat{d}\hat{f}Cov\left( X_{m}^{'},X_{f}^{'} \right) \end{matrix} \right)$$

The expected -2lnL of the model per half sibling pair is:

$$E(-2lnL)=ln|\Sigma|+2$$

**Sibling Pairs at X Chromosome loci**

We assume that unstandardized male genotypes are coded *G_1_* $\in$ {0,2} and female genotypes coded as *G_2_* $\in$ {0,1 or 2}. This means that males have double the variance of females at non-pseudosomal X-linked loci. We assume that the regression coefficient of offspring phenotype on offspring genotype is the same in males and females. We parameterize our model so that X chromosome genotypes are standardized relative to the female genotypic variance, such that female siblings and mothers have unit variance, while male siblings and fathers have a variance equal to two. We also assume that male and female phenotypes are standardized. As covariances between sib pair genotypes are sex-dependent, three separate models must be constructed for female-female, male-male, and female-male sib pairs.

The causal model for a sib-pair where sibling one is male and sibling two is female is then

$$Y_{1}=bX_{1}+dX_{m}+fX_{f}+\tau+\varepsilon_{1}$$

$$Y_{2}=bX_{2}+dX_{m}+fX_{f}+\tau+\varepsilon_{2}$$

$$X_{1}=X_{m}+\eta_{M}$$

$$X_{2}=0.5X_{m}+{0.5X}_{f}+\eta_{F}$$

where *τ* is a random effect shared by the siblings, *ε*_1_ and *ε*_2_ are uncorrelated error terms for the two phenotypes, and *η*_M_ and *η*_F_ are random effects due to segregation. The variances of the random effects are

$$Var\left( \varepsilon_{1} \right)=\sigma_{\varepsilon1}^{2}$$

$$Var\left( \varepsilon_{2} \right)=\sigma_{\varepsilon2}^{2}$$

$$Var\left( \tau\right)=\varphi^{2}$$

$$Var\left( \eta_{M} \right)=1$$

$$Var\left( \eta_{F} \right)=0.25$$

where $Var\left( \varepsilon_{1} \right)=Var\left( \varepsilon_{2} \right)=\sigma^{2}$ in the case of same sex siblings.

**Covariances**

We assume random mating, so that Cov(X_m_,X_f_) = 0

So that the covariances between genotypes are

Mother-Daughter

Cov(*X*_2_,*X*_m_) = 0.5

Mother-Son

Cov(*X*_1_,*X*_m_) = 1

Father-Daughter

Cov(*X*_2_,*X*_f_) = 1

Father-Son

Cov(*X*_1_,*X*_f_) = 0

Brother-Brother

Cov(*X*_1_,*X*_2_) = 1

Sister-Sister

Cov(*X*_1_,*X*_2_) = 0.75

Brother-Sister

Cov(*X*_1_,*X*_2_) = 0.5

The covariances between phenotypes and genotypes for the following sibling pair combinations are then:

Female-Female

Cov(*Y*_1,_*X*_1_) = Cov(*Y*_2_,*X*_2_) = $b+0.5d+f$

Cov(*Y*_1_,*X*_2_) = Cov(*Y*_2_,*X*_1_) = $0.75b+0.5d+f$

Cov(*Y*_1_,*X*_m_) = Cov(*Y*_2_,*X*_m_) =$0.5b+d$

Cov(*Y*_1_,*X*_f_) = Cov(*Y*_2_,*X*_f_) = $b+2f$

Male-Male

Cov(*Y*_1,_*X*_1_) = Cov(*Y*_2_,*X*_2_) = $2b+d$

Cov(*Y*_1_,*X*_2_) = Cov(*Y*_2_,*X*_1_) = $b+d$

Cov(*Y*_1_,*X*_m_) = Cov(*Y*_2_,*X*_m_) = $b+d$

Cov(*Y*_1_,*X*_f_) = Cov(*Y*_2_,*X*_f_) = $2f$

Male-Female

Cov(*Y*_1,_*X*_1_) = $2b+d$

Cov(*Y*_2_,*X*_2_) = $b+0.5d+f$

Cov(*Y*_1_,*X*_2_) = $0.5\left( b+d \right)+f$

Cov(*Y*_2_,*X*_1_) = $\left( 0.5b+d \right)$

Cov(*Y*_1_,*X*_m_) = $b+d$

Cov(*Y*_2_,*X*_m_) = $0.5b+d$

Cov(*Y*_1_,*X*_f_) = $2f$

Cov(*Y*_2_,*X*_f_) = $b+2f$

The covariance of the two phenotypes is

Female-Female

*Cov(Y_1_,Y_2_) =* $0.75b^{2}+d^{2}+{2f}^{2}+bd+2bf+\varphi^{2}$

Male-Male

*Cov(Y_1_,Y_2_) =* $b^{2}+d^{2}+{2f}^{2}+2bd+\varphi^{2}$

Male-Female

*Cov(Y_1_,Y_2_) =* $0.5b^{2}+d^{2}+{2f}^{2}+1.5bd+bf+\varphi^{2}$

The phenotypic variance of offspring can be decomposed as follows

Male

$$Var\left( Y \right)=\left( 2b^{2}+d^{2}+2f^{2}+2bd \right)+\varphi^{2}+\sigma_{\varepsilon1}^{2}=1$$

Female

$$Var\left( Y \right)=\left( b^{2}+d^{2}+{2f}^{2}+bd+2bf \right)+\varphi^{2}+\sigma_{\varepsilon2}^{2}=1$$

The variance of the imputed maternal genotype, $Var(X_{m}^{'})$, is a function of p or the expected female heterozygosity *H* such that for the following sib-pairs

Female-Female

$$Var\left( X_{m}^{'} \right)=\frac{2H^{2}+4H-7}{-3H-12}$$

Male-Male

$$Var\left( X_{m}^{'} \right)=\frac{3}{H+4}$$

Male-Female

$$Var\left( X_{m}^{'} \right)=\frac{2H^{2}+6H+7}{(2H+3)(H+4)}$$

The variance of the imputed paternal genotype, $Var(X_{f}^{'})$, is a function of the male heterozygote frequency $H_{f}$ $H_{f}= 4pq$

Female-Female

$$Var\left( X_{f}^{'} \right)=\frac{16-H_{f}}{12}$$

Male-Female

$$Var\left( X_{f}^{'} \right)=\frac{H_{f}+4}{(H_{f}+3)}$$

Similarly, the covariance between imputed maternal and paternal genotypes are a function of the expected female heterozygosity *H*:

Female-Female:

$$Cov\left( X_{m}^{'},X_{f}^{'} \right)=\frac{\sqrt{2}(1+H)}{6}$$

Male-Female:

$$Cov\left( X_{m}^{'},X_{f}^{'} \right)=\frac{\sqrt{2}}{2(3+2H)}$$

and the covariance between imputed maternal or paternal genotypes and a given child’s phenotype *Y* are

for female offspring:

$$Cov\left( X_{m}^{'},Y \right)=0.5b+dVar\left( X_{m}^{'} \right)+fCov\left( X_{m}^{'},X_{f}^{'} \right)$$

$$Cov\left( X_{f}^{'},Y \right)=b+dCov\left( X_{m}^{'},X_{f}^{'} \right)+fVar(X_{f}^{'})$$

for male offspring:

$$Cov\left( X_{m}^{'},Y \right)=b+dVar\left( X_{m}^{'} \right)+fCov\left( X_{m}^{'},X_{f}^{'} \right)$$

$$Cov\left( X_{f}^{'},Y \right)=dCov\left( X_{m}^{'},X_{f}^{'} \right)+fVar(X_{f}^{'})$$

**Power calculation under linear mixed model analysis for X chromosome loci**

When actual maternal and paternal genotypes are available, the linear mixed model is

$$Y_{1}=bX_{1}+dX_{m}+fX_{f}+\tau+\varepsilon_{1}$$

$$Y_{2}=bX_{2}+dX_{m}+fX_{f}+\tau+\varepsilon_{2}$$

The fixed effects *b*, *d,* and *f* are estimated by generalised least squares (GLS), where the covariance matrix of random effects is:

$$\Omega=\left( \begin{matrix} \sigma^{2}{+\varphi}^{2} & \varphi^{2} \\ \varphi^{2} & \sigma^{2}{+\varphi}^{2} \end{matrix} \right)$$

The inverse of the covariance matrix of random effects is

$$\Omega^{-1}=\frac{1}{\sigma^{2}(\sigma^{2}{+2\varphi}^{2})}\left( \begin{matrix} \sigma^{2}{+\varphi}^{2} & {-\varphi}^{2} \\ {-\varphi}^{2} & \sigma^{2}{+\varphi}^{2} \end{matrix} \right)$$

The asymptotic GLS estimates of a vector of parameters $\beta$ are given by

$$\hat{\beta}={E\left( X^{T}\Omega^{-1}X \right)}^{-1}E\left( X^{T}\Omega^{-1}Y \right)$$

We consider the following models for each possible sibling pair combination:

Null model of no association

The residual covariance matrix is simply the covariance matrix of Y:

Female-Female

$$\Sigma=\Sigma_{Y}=\left( \begin{matrix} 1 & 0.75b^{2}+d^{2}+{2f}^{2}+bd+2bf+\varphi^{2} \\ 0.75b^{2}+d^{2}+{2f}^{2}+bd+2bf+\varphi^{2} & 1 \end{matrix} \right)$$

Male-Male

$$\Sigma=\Sigma_{Y}=\left( \begin{matrix} 1 & b^{2}+d^{2}+{2f}^{2}+2bd+\varphi^{2} \\ b^{2}+d^{2}+{2f}^{2}+2bd+\varphi^{2} & 1 \end{matrix} \right)$$

Female-Male

$$\Sigma=\Sigma_{Y}=\left( \begin{matrix} 1 & 0.5b^{2}+d^{2}+{2f}^{2}+1.5bd+bf+\varphi^{2} \\ 0.5b^{2}+d^{2}+{2f}^{2}+1.5bd+bf+\varphi^{2} & 1 \end{matrix} \right)$$

The expected -2lnL of the model per sibling pair is therefore

Female-Female:

$$E(-2lnL)={ln(1-\left( 0.75b^{2}+d^{2}+{2f}^{2}+bd+2bf+\varphi^{2} \right)}^{2})+2$$

Male-Male:

$$E(-2lnL)=ln({1-\left( b^{2}+d^{2}+{2f}^{2}+2bd+\varphi^{2} \right)}^{2})+2$$

Female-Male:

$$E(-2lnL)={ln(1-\left( 0.5b^{2}+d^{2}+{2f}^{2}+1.5bd+bf+\varphi^{2} \right)}^{2})+2$$

Both parental genotypes only in model (terms for $X_{m}$ and $X_{f}$ only):

The *X* matrix contains two columns with elements $X_{m}$ and $X_{f}$. The asymptotic GLS estimates for the regression coefficients of $X_{m}$ and $X_{f}$ for a sibling pair are

$$\hat{g}=\left( \hat{d},\hat{f} \right)^{T}=E\left( X^{T}\Omega^{-1}X \right)^{-1}E\left( X^{T}\Omega^{-1}Y \right)$$

$$=E\left( \begin{matrix} \frac{2X_{m}^{2}}{\sigma^{2}{+2\varphi}^{2}} & \frac{2X_{m}X_{f}}{\sigma^{2}{+2\varphi}^{2}} \\ \frac{2X_{m}X_{f}}{\sigma^{2}{+2\varphi}^{2}} & \frac{2X_{f}^{2}}{\sigma^{2}{+2\varphi}^{2}} \end{matrix} \right)^{-1}E\left( \begin{matrix} \frac{X_{m}\left( Y_{1}+Y_{2} \right)}{\sigma^{2}{+2\varphi}^{2}} \\ \frac{X_{f}\left( Y_{1}+Y_{2} \right)}{\sigma^{2}{+2\varphi}^{2}} \end{matrix} \right)$$

$$=\left( \begin{matrix} \frac{Cov\left( X_{m},Y_{1} \right)+Cov\left( X_{m},Y_{2} \right)}{2} & \frac{Cov\left( X_{f},Y_{1} \right)+Cov\left( X_{f},Y_{2} \right)}{4} \end{matrix} \right)^{T}$$

Thus, the $\hat{g}$ matrix for each of the following sib pairs are

Female-Female

$$\hat{g}=\left( \begin{matrix} 0.5b+d & 0.5b+f \end{matrix} \right)^{T}$$

Male-Male

$$\hat{g}=\left( \begin{matrix} b+d & f \end{matrix} \right)^{T}$$

Female-Male

$$\hat{g}=\left( \begin{matrix} 0.5b+d & 0.25b+f \end{matrix} \right)^{T}$$

The residual covariance matrix is

$$\Sigma=E\left( Y-X\hat{g} \right)\left( Y-X\hat{g} \right)^{T}$$

$$=\Sigma_{Y}-E(X\hat{g}Y^{T}+Y\hat{g}^{T}X^{T}-X\hat{g}\hat{g}^{T}X^{T})$$

where

$$E(X\hat{g}Y^{T})=E(Y\hat{g}^{T}X^{T})=\left( \begin{matrix} \hat{d}Cov\left( X_{m},Y_{1} \right)+\hat{f}Cov(X_{f},Y_{1}) & \hat{d}Cov\left( X_{m},Y_{2} \right)+\hat{f}Cov(X_{f},Y_{2}) \\ \hat{d}Cov\left( X_{m},Y_{1} \right)+\hat{f}Cov(X_{f},Y_{1}) & \hat{d}Cov\left( X_{m},Y_{2} \right)+\hat{f}Cov(X_{f},Y_{2}) \end{matrix} \right)$$

$$E\left( X\hat{g}\hat{g}^{T}X^{T} \right)=\left( \begin{matrix} \hat{d}^{2}Var\left( X_{m} \right)+2\hat{d}\hat{f}Cov\left( X_{m},X_{f} \right)+\hat{f}^{2}Var(X_{f}) & \hat{d}^{2}Var\left( X_{m} \right)+2\hat{d}\hat{f}Cov\left( X_{m},X_{f} \right)+\hat{f}^{2}Var(X_{f}) \\ \hat{d}^{2}Var\left( X_{m} \right)+2\hat{d}\hat{f}Cov\left( X_{m},X_{f} \right)+\hat{f}^{2}Var(X_{f}) & \hat{d}^{2}Var\left( X_{m} \right)+2\hat{d}\hat{f}Cov\left( X_{m},X_{f} \right)+\hat{f}^{2}Var(X_{f}) \end{matrix} \right)$$

The expected -2lnL of the model per sibling pair is

$$E(-2lnL)=ln|\Sigma|+2$$

Offspring genotypes in model only (terms for $X_{1}$ and $X_{2}$ only):

The *X* matrix contains one column with elements $X_{1}$ and $X_{2}$. The asymptotic GLS estimate of the regression coefficient of $X_{1}$ and $X_{2}$ is

$$\hat{b}=E\left( X^{T}\Omega^{-1}X \right)^{-1}E\left( X^{T}\Omega^{-1}Y \right)$$

$$=E\left( \left( \sigma^{2}+\varphi^{2} \right)\left( X_{1}^{2}+X_{2}^{2} \right)-2X_{1}X_{2}\varphi^{2} \right)^{-1}E\left( \left( \sigma^{2}+\varphi^{2} \right)\left( X_{1}Y_{1}+X_{2}Y_{2} \right)-\varphi^{2}\left( X_{1}Y_{2}+X_{2}Y_{1} \right) \right)$$

$$=\frac{(\sigma^{2}+\varphi^{2})\left( Cov\left( X_{1},Y_{1} \right)+Cov\left( X_{2},Y_{2} \right) \right)-\varphi^{2}\left( Cov\left( X_{1},Y_{2} \right)+Cov\left( X_{2},Y_{1} \right) \right)}{\left( \sigma^{2}+\varphi^{2} \right)\left( Var\left( X_{1} \right)+Var\left( X_{2} \right) \right)-2\varphi^{2}\left( Cov\left( X_{1},X_{2} \right) \right)}$$

Thus, for each of the following sib-pair combinations,

Female-Female:

$$\hat{b}=b+\frac{\sigma^{2}\left( d+f \right)}{2\sigma^{2}+0.5\varphi^{2}}$$

Male-Male:

$$\hat{b}=b+\frac{\sigma^{2}d}{2\sigma^{2}+\varphi^{2}}$$

Female-Male:

$$\hat{b}=b+\frac{d\left( 2\sigma_{1}^{2}+\sigma_{2}^{2} \right)+2f\sigma_{2}^{2}}{2(2\sigma_{1}^{2}+\sigma_{2}^{2}+2\varphi^{2})}$$

The residual covariance matrix is

$$\Sigma=E\left( Y-X\hat{b} \right)\left( Y-X\hat{b} \right)^{T}$$

$$=\Sigma_{Y}-\hat{b}\left( \Sigma_{YX}+\Sigma_{XY} \right)+\hat{b}^{2}\Sigma_{X}$$

Where for the following sib-pair combinations,

Female-Female:

$$\Sigma_{YX}=\Sigma_{XY}=\left( \begin{matrix} b+0.5d+f & 0.75b+0.5d+f \\ 0.75b+0.5d+f & b+0.5d+f \end{matrix} \right)$$

$$\Sigma_{X}=\left( \begin{matrix} 1 & 0.75 \\ 0.75 & 1 \end{matrix} \right)$$

Male-Male:

$$\Sigma_{YX}=\Sigma_{XY}=\left( \begin{matrix} 2b+d & b+d \\ b+d & 2b+d \end{matrix} \right)$$

$$\Sigma_{X}=\left( \begin{matrix} 2 & 1 \\ 1 & 2 \end{matrix} \right)$$

Female-Male:

$$\Sigma_{YX}=\left( \begin{matrix} b+0.5d+f & 0.5b+d \\ 0.5b+0.5d+f & 2b+d \end{matrix} \right)$$

$$\Sigma_{XY}=\left( \begin{matrix} b+0.5d+f & 0.5\left( b+d \right)+f \\ \left( 0.5b+d \right) & 2b+d \end{matrix} \right)$$

$$\Sigma_{X}=\left( \begin{matrix} 1 & 0.5 \\ 0.5 & 2 \end{matrix} \right)$$

and $\Sigma_{Y}$ is defined as above for each sibling pair combination

Full Omnibus Model (terms for $X_{m}$, $X_{f}$, $X_{1}$ and $X_{2}$):

The *X* matrix contains three columns; column 1 with elements $X_{1}$ and $X_{2}$, column 2 with elements $X_{m}$ and $X_{m}$, and column 3 with elements $X_{f}$ and $X_{f}$. The asymptotic GLS estimate of the regression coefficients of columns 1, 2, and 3 are

$$\hat{g}={(\hat{b},\hat{d}, \hat{f})}^{T}=E\left( X^{T}\Omega^{-1}X \right)^{-1}E\left( X^{T}\Omega^{-1}Y \right)$$

$$={(b,d,f)}^{T}$$

The residual covariance matrix is

$$\Sigma=E\left( Y-X\hat{g} \right)\left( Y-X\hat{g} \right)^{T}$$

$$=\Omega$$

$$=\left( \begin{matrix} \sigma^{2}{+\varphi}^{2} & \varphi^{2} \\ \varphi^{2} & \sigma^{2}{+\varphi}^{2} \end{matrix} \right)$$

The expected -2lnL of the model per sibling pair is

$$E(-2lnL)=ln|\Sigma|+2$$

Imputed parental genotypes only in model (terms for $X_{m}^{'}$ and $X_{f}^{'}$only)

When only imputed maternal genotypes are available, the linear mixed model becomes

$$Y_{1}=bX_{1}+dX_{m}^{'}+fX_{f}^{'}+\tau+\varepsilon_{1}$$

$$Y_{2}=bX_{2}+dX_{m}^{'}+fX_{f}^{'}+\tau+\varepsilon_{2}$$

The *X* matrix contains two columns, one with elements $X_{m}^{'}$ and another with elements $X_{f}^{'}$. The asymptotic GLS estimate of the regression coefficient of $X_{m}^{'}$ and $X_{f}^{'}$ are

$$\hat{g}=\left( \hat{d},\hat{f} \right)^{T}=E\left( X^{T}\Omega^{-1}X \right)^{-1}E\left( X^{T}\Omega^{-1}Y \right)$$

$$=E\left( \begin{matrix} \frac{2{X_{m}^{'}}^{2}}{\sigma^{2}{+2\varphi}^{2}} & \frac{2X_{m}^{'}X_{f}^{'}}{\sigma^{2}{+2\varphi}^{2}} \\ \frac{2X_{m}'X_{f}'}{\sigma^{2}{+2\varphi}^{2}} & \frac{2{X_{f}^{'}}^{2}}{\sigma^{2}{+2\varphi}^{2}} \end{matrix} \right)^{-1}E\left( \begin{matrix} \frac{X_{m}^{'}\left( Y_{1}+Y_{2} \right)}{\sigma^{2}{+2\varphi}^{2}} \\ \frac{X_{f}^{'}\left( Y_{1}+Y_{2} \right)}{\sigma^{2}{+2\varphi}^{2}} \end{matrix} \right)$$

$$=\left( \begin{matrix} \frac{Var(X_{f}^{'})\left( Cov\left( X_{m}^{'},Y_{1} \right)+Cov\left( X_{m}^{'},Y_{2} \right) \right)-Cov(X_{m}^{'},X_{f}^{'})\left( Cov\left( X_{f}^{'},Y_{1} \right)+Cov\left( X_{f}^{'},Y_{2} \right) \right)}{2Var\left( X_{m}^{'} \right)2Var\left( X_{f}^{'} \right)-{2Cov(X_{m}^{'},X_{f}^{'})}^{2}} \\ \frac{Var(X_{m}^{'})\left( Cov\left( X_{f}^{'},Y_{1} \right)+Cov\left( X_{f}^{'},Y_{2} \right) \right)-Cov(X_{m}^{'},X_{f}^{'})\left( Cov\left( X_{m}^{'},Y_{1} \right)+Cov\left( X_{m}^{'},Y_{2} \right) \right)}{2Var\left( X_{m}^{'} \right)2Var\left( X_{f}^{'} \right)-{2Cov(X_{m}^{'},X_{f}^{'})}^{2}} \end{matrix} \right)$$

The residual covariance matrix is

$$\Sigma=E\left( Y-X\hat{g} \right)\left( Y-X\hat{g} \right)^{T}$$

$$=\Sigma_{Y}-E(X\hat{g}Y^{T}+Y\hat{g}^{T}X^{T}-X\hat{g}\hat{g}^{T}X^{T})$$

where

$$E(X\hat{g}Y^{T})=E(Y\hat{g}^{T}X^{T})=\left( \begin{matrix} \hat{d}Cov\left( X_{m}^{'},Y_{1} \right)+\hat{f}Cov(X_{f}^{'},Y_{1}) & \hat{d}Cov\left( X_{m}^{'},Y_{2} \right)+\hat{f}Cov(X_{f}^{'},Y_{2}) \\ \hat{d}Cov\left( X_{m}^{'},Y_{1} \right)+\hat{f}Cov(X_{f}^{'},Y_{1}) & \hat{d}Cov\left( X_{m}^{'},Y_{2} \right)+\hat{f}Cov(X_{f}^{'},Y_{2}) \end{matrix} \right)$$

$$E\left( X\hat{g}\hat{g}^{T}X^{T} \right)=\left( \begin{matrix} \hat{d}^{2}Var\left( X_{m}^{'} \right)+2\hat{d}\hat{f}Cov\left( X_{m}^{'},X_{f}^{'} \right)+\hat{f}^{2}Var(X_{f}^{'}) & \hat{d}^{2}Var\left( X_{m}^{'} \right)+2\hat{d}\hat{f}Cov\left( X_{m}^{'},X_{f}^{'} \right)+\hat{f}^{2}Var(X_{f}^{'}) \\ \hat{d}^{2}Var\left( X_{m}^{'} \right)+2\hat{d}\hat{f}Cov\left( X_{m}^{'},X_{f}^{'} \right)+\hat{f}^{2}Var(X_{f}^{'}) & \hat{d}^{2}Var\left( X_{m}^{'} \right)+2\hat{d}\hat{f}Cov\left( X_{m}^{'},X_{f}^{'} \right)+\hat{f}^{2}Var(X_{f}^{'}) \end{matrix} \right)$$

The expected -2lnL of the model per sibling pair is

$$E(-2lnL)=ln|\Sigma|+2$$

Full omnibus model with imputed parental genotypes (terms for $X_{m}^{'}$, $X_{f}^{'}$, $X_{1}$ and $X_{2}$):

The *X* matrix contains three columns; column 1 with elements $X_{1}$ and $X_{2}$, column 2 with elements $X_{m}^{'}$ and $X_{m}^{'}$, and column 3 with elements $X_{f}^{'}$ and $X_{f}^{'}$. The asymptotic GLS estimate of the regression coefficients of columns 1, 2, and 3 are

$${\hat{g}=(\hat{b},\hat{d}, \hat{f})}^{T}=E\left( X^{T}\Omega^{-1}X \right)^{-1}E\left( X^{T}\Omega^{-1}Y \right)$$

where

$$E\left( X^{T}\Omega^{-1}X \right)^{-1}=\left( \begin{matrix} \frac{\left( \sigma_{1}^{2}{+\varphi}^{2} \right)Var\left( X_{1} \right)\left( \sigma_{2}^{2}{+\varphi}^{2} \right)Var\left( X_{2} \right)-2\varphi^{2}\left( Cov(X_{1},X_{2}) \right)}{\sigma_{1}^{2}\sigma_{2}^{2}{+\varphi}^{2}(\sigma_{1}^{2}+\sigma_{2}^{2})} & \frac{\sigma_{1}^{2}Cov\left( X_{1},X_{m}^{'} \right)+\sigma_{2}^{2}Cov\left( X_{2},X_{m}^{'} \right)}{\sigma_{1}^{2}\sigma_{2}^{2}{+\varphi}^{2}(\sigma_{1}^{2}+\sigma_{2}^{2})} & \frac{\sigma_{1}^{2}Cov\left( X_{1},X_{f}^{'} \right)+\sigma_{2}^{2}Cov\left( X_{2},X_{f}^{'} \right)}{\sigma_{1}^{2}\sigma_{2}^{2}{+\varphi}^{2}(\sigma_{1}^{2}+\sigma_{2}^{2})} \\ \frac{\sigma_{1}^{2}Cov\left( X_{1},X_{m}^{'} \right)+\sigma_{2}^{2}Cov\left( X_{2},X_{m}^{'} \right)}{\sigma_{1}^{2}\sigma_{2}^{2}{+\varphi}^{2}(\sigma_{1}^{2}+\sigma_{2}^{2})} & \frac{(\sigma_{1}^{2}+\sigma_{2}^{2})Var(X_{m}^{'})}{\sigma_{1}^{2}\sigma_{2}^{2}{+\varphi}^{2}(\sigma_{1}^{2}+\sigma_{2}^{2})} & \frac{(\sigma_{1}^{2}+\sigma_{2}^{2})Cov(X_{m}^{'},X_{f}^{'})}{\sigma_{1}^{2}\sigma_{2}^{2}{+\varphi}^{2}(\sigma_{1}^{2}+\sigma_{2}^{2})} \\ \frac{\sigma_{1}^{2}Cov\left( X_{1},X_{f}^{'} \right)+\sigma_{2}^{2}Cov\left( X_{2},X_{f}^{'} \right)}{\sigma_{1}^{2}\sigma_{2}^{2}{+\varphi}^{2}(\sigma_{1}^{2}+\sigma_{2}^{2})} & \frac{(\sigma_{1}^{2}+\sigma_{2}^{2})Cov(X_{m}^{'},X_{f}^{'})}{\sigma_{1}^{2}\sigma_{2}^{2}{+\varphi}^{2}(\sigma_{1}^{2}+\sigma_{2}^{2})} & \frac{(\sigma_{1}^{2}+\sigma_{2}^{2})Var(X_{f}^{'})}{\sigma_{1}^{2}\sigma_{2}^{2}{+\varphi}^{2}(\sigma_{1}^{2}+\sigma_{2}^{2})} \end{matrix} \right)^{-1}$$

$$E\left( X^{T}\Omega^{-1}Y \right)=\left( \begin{matrix} \frac{\left( \sigma_{1}^{2}{+\varphi}^{2} \right)Cov\left( X_{1},Y_{1} \right)+\left( \sigma_{1}^{2}{+\varphi}^{2} \right)Cov\left( X_{2},Y_{2} \right)-\varphi^{2}\left( Cov\left( X_{1},Y_{2} \right)+Cov\left( X_{2},Y_{1} \right) \right)}{\sigma_{1}^{2}\sigma_{2}^{2}{+\varphi}^{2}(\sigma_{1}^{2}+\sigma_{2}^{2})} \\ \frac{\sigma_{1}^{2}Cov\left( Y_{1},X_{m}^{'} \right)+\sigma_{2}^{2}Cov\left( Y_{2},X_{m}^{'} \right)}{\sigma_{1}^{2}\sigma_{2}^{2}{+\varphi}^{2}(\sigma_{1}^{2}+\sigma_{2}^{2})} \\ \frac{\sigma_{1}^{2}Cov\left( Y_{1},X_{f}^{'} \right)+\sigma_{2}^{2}Cov\left( Y_{2},X_{f}^{'} \right)}{\sigma_{1}^{2}\sigma_{2}^{2}{+\varphi}^{2}(\sigma_{1}^{2}+\sigma_{2}^{2})} \end{matrix} \right)$$

The residual covariance matrix is

$$\Sigma=E\left( Y-X\hat{g} \right)\left( Y-X\hat{g} \right)^{T}$$

$$=\Sigma_{Y}-E(X\hat{g}Y^{T}+Y\hat{g}^{T}X^{T}-X\hat{g}\hat{g}^{T}X^{T})$$

where

$$E(X\hat{g}Y^{T})=E(Y\hat{g}^{T}X^{T})=\left( \begin{matrix} \hat{b}Cov\left( X_{1},Y_{1} \right)+\hat{d}Cov\left( X_{m}^{'},Y_{1} \right)+\hat{f}Cov(X_{f}^{'},Y_{1}) & \hat{b}Cov\left( X_{1},Y_{2} \right)+\hat{d}Cov\left( X_{m}^{'},Y_{2} \right)+\hat{f}Cov(X_{f}^{'},Y_{2}) \\ \hat{b}Cov\left( X_{2},Y_{1} \right)+\hat{d}Cov\left( X_{m}^{'},Y_{1} \right)+\hat{f}Cov(X_{f}^{'},Y_{1}) & \hat{b}Cov\left( X_{2},Y_{2} \right)+\hat{d}Cov\left( X_{m}^{'},Y_{2} \right)+\hat{f}Cov(X_{f}^{'},Y_{2}) \end{matrix} \right)$$

$$E\left( X\hat{g}\hat{g}^{T}X^{T} \right)=\left( \begin{matrix} \hat{b}^{2}Var\left( X_{1} \right){+\hat{d}}^{2}Var\left( X_{m}^{'} \right)+\hat{f}^{2}Var\left( X_{f}^{'} \right)+2\hat{b}\hat{d}Cov\left( X_{m}^{'},X_{1} \right)+2\hat{b}\hat{f}Cov\left( X_{f}^{'},X_{1} \right)+2\hat{d}\hat{f}Cov\left( X_{f}^{'},X_{m}^{'} \right) & \hat{b}^{2}Cov\left( X_{1},X_{2} \right){+\hat{d}}^{2}Var\left( X_{m}^{'} \right)+\hat{f}^{2}Var\left( X_{f}^{'} \right)+2\hat{b}\hat{d}Cov\left( X_{m}^{'},X_{1} \right)+2\hat{b}\hat{f}Cov\left( X_{f}^{'},X_{1} \right)+2\hat{d}\hat{f}Cov\left( X_{f}^{'},X_{m}^{'} \right) \\ \hat{b}^{2}Cov\left( X_{1},X_{2} \right){+\hat{d}}^{2}Var\left( X_{m}^{'} \right)+\hat{f}^{2}Var\left( X_{f}^{'} \right)+2\hat{b}\hat{d}Cov\left( X_{m}^{'},X_{2} \right)+2\hat{b}\hat{f}Cov\left( X_{f}^{'},X_{2} \right)+2\hat{d}\hat{f}Cov\left( X_{f}^{'},X_{m}^{'} \right) & \hat{b}^{2}Var\left( X_{2} \right){+\hat{d}}^{2}Var\left( X_{m}^{'} \right)+\hat{f}^{2}Var\left( X_{f}^{'} \right)+2\hat{b}\hat{d}Cov\left( X_{m}^{'},X_{2} \right)+2\hat{b}\hat{f}Cov\left( X_{f}^{'},X_{2} \right)+2\hat{d}\hat{f}Cov\left( X_{f}^{'},X_{m}^{'} \right) \end{matrix} \right)$$

The expected -2lnL of the model per sibling pair is

$$E(-2lnL)=ln|\Sigma|+2$$

**Supplementary Table** **1** Probabilities (P) and expected dosages for imputed parental genotypes conditional on observed sibling pair genotypes at autosomal loci. The symbol *p* denotes the frequency of the trait decreasing allele *A*. The expected parental dosage refers to the expected number of trait increasing alleles *a*.

|  | **Parental Genotype Probabilities Conditional on Observed Sibling Genotypes** | | |  |
| --- | --- | --- | --- | --- |
|  | **P(*AA*)** | **P(*Aa*)** | **P(*aa*)** | **Expected Parental**  **Dosage** |
| **Sibling Genotypes** |  |  |  |  |
| ***AA*, *AA*** | $\frac{2p}{p+1}$ | $\frac{1-p}{p+1}$ | 0 | $\frac{1-p}{p+1}$ |
| ***AA*, *Aa*** | $\frac{p}{p+1}$ | $\frac{1}{p+1}$ | 0 | $\frac{1}{p+1}$ |
| ***AA*, *aa*** | 0 | 1 | 0 | 1 |
| ***Aa*, *Aa*** | $\frac{0.5\left( p-2 \right)p}{p^{2}-p-1}$ | $\frac{0.5}{1+\left( 1-p \right)p}$ | $\frac{0.5\left( p-1 \right)(p+1)}{p^{2}-p-1}$ | $\frac{p^{2}-1.5}{p^{2}-p-1}$ |
| ***Aa*, *aa*** | 0 | $\frac{1}{2-p}$ | $\frac{1-p}{2-p}$ | $\frac{3-2p}{2-p}$ |
| ***aa*, *aa*** | 0 | $\frac{p}{2-p}$ | $\frac{2-2p}{2-p}$ | $\frac{4-3p}{2-p}$ |

**Supplementary Table** **2** Probabilities (P) and expected dosages for imputed parental genotypes conditional on observed sibling pair genotypes at non-pseudosomal X chromosome loci. The symbol *p* denotes the frequency of the trait decreasing allele *A*. The expected parental dosage refers to the expected number of trait increasing alleles *a*. For opposite sex pairs, the genotype of the male sibling is given first. NA= not applicable due to the genotype combination being impossible.

|  | **Maternal Genotype** | | | | **Paternal Genotype** | | |
| --- | --- | --- | --- | --- | --- | --- | --- |
| **Sibling Genotypes** |  |  |  |  |  |  |  |
| **Male Pairs** | **P(*AA*)** | **P(*Aa*)** | **P(*aa*)** | **Expected Dosage** | **P(*A*)** | **P(*a*)** | **Expected Dosage** |
| ***A*, *A*** | $\frac{2p}{p+1}$ | $\frac{1-p}{p+1}$ | 0 | $\frac{1-p}{p+1}$ | $p$ | $1-p$ | $2(1-p$) |
| ***A*, *a*** | 0 | 1 | 0 | 1 | $p$ | $1-p$ | $2(1-p$) |
| ***a*, *a*** | 0 | $\frac{p}{2-p}$ | $\frac{2-2p}{2-p}$ | $\frac{4-3p}{2-p}$ | $p$ | $1-p$ | $2(1-p$) |
| **Female Pairs** | **P(*AA*)** | **P(*Aa*)** | **P(*aa*)** | **Expected Dosage** | **P(*A*)** | **P(*a*)** | **Expected Dosage** |
| ***AA*, *AA*** | $\frac{2p}{p+1}$ | $\frac{1-p}{p+1}$ | 0 | $\frac{1-p}{p+1}$ | 1 | 0 | 0 |
| ***AA*, *Aa*** | 0 | 1 | 0 | 1 | 1 | 0 | 0 |
| ***AA*, *aa*** | NA | NA | NA | NA | NA | NA | NA |
| ***Aa*, *Aa*** | $\frac{2}{3}p$ | $\frac{1}{3}$ | $\frac{2}{3}(1-p)$ | $\frac{5}{3}-\frac{4}{3}p$ | $\frac{2}{3}-\frac{1}{3}p$ | $\frac{1}{3}(1+p)$ | $\frac{2}{3}(1+p)$ |
| ***Aa*, *aa*** | 0 | 1 | 0 | 1 | 0 | 1 | 2 |
| ***aa*, *aa*** | 0 | $\frac{p}{2-p}$ | $\frac{2-2p}{2-p}$ | $\frac{4-3p}{2-p}$ | 0 | 1 | 2 |
| **Opposite Sex Pairs** | **P(*AA*)** | **P(*Aa*)** | **P(*aa*)** | **Expected Dosage** | **P(*A*)** | **P(*a*)** | **Expected Dosage** |
| ***A*, *AA*** | $\frac{2p}{p+1}$ | $\frac{1-p}{p+1}$ | 0 | $\frac{1-p}{p+1}$ | 1 | 0 | 0 |
| ***A*, *Aa*** | $\frac{2p}{2p+1}$ | $\frac{1}{2p+1}$ | 0 | $\frac{1}{2p+1}$ | $\frac{p}{2p+1}$ | $\frac{p+1}{2p+1}$ | $\frac{2p+2}{2p+1}$ |
| ***A*, *aa*** | 0 | 1 | 0 | 1 | 0 | 1 | 2 |
| ***a*, *AA*** | 0 | 1 | 0 | 1 | 1 | 0 | 0 |
| ***a*, *Aa*** | 0 | $\frac{1}{3-2p}$ | $\frac{2-2p}{3-2p}$ | $\frac{5-4p}{3-2p}$ | $\frac{2-p}{3-2p}$ | $\frac{1-p}{3-2p}$ | $\frac{2-2p}{3-2p}$ |
| ***a*, *aa*** | 0 | $\frac{p}{2-p}$ | $\frac{2-2p}{2-p}$ | $\frac{4-3p}{2-p}$ | 0 | 1 | 2 |

**Supplementary Table 3** Genotype probabilities (P) and expected dosages for imputed genotypes of shared parent conditional on observed half sibling pair genotypes at autosomal loci. The symbol *p* denotes the frequency of the trait decreasing allele *A*. The expected parental dosage refers to the expected number of trait increasing alleles *a*.

|  | **Shared Parental Genotype** | | |  |
| --- | --- | --- | --- | --- |
| **Half Sibling**  **Genotype** | **P(*AA*)** | **P(*Aa*)** | **P(*aa*)** | **Expected Dosage** |
| ***AA*, *AA*** | $\frac{2p}{p+1}$ | $\frac{1-p}{p+1}$ | $0$ | $\frac{1-p}{p+1}$ |
| ***AA*, *Aa*** | $\frac{2p}{2p+1}$ | $\frac{1}{2p+1}$ | $0$ | $\frac{1}{2p+1}$ |
| ***AA*, *aa*** | $0$ | $1$ | $0$ | 1 |
| ***Aa*, *Aa*** | $\frac{2p(1-p)}{-4p^{2}+4p+1}$ | $\frac{1}{-4p^{2}+4p+1}$ | $\frac{2p(1-p)}{-4p^{2}+4p+1}$ | 1 |
| ***Aa*, *aa*** | $0$ | $\frac{1}{3-2p}$ | $\frac{2-2p}{3-2p}$ | $\frac{5-4p}{3-2p}$ |
| ***aa*, *aa*** | $0$ | $\frac{p}{2-p}$ | $\frac{2(1-p)}{2-p}$ | $\frac{4-3p}{2-p}$ |
| ***Aa*, *AA*** | $\frac{2p}{2p+1}$ | $\frac{1}{2p+1}$ | $0$ | $\frac{1}{2p+1}$ |
| ***aa*, *AA*** | $0$ | $1$ | $0$ | 1 |
| ***aa*, *Aa*** | $0$ | $\frac{1}{3-2p}$ | $\frac{2-2p}{3-2p}$ | $\frac{5-4p}{3-2p}$ |

**Supplementary Table** **4** Genotype probabilities (P) and expected dosages of imputed genotypes for the father of the first maternal half sibling and the father of the second maternal half sibling conditional on observed maternal half sibling genotypes at autosomal loci. The symbol *p* denotes the frequency of the trait decreasing allele *A*. The expected parental dosage refers to the expected number of trait increasing alleles *a*. In the case of paternal half siblings, the probabilities and expected dosages are the same for the mother of the first half sibling and the mother of the second half sibling.

|  | **Father of First Half Sibling** | | | | **Father of Second Half Sibling** | | | |
| --- | --- | --- | --- | --- | --- | --- | --- | --- |
|  | **P(*AA*)** | **P(*Aa*)** | **P(*aa*)** | **Expected Dosage** | **P(*AA*)** | **P(*Aa*)** | **P(*aa*)** | **Expected Dosage** |
| ***AA*, *AA*** | $p$ | $1-p$ | $0$ | $1-p$ | $p$ | $1-p$ | 0 | $1-p$ |
| ***AA*, *Aa*** | $p$ | $1-p$ | $0$ | $1-p$ | $\frac{p^{2}}{1+2p}$ | $\frac{2p}{1+2p}$ | $\frac{{1-p}^{2}}{1+2p}$ | $\frac{{-2p}^{2}+2p+2}{1+2p}$ |
| ***AA*, *aa*** | $p$ | $1-p$ | $0$ | $1-p$ | 0 | $p$ | $1-p$ | $2-p$ |
| ***Aa*, *Aa*** | $\frac{3p^{2}-2p^{3}}{-4p^{2}+4p+1}$ | $\frac{-4p^{2}+4p}{-4p^{2}+4p+1}$ | $\frac{2p^{3}-3p^{2}+1}{-4p^{2}+4p+1}$ | $\frac{4p^{3}-10p^{2}+4p+2}{-4p^{2}+4p+1}$ | $\frac{3p^{2}-2p^{3}}{-4p^{2}+4p+1}$ | $\frac{-4p^{2}+4p}{-4p^{2}+4p+1}$ | $\frac{2p^{3}-3p^{2}+1}{-4p^{2}+4p+1}$ | $\frac{4p^{3}-10p^{2}+4p+2}{-4p^{2}+4p+1}$ |
| ***Aa*, *aa*** | $\frac{p(2-p)}{3-2p}$ | $\frac{2-2p}{3-2p}$ | $\frac{{(1-p)}^{2}}{3-2p}$ | $\frac{4-6p+{2p}^{2}}{3-2p}$ | 0 | $p$ | $1-p$ | $2-p$ |
| ***aa*, *aa*** | $0$ | $p$ | $1-p$ | $2-p$ | 0 | $p$ | $1-p$ | $2-p$ |
| ***Aa*, *AA*** | $\frac{p^{2}}{2p+1}$ | $\frac{2p}{2p+1}$ | $\frac{1-p^{2}}{2p+1}$ | $\frac{{-2p}^{2}+2p+2}{1+2p}$ | $p$ | $1-p$ | 0 | $1-p$ |
| ***aa*, *AA*** | $0$ | $p$ | $1-p$ | $2-p$ | $p$ | $1-p$ | 0 | $1-p$ |
| ***aa*, *Aa*** | $0$ | $p$ | $1-p$ | $2-p$ | $\frac{2p-p^{2}}{3-2p}$ | $\frac{2-2p}{3-2p}$ | $\frac{{(1-p)}^{2}}{3-2p}$ | $\frac{4-6p+{2p}^{2}}{3-2p}$ |

**Supplementary Table 15.** Computational performance of IMPISH. Reported runtimes of IMPISH for imputation and performing association analyses. All tests were performed in the same computing environment with 256 GB memory and 16 CPU cores with solid-state disk in one computer node.

| Number of SNPs | Sample size  (Sibling pairs) | Imputation (hours) | Association analysis (hours) |
| --- | --- | --- | --- |
| 500,000 | 1,000 | 1.03 | 0.54 |
| 500,000 | 5,000 | 2.61 | 3.32 |
| 500,000 | 10,000 | 6.20 | 6.11 |
| 500,000 | 20,000 | 12.27 | 8.42 |

**Supplementary Figure 1.** Comparison of maternal (d), paternal (f) and fetal (b) genetic effects estimated using genotyped parental genotypes versus imputed parental genotypes in simulations. We simulated the size of genetic effects accounting for 0.1% of the variance in the offspring trait (*b^2^* = *d^2^* = *f^2^* = 0.1%), frequency of the trait decreasing allele of *p* = 0.1, and shared residual variance at $\varphi^{2}$ = 0.2. For all simulations we used N = 2000 sibling pairs/half sibling pairs, and 1000 replications. Red lines indicate the expected beta coefficients for parental (d) and fetal effects (f) in full/half sibling pairs at autosomal loci. Blue lines indicate the expected beta coefficients for paternal (f) and male fetal (b) effects at X chromosomal loci. Green lines indicate the expected beta coefficients for maternal (d) and female fetal (b) effect at X chromosomal loci. For opposite sex sibling pairs at X chromosomal loci, we simulated the fetal effect (b) to be the same for both siblings assuming using male genotypes. The full omnibus model was simulated and fitted in all simulations. R codes implementing the simulations are provided in the Supplementary Materials.

**R codes for simulations for exploring bias, power and type 1 error of genetic association**

**Full sibling pairs at autosomal loci**

library(lmerTest)

library(MASS)

rm(list=ls())

Nrep = 1000	#Number of simulation replicates

N = 2000	#Sample size

Vm = 0.001	#Variance explained by maternal effect

Vf = 0.001	#Variance explained by fetal effect

p = 0.1	#Increaser allele frequency

Opp = FALSE	#Boolean variable to denote whether maternal and fetal effects in opposite directions (TRUE = opposite directions)

rho = 0.2	#Covariance between sibling residual variances

q <- 1-p	#Decreaser allele frequency

#Create vectors to save results from regression analyses

#For analysis using simulated parental and fetal genotypes

beta_genotyped_mat <- vector(length = Nrep)	#Fitting regression with maternal genotypes known (maternal coefficient)

beta_genotyped_fet <- vector(length = Nrep)	#Fitting regression with maternal genotypes known (fetal coefficient)

se_genotyped_mat <- vector(length = Nrep)

se_genotyped_fet <- vector(length = Nrep)

pval_genotyped_mat <- vector(length = Nrep)

pval_genotyped_fet <- vector(length = Nrep)

#Analysis using imputed parental genotypes and simulated fetal genotypes

beta_imputed_mat <- vector(length = Nrep)	#Fitting regression with genotypes imputed (maternal coefficient)

beta_imputed_fet <- vector(length = Nrep)	#Fitting regression with genotypes imputed (fetal coefficient)

se_imputed_mat <- vector(length = Nrep)

se_imputed_fet <- vector(length = Nrep)

pval_imputed_mat <- vector(length = Nrep)

pval_imputed_fet <- vector(length = Nrep)

#Analysis using only sib genotypes

beta_sib <- vector(length = Nrep)	#Fitting incorrect model to just sibs

se_sib <- vector(length = Nrep)

pval_sib <- vector(length = Nrep)

#ANOVA for full model against null model

pval_imputed_omni <- vector(length = Nrep)

pval_genotyped_omni <- vector(length = Nrep)

a <- sqrt(1/(2*p*q))	#Create genetic variable of variance one. Assume no dominance.

BetaM <- sqrt(Vm)	#Path coefficient for maternal effect

BetaF <- sqrt(Vf)	#Path coefficient for fetal effect

if(Opp == TRUE) {	#If Opp is true then maternal and fetal effects have opposite directions of effect

BetaF = -BetaF

}

Ve <- (1 - BetaM^2 - BetaF^2 - 2*0.5*BetaM*BetaF) #Residual variance in trait

for(j in 1:Nrep) {

#Sample mothers' genotypes

Zm <- sample(x = c("AA","Aa","aa"), size = N, replace = TRUE, prob = c(p^2, 2*p*q, q^2))

#Sample fathers' genotypes

Zp <- sample(x = c("AA","Aa","aa"), size = N, replace = TRUE, prob = c(p^2, 2*p*q, q^2))

Z_sib1 <- vector(length = N)

Z_sib2 <- vector(length = N)

Z_imp <- vector(length = N)	#Imputed vector of genotypes at mother's locus

#Simulate sib 1 genotype

r <- runif(N)

for (i in 1:N) {

Z_sib1[i] <- switch(paste(Zm[i],Zp[i]),

'AA AA'='AA',

'AA Aa'=ifelse(r[i] <= 0.5, 'AA', 'Aa'),

'AA aa'='Aa',

'Aa AA'=ifelse(r[i] <= 0.5, 'AA', 'Aa'),

'Aa Aa'=ifelse(r[i] <= 0.25, 'aa', ifelse(r[i] > 0.75, 'AA', 'Aa')),

'Aa aa'=ifelse(r[i] <= 0.5, 'aa', 'Aa'),

'aa AA'='Aa',

'aa Aa'=ifelse(r[i] <= 0.5, 'aa', 'Aa'),

'aa aa'='aa')

}

#Simulate sib 2 genotype

r <- runif(N)

for (i in 1:N) {

Z_sib2[i] <- switch(paste(Zm[i], Zp[i]),

'AA AA'='AA',

'AA Aa'=ifelse(r[i] <= 0.5, 'AA', 'Aa'),

'AA aa'='Aa',

'Aa AA'=ifelse(r[i] <= 0.5, 'AA', 'Aa'),

'Aa Aa'=ifelse(r[i] <= 0.25, 'aa', ifelse(r[i] > 0.75, 'AA', 'Aa')),

'Aa aa'=ifelse(r[i] <= 0.5, 'aa', 'Aa'),

'aa AA'='Aa',

'aa Aa'=ifelse(r[i] <= 0.5, 'aa', 'Aa'),

'aa aa'='aa')

}

#Change coding to 0, 1 and 2 in order to calculate allele frequencies

Z_sib1_012 <- ifelse(Z_sib1=='AA', 0, ifelse(Z_sib1=='Aa', 1, 2))

Z_sib2_012 <- ifelse(Z_sib2=='AA', 0, ifelse(Z_sib2=='Aa', 1, 2))

#Calculate observed allele frequency based on sibling genotypes

q_obs <- sum(Z_sib1_012+Z_sib2_012)/(2*N*2)

p_obs <- 1 - q_obs

#Impute mother's genotypes

#Calculate conditional probability of mother's genotypes given sibs genotypes

p_AA_AA_AA = p_obs*(2*p_obs+2)/(p_obs+1)^2	#P(Mum = AA | Sib = AA and Sib = AA)

p_Aa_AA_AA = 2/(p_obs+1) - 1	#P(Mum = Aa | Sib = AA and Sib = AA)

p_aa_AA_AA = 0	#P(Mum = aa | Sib = AA and Sib = AA)

p_AA_AA_Aa = p_obs/(p_obs+1)	#P(Mum = AA | Sib = AA and Sib = Aa)

p_Aa_AA_Aa = 1/(p_obs+1)	#P(Mum = Aa | Sib = AA and Sib = Aa)

p_aa_AA_Aa = 0	#P(Mum = aa | Sib = AA and Sib = Aa)

p_AA_AA_aa = 0	#P(Mum = AA | Sib = AA and Sib = aa)

p_Aa_AA_aa = 1	#P(Mum = Aa | Sib = AA and Sib = aa)

p_aa_AA_aa = 0	#P(Mum = aa | Sib = AA and Sib = aa)

p_AA_Aa_Aa = (0.5*(p_obs-2)*p_obs)/(p_obs^2-p_obs-1)	#P(Mum = AA | Sib = Aa and Sib = Aa)

p_Aa_Aa_Aa = -0.5 / ((p_obs-1)*p_obs-1)	#P(Mum = Aa | Sib = Aa and Sib = Aa)

p_aa_Aa_Aa = (0.5*(p_obs-1)*(p_obs+1))/(p_obs^2-p_obs-1)	#P(Mum = aa | Sib = Aa and Sib = Aa)

p_AA_aa_Aa = 0	#P(Mum = AA | Sib = aa and Sib = Aa)

p_Aa_aa_Aa = 1/(q_obs+1)	#P(Mum = Aa | Sib = aa and Sib = Aa)

p_aa_aa_Aa = q_obs/(q_obs+1)	#P(Mum = aa | Sib = aa and Sib = Aa)

p_AA_aa_aa = 0	#P(Mum = AA | Sib = aa and Sib = aa)

p_Aa_aa_aa = 2/(q_obs+1) - 1	#P(Mum = Aa | Sib = aa and Sib = aa)

p_aa_aa_aa = q_obs*(2*q_obs+2)/(q_obs+1)^2	#P(Mum = aa | Sib = aa and Sib = aa)

for (i in 1:N) {

Z_imp[i] <- switch(paste(Z_sib1[i], Z_sib2[i]),

'AA AA'= -a*p_AA_AA_AA + 0*p_Aa_AA_AA + a*p_aa_AA_AA,

'AA Aa'= -a*p_AA_AA_Aa + 0*p_Aa_AA_Aa + a*p_aa_AA_Aa,

'AA aa'= -a*p_AA_AA_aa + 0*p_Aa_AA_aa + a*p_aa_AA_aa,

'Aa AA'= -a*p_AA_AA_Aa + 0*p_Aa_AA_Aa + a*p_aa_AA_Aa,

'Aa Aa'= -a*p_AA_Aa_Aa + 0*p_Aa_Aa_Aa + a*p_aa_Aa_Aa,

'Aa aa'= -a*p_AA_aa_Aa + 0*p_Aa_aa_Aa + a*p_aa_aa_Aa,

'aa AA'= -a*p_AA_AA_aa + 0*p_Aa_AA_aa + a*p_aa_AA_aa,

'aa Aa'= -a*p_AA_aa_Aa + 0*p_Aa_aa_Aa + a*p_aa_aa_Aa,

'aa aa'= -a*p_AA_aa_aa + 0*p_Aa_aa_aa + a*p_aa_aa_aa)

}

#Change coding AA/Aa/aa to the genetic value -a/0/a

Z_sib1 <- ifelse(Z_sib1=='AA', -a, ifelse(Z_sib1=='Aa', 0, a))

Z_sib2 <- ifelse(Z_sib2=='AA', -a, ifelse(Z_sib2=='Aa', 0, a))

Zm <- ifelse(Zm=='AA', -a, ifelse(Zm=='Aa', 0, a))

Zp <- ifelse(Zp=='AA', -a, ifelse(Zp=='Aa', 0, a))

#Create correlated error variables for sib 1 and sib 2

Sigma <- matrix(c(Ve , rho, rho, Ve),2,2)

e <- mvrnorm(n = N, mu = c(0, 0), Sigma)

#Simulate offspring outcome

Y_sib1 <- BetaM*Zm + BetaF*Z_sib1 + e[,1]

Y_sib2 <- BetaM*Zm + BetaF*Z_sib2 + e[,2]

#Create a test dataset of simulated and imputed genotypes and phenotype (offspring outcome)

test <- data.frame(rbind(cbind(1:N, Y_sib1, Zm, Z_imp, Z_sib1),

cbind(1:N, Y_sib2, Zm, Z_imp, Z_sib2)))

colnames(test) <- c("fam", "Y","Zm", "Z_imp", "Z")

#Run analyses

lmer_genotyped <- lmer(Y ~ Zm + Z + (1|fam), data = test, REML = FALSE); results_genotyped <- summary(lmer_genotyped)

lmer_imputed <- lmer(Y ~ Z_imp + Z + (1|fam), data = test, REML = FALSE); results_imputed <- summary(lmer_imputed)

lmer_sib <- lmer(Y ~ Z + (1|fam), data = test, REML = FALSE); results_sib <- summary(lmer_sib)

lmer_null <- lmer(Y ~ (1|fam), data = test, REML = FALSE)

beta_genotyped_mat[j] <- results_genotyped$coefficient[2,1]

se_genotyped_mat[j] <- results_genotyped$coefficient[2,2]

pval_genotyped_mat[j] <- results_genotyped$coefficient[2,5]

beta_genotyped_fet[j] <- results_genotyped$coefficient[3,1]

se_genotyped_fet[j] <- results_genotyped$coefficient[3,2]

pval_genotyped_fet[j] <- results_genotyped$coefficient[3,5]

pval_genotyped_omni[j] <- anova(lmer_genotyped,lmer_null)[2,8]

beta_imputed_mat[j] <- results_imputed$coefficient[2,1]

se_imputed_mat[j] <- results_imputed$coefficient[2,2]

pval_imputed_mat[j] <- results_imputed$coefficient[2,5]

beta_imputed_fet[j] <- results_imputed$coefficient[3,1]

se_imputed_fet[j] <- results_imputed$coefficient[3,2]

pval_imputed_fet[j] <- results_imputed$coefficient[3,5]

pval_imputed_omni[j] <- anova(lmer_imputed,lmer_null)[2,8]

beta_sib[j] <- results_sib$coefficient[2,1]

se_sib[j] <- results_sib$coefficient[2,2]

pval_sib[j] <- results_sib$coefficient[2,5]

}

**Half sibling pairs at autosomal loci**

library(lmerTest)

library(MASS)

rm(list=(ls()))

Nrep = 1000	#Number of simulation replicates

N = 2000	#Sample size

Vm = 0.001	#Variance explained by maternal effect

Vp = 0.001	#Variance explained by paternal effect

Vf = 0.001	#Variance explained by fetal effect

p = 0.1	#Increaser allele frequency

Opp = FALSE	#Boolean variable to denote whether maternal and fetal effects in opposite directions (TRUE = opposite directions)

rho = 0.2	#Covariance between half sibling residual variances

q <- 1-p	#Decreaser allele frequency

#Create vectors to save results from regression analyses

#For analysis using simulated genotypes

beta_genotyped_mat <- vector(length = Nrep) #Fitting regression with maternal and paternal genotypes known (maternal coefficient)

beta_genotyped_pat <- vector(length = Nrep) #Fitting regression with maternal and paternal genotypes known (paternal coefficient)

beta_genotyped_fet <- vector(length = Nrep) #Fitting regression with maternal and paternal genotypes known (fetal coefficient)

se_genotyped_mat <- vector(length = Nrep)

se_genotyped_pat <- vector(length = Nrep)

se_genotyped_fet <- vector(length = Nrep)

pval_genotyped_mat <- vector(length = Nrep)

pval_genotyped_pat <- vector(length = Nrep)

pval_genotyped_fet <- vector(length = Nrep)

#Analysis using imputed maternal and paternal genotypes

beta_imputed_mat <- vector(length = Nrep)	#Fitting regression with genotypes imputed (maternal coefficient)

beta_imputed_pat <- vector(length = Nrep)	#Fitting regression with genotypes imputed (paternal coefficient)

beta_imputed_fet <- vector(length = Nrep)	#Fitting regression with genotypes imputed (fetal coefficient)

se_imputed_mat <- vector(length = Nrep)

se_imputed_pat <- vector(length = Nrep)

se_imputed_fet <- vector(length = Nrep)

pval_imputed_mat <- vector(length = Nrep)

pval_imputed_pat <- vector(length = Nrep)

pval_imputed_fet <- vector(length = Nrep)

#Analysis using only half sibling genotypes

beta_fet <- vector(length = Nrep)	#Fitting incorrect model including only half siblings

se_fet <- vector(length = Nrep)

pval_fet <- vector(length = Nrep)

#ANOVA for full model against null model

pval_genotyped_omni <- vector(length = Nrep)

pval_imputed_omni <- vector(length = Nrep)

a <- sqrt(1/2*p*q)	#Create genetic variable of one. Assume no dominance.

BetaM <- sqrt(Vm)	#Path coefficient for maternal effect

BetaP <- sqrt(Vp)	#Path coefficient for paternal effect

BetaF <- sqrt(Vf)	#Path coefficient for fetal effect

if(Opp == TRUE) {	#If Opp is true then maternal and fetal effects have opposite directions of effect

BetaF = -BetaF

}

Ve <- (1 - BetaM^2 - BetaP^2 - BetaF^2 - 2*0.5*BetaM*BetaF - 2*0.5*BetaP*BetaF) #Residual variance in trait for half siblings

for(j in 1:Nrep) {

#Sample mothers' genotypes

Zm <- sample(x = c('AA','Aa','aa'), size = N, replace = TRUE, prob = c(p^2, 2*p*q, q^2))

#Sample fathers' genotypes

Zp1 <- sample(x = c('AA','Aa','aa'), size = N, replace = TRUE, prob = c(p^2, 2*p*q, q^2))

Zp2 <- sample(x = c('AA','Aa','aa'), size = N, replace = TRUE, prob = c(p^2, 2*p*q, q^2))

Z_hsib1 <- vector(length = N)

Z_hsib2 <- vector(length = N)

Zm_imp <- vector(length = N)	#Imputed vector of genotypes at mum's locus

Zp1_imp <- vector(length = N)	#Imputed vector of genotypes at dad1's locus

Zp2_imp <- vector(length = N)	#Imputed vector of genotypes at dad2's locus

#Simulate half sibling 1 genotype

r <- runif(N)

for (i in 1:N) {

Z_hsib1[i] <- switch(paste(Zm[i],Zp1[i]),

'AA AA'='AA',

'AA Aa'=ifelse(r[i] <= 0.5, 'AA', 'Aa'),

'AA aa'='Aa',

'Aa AA'=ifelse(r[i] <= 0.5, 'AA', 'Aa'),

'Aa Aa'=ifelse(r[i] <= 0.25, 'aa', ifelse(r[i] > 0.75, 'AA', 'Aa')),

'Aa aa'=ifelse(r[i] <= 0.5, 'aa', 'Aa'),

'aa AA'='Aa',

'aa Aa'=ifelse(r[i] <= 0.5, 'aa', 'Aa'),

'aa aa'='aa')

}

#Simulate half sibling 2 genotype

r <- runif(N)

for (i in 1:N) {

Z_hsib2[i] <- switch(paste(Zm[i],Zp2[i]),

'AA AA'='AA',

'AA Aa'=ifelse(r[i] <= 0.5, 'AA', 'Aa'),

'AA aa'='Aa',

'Aa AA'=ifelse(r[i] <= 0.5, 'AA', 'Aa'),

'Aa Aa'=ifelse(r[i] <= 0.25, 'aa', ifelse(r[i] > 0.75, 'AA', 'Aa')),

'Aa aa'=ifelse(r[i] <= 0.5, 'aa', 'Aa'),

'aa AA'='Aa',

'aa Aa'=ifelse(r[i] <= 0.5, 'aa', 'Aa'),

'aa aa'='aa')

}

#Change coding to 0, 1 and 2 in order to calculate allele frequencies

Z_hsib1_012 <- ifelse(Z_hsib1=='AA', 0, ifelse(Z_hsib1=='Aa', 1, 2))

Z_hsib2_012 <- ifelse(Z_hsib2=='AA', 0, ifelse(Z_hsib2=='Aa', 1, 2))

#Calculate observed allele frequency based on half sibling genotypes

q_obs <- sum(Z_hsib1_012, Z_hsib2_012)/(2*N*2)

p_obs <- 1 - q_obs

#Impute mother's genotypes

#Calculate conditional probability of mother's genotypes given half sibs genotypes

p_AA_AA_AA = (2*p_obs)/(p_obs+1)	#P(Mum = AA | HSib = AA and HSib = AA)

p_Aa_AA_AA = (1-p_obs)/(p_obs+1)	#P(Mum = Aa | HSib = AA and HSib = AA)

p_aa_AA_AA = 0	#P(Mum = aa | HSib = AA and HSib = AA)

p_AA_AA_Aa = (2*p_obs)/(2*p_obs+1)	#P(Mum = AA | HSib = AA and HSib = Aa)

p_Aa_AA_Aa = 1/(2*p_obs+1)	#P(Mum = Aa | HSib = AA and HSib = Aa)

p_aa_AA_Aa = 0	#P(Mum = aa | HSib = AA and HSib = Aa)

p_AA_AA_aa = 0	#P(Mum = AA | HSib = AA and HSib = aa)

p_Aa_AA_aa = 1	#P(Mum = Aa | HSib = AA and HSib = aa)

p_aa_AA_aa = 0	#P(Mum = aa | HSib = AA and HSib = aa)

p_AA_Aa_Aa = ((2*p_obs)*(1-p_obs))/((4*p_obs)*(1-p_obs)+1)	#P(Mum = AA | HSib = Aa and HSib = Aa)

p_Aa_Aa_Aa = (1-p_obs)/((p_obs-1)*(4*p_obs^2-4*p_obs-1))	#P(Mum = Aa | HSib = Aa and HSib = Aa)

p_aa_Aa_Aa = ((2*q_obs)*(1-q_obs))/((4*q_obs)*(1-q_obs)+1)	#P(Mum = aa | HSib = Aa and HSib = Aa)

p_AA_aa_Aa = 0	#P(Mum = AA | HSib = aa and HSib = Aa)

p_Aa_aa_Aa = 1/(2*q_obs+1)	#P(Mum = Aa | HSib = aa and HSib = Aa)

p_aa_aa_Aa = (2*q_obs)/(2*q_obs+1)	#P(Mum = aa | HSib = aa and HSib = Aa)

p_AA_aa_aa = 0	#P(Mum = AA | HSib = aa and HSib = aa)

p_Aa_aa_aa = (1-q_obs)/(q_obs+1)	#P(Mum = Aa | HSib = aa and HSib = aa)

p_aa_aa_aa = (2*q_obs)/(q_obs+1)	#P(Mum = aa | HSib = aa and HSib = aa)

for (i in 1:N) {

Zm_imp[i] <- switch(paste(Z_hsib1[i],Z_hsib2[i]),

'AA AA'= -a*p_AA_AA_AA + 0*p_Aa_AA_AA + a*p_aa_AA_AA,

'AA Aa'= -a*p_AA_AA_Aa + 0*p_Aa_AA_Aa + a*p_aa_AA_Aa,

'AA aa'= -a*p_AA_AA_aa + 0*p_Aa_AA_aa + a*p_aa_AA_aa,

'Aa AA'= -a*p_AA_AA_Aa + 0*p_Aa_AA_Aa + a*p_aa_AA_Aa,

'Aa Aa'= -a*p_AA_Aa_Aa + 0*p_Aa_Aa_Aa + a*p_aa_Aa_Aa,

'Aa aa'= -a*p_AA_aa_Aa + 0*p_Aa_aa_Aa + a*p_aa_aa_Aa,

'aa AA'= -a*p_AA_AA_aa + 0*p_Aa_AA_aa + a*p_aa_AA_aa,

'aa Aa'= -a*p_AA_aa_Aa + 0*p_Aa_aa_Aa + a*p_aa_aa_Aa,

'aa aa'= -a*p_AA_aa_aa + 0*p_Aa_aa_aa + a*p_aa_aa_aa)

}

#Impute father's genotypes

#Calculate conditional probability of father's genotypes given half sibling genotype

p_dAA_AA_AA <- p_obs	#P(Dad = AA | HSib = AA and HSib = AA)

p_dAa_AA_AA <- q_obs	#P(Dad = Aa | HSib = AA and HSib = AA)

p_daa_AA_AA <- 0	#P(Dad = aa | HSib = AA and HSib = AA)

p_dAA_AA_Aa <- p_obs	#P(Dad = AA | HSib = AA and HSib = Aa)

p_dAa_AA_Aa <- q_obs	#P(Dad = Aa | HSib = AA and HSib = Aa)

p_daa_AA_Aa <- 0	#P(Dad = aa | HSib = AA and HSib = Aa)

p_dAA_AA_aa <- p_obs	#P(Dad = AA | HSib = AA and HSib = aa)

p_dAa_AA_aa <- q_obs	#P(Dad = Aa | HSib = AA and HSib = aa)

p_daa_AA_aa <- 0	#P(Dad = aa | HSib = AA and HSib = aa)

p_dAA_Aa_AA <- p_obs^2/(2*p_obs+1)	#P(Dad = AA | HSib = Aa and HSib = AA)

p_dAa_Aa_AA <- 2*p_obs/(2*p_obs+1)	#P(Dad = Aa | HSib = Aa and HSib = AA)

p_daa_Aa_AA <- (1-p_obs^2)/(2*p_obs+1)	#P(Dad = aa | HSib = Aa and HSib = AA)

p_dAA_dAA_Aa_Aa <- (2*p_obs^3-p_obs^4)/(1+4*p_obs-4*p_obs^2)	#P(Dad1 = AA and Dad2 = AA | HSib1 = Aa and HSib2 = Aa)

p_dAA_dAa_Aa_Aa <- (2*p_obs^2-2*p_obs^3)/(1+4*p_obs-4*p_obs^2)	#P(Dad1 = AA and Dad2 = Aa | HSib1 = Aa and HSib2 = Aa)

p_dAA_daa_Aa_Aa <- (p_obs^2*q_obs^2)/(1+4*p_obs-4*p_obs^2)	#P(Dad1 = AA and Dad2 = aa | HSib1 = Aa and HSib2 = Aa)

p_dAa_dAA_Aa_Aa <- p_dAA_dAa_Aa_Aa	#P(Dad1 = Aa and Dad2 = AA | HSib1 = Aa and HSib2 = Aa)

p_dAa_dAa_Aa_Aa <- (2*p_obs*q_obs)/(1+4*p_obs-4*p_obs^2) #P(Dad1 = Aa and Dad2 = Aa | HSib1 = Aa and HSib2 = Aa)

p_dAa_daa_Aa_Aa <- (2*q_obs^2-2*q_obs^3)/(1+4*q_obs-4*q_obs^2)	#P(Dad1 = Aa and Dad2 = aa | HSib1 = Aa and HSib2 = Aa)

p_daa_dAA_Aa_Aa <- p_dAA_daa_Aa_Aa	#P(Dad1 = aa and Dad2 = AA | HSib1 = Aa and HSib2 = Aa)

p_daa_dAa_Aa_Aa <- p_dAa_daa_Aa_Aa	#P(Dad1 = aa and Dad2 = Aa | HSib1 = Aa and HSib2 = Aa)

p_daa_daa_Aa_Aa <- (2*q_obs^3-q_obs^4)/(1+4*q_obs-4*q_obs^2) #P(Dad1 = aa and Dad2 = aa | HSib1 = Aa and HSib2 = Aa)

p_dAA_Aa_aa <- p_obs*(2-p_obs)/(3-2*p_obs)	#P(Dad = AA | HSib = Aa and HSib = aa)

p_dAa_Aa_aa <- (2-2*p_obs)/(3-2*p_obs)	#P(Dad = Aa | HSib = Aa and HSib = aa)

p_daa_Aa_aa <- (1-p_obs)^2/(3-2*p_obs)	#P(Dad = aa | HSib = Aa and HSib = aa)

p_dAA_aa_AA <- 0	#P(Dad = AA | HSib = aa and HSib = AA)

p_dAa_aa_AA <- p_obs	#P(Dad = Aa | HSib = aa and HSib = AA)

p_daa_aa_AA <- q_obs	#P(Dad = aa | HSib = aa and HSib = AA)

p_dAA_aa_Aa <- 0	#P(Dad = AA | HSib = aa and HSib = Aa)

p_dAa_aa_Aa <- p_obs	#P(Dad = Aa | HSib = aa and HSib = Aa)

p_daa_aa_Aa <- q_obs	#P(Dad = aa | HSib = aa and HSib = Aa)

p_dAA_aa_aa <- 0	#P(Dad = AA | HSib = aa and HSib = aa)

p_dAa_aa_aa <- p_obs	#P(Dad = Aa | HSib = aa and HSib = aa)

p_daa_aa_aa <- q_obs	#P(Dad = aa | HSib = aa and HSib = aa)

for (i in 1:N) {

Zp1_imp[i] <- switch(paste(Z_hsib1[i],Z_hsib2[i]),

'AA AA'= -a*p_dAA_AA_AA + 0*p_dAa_AA_AA + a*p_daa_AA_AA,

'AA Aa'= -a*p_dAA_AA_Aa + 0*p_dAa_AA_Aa + a*p_daa_AA_Aa,

'AA aa'= -a*p_dAA_AA_aa + 0*p_dAa_AA_aa + a*p_daa_AA_aa,

'Aa AA'= -a*p_dAA_Aa_AA + 0*p_dAa_Aa_AA + a*p_daa_Aa_AA,

'Aa Aa'= -a*(p_dAA_dAA_Aa_Aa + p_dAA_dAa_Aa_Aa + p_dAA_daa_Aa_Aa) +

0*(p_dAa_dAA_Aa_Aa + p_dAa_dAa_Aa_Aa + p_dAa_daa_Aa_Aa) +

a*(p_daa_dAA_Aa_Aa + p_daa_dAa_Aa_Aa + p_daa_daa_Aa_Aa),

'Aa aa'= -a*p_dAA_Aa_aa + 0*p_dAa_Aa_aa + a*p_daa_Aa_aa,

'aa AA'= -a*p_dAA_aa_AA + 0*p_dAa_aa_AA + a*p_daa_aa_AA,

'aa Aa'= -a*p_dAA_aa_Aa + 0*p_dAa_aa_Aa + a*p_daa_aa_Aa,

'aa aa'= -a*p_dAA_aa_aa + 0*p_dAa_aa_aa + a*p_daa_aa_aa)

Zp2_imp[i] <- switch(paste(Z_hsib2[i],Z_hsib1[i]),

'AA AA'= -a*p_dAA_AA_AA + 0*p_dAa_AA_AA + a*p_daa_AA_AA,

'AA Aa'= -a*p_dAA_AA_Aa + 0*p_dAa_AA_Aa + a*p_daa_AA_Aa,

'AA aa'= -a*p_dAA_AA_aa + 0*p_dAa_AA_aa + a*p_daa_AA_aa,

'Aa AA'= -a*p_dAA_Aa_AA + 0*p_dAa_Aa_AA + a*p_daa_Aa_AA,

'Aa Aa'= -a*(p_dAA_dAA_Aa_Aa + p_dAA_dAa_Aa_Aa + p_dAA_daa_Aa_Aa) +

0*(p_dAa_dAA_Aa_Aa + p_dAa_dAa_Aa_Aa + p_dAa_daa_Aa_Aa) +

a*(p_daa_dAA_Aa_Aa + p_daa_dAa_Aa_Aa + p_daa_daa_Aa_Aa),

'Aa aa'= -a*p_dAA_Aa_aa + 0*p_dAa_Aa_aa + a*p_daa_Aa_aa,

'aa AA'= -a*p_dAA_aa_AA + 0*p_dAa_aa_AA + a*p_daa_aa_AA,

'aa Aa'= -a*p_dAA_aa_Aa + 0*p_dAa_aa_Aa + a*p_daa_aa_Aa,

'aa aa'= -a*p_dAA_aa_aa + 0*p_dAa_aa_aa + a*p_daa_aa_aa)

}

#Convert AA/Aa/aa to genetic value -a/0/a

Zm <- ifelse(Zm=='AA', -a, ifelse(Zm=='Aa', 0, a))

Zp1 <- ifelse(Zp1=='AA', -a, ifelse(Zp1=='Aa', 0, a))

Zp2 <- ifelse(Zp2=='AA', -a, ifelse(Zp2=='Aa', 0, a))

Z_hsib1 <- ifelse(Z_hsib1=='AA', -a, ifelse(Z_hsib1=='Aa', 0, a))

Z_hsib2 <- ifelse(Z_hsib2=='AA', -a, ifelse(Z_hsib2=='Aa', 0, a))

#Create correlated error variables for sib 1 and sib 2 in the model with both parents

Sigma <- matrix(c(Ve, rho, rho, Ve),2,2)

e <- mvrnorm(n = N, mu = c(0, 0), Sigma)

#Simulate offspring outcome

Y_hsib1 <- BetaM*Zm + BetaP*Zp1 + BetaF*Z_hsib1 + e[,1]

Y_hsib2 <- BetaM*Zm + BetaP*Zp2 + BetaF*Z_hsib2 + e[,2]

test <- data.frame(rbind(cbind(1:N, Y_hsib1, Zm, Zp1, Zm_imp, Zp1_imp, Z_hsib1),

cbind(1:N, Y_hsib2, Zm, Zp2, Zm_imp, Zp2_imp, Z_hsib2)))

colnames(test) <- c("fam", "Y","Zm", "Zp", "Zm_imp", "Zp_imp", "Z")

#Run analyses

lmer_genotyped <- lmer(Y ~ Zm + Zp + Z + (1|fam), data = test, REML = FALSE); results_genotyped <- summary(lmer_genotyped)

lmer_imputed <- lmer(Y ~ Zm_imp + Zp_imp + Z + (1|fam), data = test, REML = FALSE); results_imputed <- summary(lmer_imputed)

lmer_hsib <- lmer(Y ~ Z + (1|fam), data = test, REML = FALSE); results_hsib <- summary(lmer_hsib)

lmer_null <- lmer(Y ~ (1|fam), data = test, REML = FALSE)

beta_genotyped_mat[j] <- results_genotyped$coefficient[2,1]

se_genotyped_mat[j] <- results_genotyped$coefficient[2,2]

pval_genotyped_mat[j] <- results_genotyped$coefficient[2,5]

beta_genotyped_pat[j] <- results_genotyped$coefficient[3,1]

se_genotyped_pat[j] <- results_genotyped$coefficient[3,2]

pval_genotyped_pat[j] <- results_genotyped$coefficient[3,5]

beta_genotyped_fet[j] <- results_genotyped$coefficient[4,1]

se_genotyped_fet[j] <- results_genotyped$coefficient[4,2]

pval_genotyped_fet[j] <- results_genotyped$coefficient[4,5]

pval_genotyped_omni[j] <- anova(lmer_genotyped,lmer_null)[2,8]

beta_imputed_mat[j] <- results_imputed$coefficient[2,1]

se_imputed_mat[j] <- results_imputed$coefficient[2,2]

pval_imputed_mat[j] <- results_imputed$coefficient[2,5]

beta_imputed_pat[j] <- results_imputed$coefficient[3,1]

se_imputed_pat[j] <- results_imputed$coefficient[3,2]

pval_imputed_pat[j] <- results_imputed$coefficient[3,5]

beta_imputed_fet[j] <- results_imputed$coefficient[4,1]

se_imputed_fet[j] <- results_imputed$coefficient[4,2]

pval_imputed_fet[j] <- results_imputed$coefficient[4,5]

pval_imputed_omni[j] <- anova(lmer_imputed,lmer_null)[2,8]

beta_fet[j] <- results_hsib$coefficient[2,1]

se_fet[j] <- results_hsib$coefficient[2,2]

pval_fet[j] <- results_hsib$coefficient[2,5]

}

**Male sibling pairs at X chromosomal loci**

library(lmerTest)

library(MASS)

rm(list=ls())

Nrep = 1000	#Number of simulation replicates

N = 2000	#Sample size

Vm = 0.001	#Variance explained by maternal effect

Vp = 0.00	#Variance explained by paternal effect

Vf = 0.001	#Variance explained by fetal effect

p = 0.1	#Increaser allele frequency

Opp = FALSE	#Boolean variable to denote whether maternal and fetal effects in opposite directions (TRUE = opposite directions)

rho = 0.2	#Covariance between sibling residual variances

q <- 1-p	#Decreaser allele frequency

#Create vectors to save results from regression analyses

#For analysis using simulated genotypes:

beta_genotyped_mat <- vector(length = Nrep)	#Fitting regression with maternal and paternal genotypes known (maternal coefficient)

beta_genotyped_fet <- vector(length = Nrep)	#Fitting regression with maternal and paternal genotypes known (fetal coefficient)

se_genotyped_mat <- vector(length = Nrep)

se_genotyped_fet <- vector(length = Nrep)

pval_genotyped_mat <- vector(length = Nrep)

pval_genotyped_fet <- vector(length = Nrep)

#Analysis using imputed maternal and paternal genotypes:

beta_imputed_mat <- vector(length = Nrep)	#Fitting regression with genotypes imputed (maternal coefficient)

beta_imputed_fet <- vector(length = Nrep)	#Fitting regression with genotypes imputed (fetal coefficient)

se_imputed_mat <- vector(length = Nrep)

se_imputed_fet <- vector(length = Nrep)

pval_imputed_mat <- vector(length = Nrep)

pval_imputed_fet <- vector(length = Nrep)

#Analysis using only sibling genotypes:

beta_sib <- vector(length = Nrep)	#Fitting incorrect model including only siblings

se_sib <- vector(length = Nrep)

pval_sib <- vector(length = Nrep)

#ANOVA for full model against null model

pval_genotyped_omni <- vector(length = Nrep)

pval_imputed_omni <- vector(length = Nrep)

a <- sqrt(1/(2*p*q))	#Create genetic variable of variance one for maternal SNPs on X chromosome. Assume no dominance.

BetaM <- sqrt(Vm)	#Path coefficient for maternal effect

BetaP <- sqrt(Vp/2)	#Path coefficient for paternal effect

BetaF <- sqrt(Vf/2)	#Path coefficient for fetal effect

if(Opp == TRUE) {	#If Opp is true then maternal and fetal effects have opposite directions of effect

BetaF = -BetaF

}

Ve_y <- (1 - BetaM^2 - BetaF^2*2 - BetaP^2*2 - 2*1*BetaM*BetaF) #Residual variance in trait for male sib

for(j in 1:Nrep) {

#Sample mothers' genotypes

Zm <- sample(x = c('AA','Aa','aa'), size = N, replace = TRUE, prob = c(p^2, 2*p*q, q^2))

#Sample fathers' genotypes

Zp <- sample(x = c('A','a'), size = N, replace = TRUE, prob = c(p, q))

Z_sib1y <- vector(length = N)

Z_sib2y <- vector(length = N)

Zmyy_imp <- vector(length = N)	#Imputed vector of genotypes at mum's locus

#Simulate male sib1 genotype

r <- runif(N)

for (i in 1:N) {

Z_sib1y[i] <- switch(Zm[i],

'AA'='A',

'Aa'=ifelse(r[i] <= 0.5, 'A', 'a'),

'aa'='a')

}

#Simulate male sib2 genotype

r <- runif(N)

for (i in 1:N) {

Z_sib2y[i] <- switch(Zm[i],

'AA'='A',

'Aa'=ifelse(r[i] <= 0.5, 'A', 'a'),

'aa'='a')

}

#Change coding to 0, 1 and 2 in order to calculate allele frequencies

Z_sib1y_01 <- ifelse(Z_sib1y=='A', 0, 2)

Z_sib2y_01 <- ifelse(Z_sib2y=='A', 0, 2)

#Calculate observed allele frequency based on sibling one's genotypes

q_obs <- sum(c(Z_sib1y_01, Z_sib2y_01))/(2*N*2)

p_obs <- 1 - q_obs

#Impute parental genotypes from male-female sibs

#Calculate conditional probability of mother's genotypes given sibs genotypes

p_AA_A_A = (2*p_obs)/(p_obs+1)	#P(Mum = AA | Sib1y = A and Sib2y = A)

p_Aa_A_A = (1-p_obs)/(p_obs+1)	#P(Mum = Aa | Sib1y = A and Sib2y = A)

p_aa_A_A = 0	#P(Mum = aa | Sib1y = A and Sib2y = A)

p_AA_A_a = 0	#P(Mum = AA | Sib1y = A and Sib2y = a)

p_Aa_A_a = 1	#P(Mum = Aa | Sib1y = A and Sib2y = a)

p_aa_A_a = 0	#P(Mum = aa | Sib1y = A and Sib2y = a)

p_AA_a_A = p_AA_A_a

p_Aa_a_A = p_Aa_A_a

p_aa_a_A = p_aa_A_a

p_AA_a_a = 0	#P(Mum = AA | Sib1y = a and Sib2y = a)

p_Aa_a_a = (1-q_obs)/(q_obs+1)	#P(Mum = Aa | Sib1y = a and Sib2y = a)

p_aa_a_a = (2*q_obs)/(q_obs+1)	#P(Mum = aa | Sib1y = a and Sib2y = a)

for (i in 1:N) {

Zmyy_imp[i] <- switch(paste(Z_sib1y[i],Z_sib2y[i]),

'A A'= -a*p_AA_A_A + 0*p_Aa_A_A + a*p_aa_A_A,

'A a'= -a*p_AA_A_a + 0*p_Aa_A_a + a*p_aa_A_a,

'a A'= -a*p_AA_a_A + 0*p_Aa_a_A + a*p_aa_a_A,

'a a'= -a*p_AA_a_a + 0*p_Aa_a_a + a*p_aa_a_a)

}

#Convert AA/Aa/aa to genetic value

Zm <- ifelse(Zm=='AA', -a, ifelse(Zm=='Aa', 0, a))

Zp <- ifelse(Zp=='A', -a, a)

Z_sib1y <- ifelse(Z_sib1y=='A', -a, a)

Z_sib2y <- ifelse(Z_sib2y=='A', -a, a)

#Create correlated error variables for sib 1 and sib 2 in the model

Sigma_yy <- matrix(c(Ve_y, rho, rho, Ve_y),2,2) #Male-male sibs

e_yy <- mvrnorm(n = N, mu = c(0, 0), Sigma_yy)

#Simulate offspring outcome

Y_sib1y_yy <- BetaM*Zm + BetaP*Zp + BetaF*Z_sib1y + e_yy[,1]

Y_sib2y_yy <- BetaM*Zm + BetaP*Zp + BetaF*Z_sib2y + e_yy[,2]

test <- data.frame(rbind(cbind(1:N, Y_sib1y_yy, Zm, Zp, Zmyy_imp, Z_sib1y),

cbind(1:N, Y_sib2y_yy, Zm, Zp, Zmyy_imp, Z_sib2y)))

colnames(test) <- c("fam", "Y", "Zm", "Zp", "Zmyy_imp", "Zy")

#Run analyses

lmer_genotyped <- lmer(Y ~ Zm + Zy + (1|fam), data = test, REML = FALSE); results_genotyped <- summary(lmer_genotyped)

lmer_imputed <- lmer(Y ~ Zmyy_imp + Zy + (1|fam), data = test, REML = FALSE); results_imputed <- summary(lmer_imputed)

lmer_sib <- lmer(Y ~ Zy + (1|fam), data = test, REML = FALSE); results_sib <- summary(lmer_sib)

lmer_null <- lmer(Y ~ (1|fam), data = test, REML = FALSE)

beta_genotyped_mat[j] <- results_genotyped$coefficient[2,1]

se_genotyped_mat[j] <- results_genotyped$coefficient[2,2]

pval_genotyped_mat[j] <- results_genotyped$coefficient[2,5]

beta_genotyped_fet[j] <- results_genotyped$coefficient[3,1]

se_genotyped_fet[j] <- results_genotyped$coefficient[3,2]

pval_genotyped_fet[j] <- results_genotyped$coefficient[3,5]

pval_genotyped_omni[j] <- anova(lmer_genotyped,lmer_null)[2,8]

beta_imputed_mat[j] <- results_imputed$coefficient[2,1]

se_imputed_mat[j] <- results_imputed$coefficient[2,2]

pval_imputed_mat[j] <- results_imputed$coefficient[2,5]

beta_imputed_fet[j] <- results_imputed$coefficient[3,1]

se_imputed_fet[j] <- results_imputed$coefficient[3,2]

pval_imputed_fet[j] <- results_imputed$coefficient[3,5]

pval_imputed_omni[j] <- anova(lmer_imputed,lmer_null)[2,8]

beta_sib[j] <- results_sib$coefficient[2,1]

se_sib[j] <- results_sib$coefficient[2,2]

pval_sib[j] <- results_sib$coefficient[2,5]

}

**Female sibling pairs at X chromosomal loci**

library(lmerTest)

library(MASS)

rm(list=ls())

Nrep = 1000	#Number of simulation replicates

N = 2000	#Sample size

Vm = 0.001	#Variance explained by maternal effect

Vp = 0.001	#Variance explained by paternal effect

Vf = 0.001	#Variance explained by fetal effect

p = 0.1	#Increaser allele frequency

Opp = FALSE	#Boolean variable to denote whether maternal and fetal effects in opposite directions (TRUE = opposite directions)

rho = 0.2	#Covariance between sibling residual variances

q <- 1-p	#Decreaser allele frequency

#Create vectors to save results from regression analyses

#For analysis using simulated genotypes

beta_genotyped_mat <- vector(length = Nrep) #Fitting regression with maternal and paternal genotypes known (maternal coefficient)

beta_genotyped_pat <- vector(length = Nrep) #Fitting regression with maternal and paternal genotypes known (paternal coefficient)

beta_genotyped_fet <- vector(length = Nrep) #Fitting regression with maternal and paternal genotypes known (fetal coefficient)

se_genotyped_mat <- vector(length = Nrep)

se_genotyped_pat <- vector(length = Nrep)

se_genotyped_fet <- vector(length = Nrep)

pval_genotyped_mat <- vector(length = Nrep)

pval_genotyped_pat <- vector(length = Nrep)

pval_genotyped_fet <- vector(length = Nrep)

#Analysis using imputed maternal and paternal genotypes

beta_imputed_mat <- vector(length = Nrep)	#Fitting regression with genotypes imputed (maternal coefficient)

beta_imputed_pat <- vector(length = Nrep)	#Fitting regression with genotypes imputed (paternal coefficient)

beta_imputed_fet <- vector(length = Nrep)	#Fitting regression with genotypes imputed (fetal coefficient)

se_imputed_mat <- vector(length = Nrep)

se_imputed_pat <- vector(length = Nrep)

se_imputed_fet <- vector(length = Nrep)

pval_imputed_mat <- vector(length = Nrep)

pval_imputed_pat <- vector(length = Nrep)

pval_imputed_fet <- vector(length = Nrep)

#Analysis using only sibling genotypes

beta_sib <- vector(length = Nrep)	#Fitting incorrect model including only siblings

se_sib <- vector(length = Nrep)

pval_sib <- vector(length = Nrep)

#ANOVA for full model against null model

pval_genotyped_omni <- vector(length = Nrep)

pval_imputed_omni <- vector(length = Nrep)

a <- sqrt(1/(2*p*q)) #Create genetic variable of variance one for maternal and fetal SNPs on X chromosome. Assume no dominance.

BetaM <- sqrt(Vm) #Path coefficient for maternal effect

BetaP <- sqrt(Vp/2) #Path coefficient for paternal effect

BetaF <- sqrt(Vf) #Path coefficient for fetal effect

if(Opp == TRUE) { #If Opp is true then maternal and fetal effects have opposite directions of effect

BetaF = -BetaF

}

Ve_x <- (1 - BetaM^2 - BetaF^2 - BetaP^2*2 - 2*0.5*BetaM*BetaF - 2*0.5*BetaP*BetaF*2) #Residual variance in trait for female sib

for(j in 1:Nrep) {

#Sample mothers' genotypes

Zm <- sample(x = c('AA','Aa','aa'), size = N, replace = TRUE, prob = c(p^2, 2*p*q, q^2))

#Sample fathers' genotypes

Zp <- sample(x = c('A','a'), size = N, replace = TRUE, prob = c(p, q))

Z_sib1x <- vector(length = N)

Z_sib2x <- vector(length = N)

Zmxx_imp <- vector(length = N)	#Imputed vector of genotypes at mum's locus

Zpxx_imp <- vector(length = N)	#Imputed vector of genotypes at dad's locus

#Simulate female sib 1 genotype

r <- runif(N)

for (i in 1:N) {

Z_sib1x[i] <- switch(paste(Zm[i],Zp[i]),

'AA A'='AA',

'AA a'='Aa',

'Aa A'=ifelse(r[i] <= 0.5, 'AA', 'Aa'),

'Aa a'=ifelse(r[i] <= 0.5, 'Aa', 'aa'),

'aa A'='Aa',

'aa a'='aa')

}

#Simulate female sib 2 genotype

r <- runif(N)

for (i in 1:N) {

Z_sib2x[i] <- switch(paste(Zm[i],Zp[i]),

'AA A'='AA',

'AA a'='Aa',

'Aa A'=ifelse(r[i] <= 0.5, 'AA', 'Aa'),

'Aa a'=ifelse(r[i] <= 0.5, 'Aa', 'aa'),

'aa A'='Aa',

'aa a'='aa')

}

#Change coding to 0, 1 and 2 in order to calculate allele frequencies

Z_sib1x_012 <- ifelse(Z_sib1x=='AA', 0,

ifelse(Z_sib1x=='Aa', 1, 2))

Z_sib2x_012 <- ifelse(Z_sib2x=='AA', 0,

ifelse(Z_sib2x=='Aa', 1, 2))

#Calculate observed allele frequency based on sibling genotypes

q_obs <- sum(Z_sib1x_012, Z_sib2x_012)/(2*N*2)

p_obs <- 1 - q_obs

#Impute parental genotypes from female sibs

#Calculate conditional probability of mother's genotypes given sibs genotypes

p_AA_AA_AA = (2*p_obs)/(p_obs+1)	#P(Mum = AA | Sibx = AA and Sibx = AA)

p_Aa_AA_AA = (1-p_obs)/(p_obs+1)	#P(Mum = Aa | Sibx = AA and Sibx = AA)

p_aa_AA_AA = 0	#P(Mum = aa | Sibx = AA and Sibx = AA)

p_AA_AA_Aa = 0	#P(Mum = AA | Sibx = AA and Sibx = Aa)

p_Aa_AA_Aa = 1	#P(Mum = Aa | Sibx = AA and Sibx = Aa)

p_aa_AA_Aa = 0	#P(Mum = aa | Sibx = AA and Sibx = Aa)

p_AA_AA_aa = 0	#P(Mum = AA | Sibx = AA and Sibx = aa)

p_Aa_AA_aa = 1	#P(Mum = Aa | Sibx = AA and Sibx = aa)

p_aa_AA_aa = 0	#P(Mum = aa | Sibx = AA and Sibx = aa)

p_AA_Aa_Aa = 2/3*p_obs	#P(Mum = AA | Sibx = Aa and Sibx = Aa)

p_Aa_Aa_Aa = 1/3	#P(Mum = Aa | Sibx = Aa and Sibx = Aa)

p_aa_Aa_Aa = 2/3*q_obs	#P(Mum = aa | Sibx = Aa and Sibx = Aa)

p_AA_aa_Aa = 0	#P(Mum = AA | Sibx = aa and Sibx = Aa)

p_Aa_aa_Aa = 1	#P(Mum = Aa | Sibx = aa and Sibx = Aa)

p_aa_aa_Aa = 0	#P(Mum = aa | Sibx = aa and Sibx = Aa)

p_AA_aa_aa = 0	#P(Mum = AA | Sibx = aa and Sibx = aa)

p_Aa_aa_aa = (1-q_obs)/(q_obs+1)	#P(Mum = Aa | Sibx = aa and Sibx = aa)

p_aa_aa_aa = (2*q_obs)/(q_obs+1)	#P(Mum = aa | Sibx = aa and Sibx = aa)

for (i in 1:N) {

Zmxx_imp[i] <- switch(paste(Z_sib1x[i], Z_sib2x[i]),

'AA AA'= -a*p_AA_AA_AA + 0*p_Aa_AA_AA + a*p_aa_AA_AA,

'AA Aa'= -a*p_AA_AA_Aa + 0*p_Aa_AA_Aa + a*p_aa_AA_Aa,

'AA aa'= -a*p_AA_AA_aa + 0*p_Aa_AA_aa + a*p_aa_AA_aa,

'Aa AA'= -a*p_AA_AA_Aa + 0*p_Aa_AA_Aa + a*p_aa_AA_Aa,

'Aa Aa'= -a*p_AA_Aa_Aa + 0*p_Aa_Aa_Aa + a*p_aa_Aa_Aa,

'Aa aa'= -a*p_AA_aa_Aa + 0*p_Aa_aa_Aa + a*p_aa_aa_Aa,

'aa AA'= -a*p_AA_aa_AA + 0*p_Aa_aa_AA + a*p_aa_aa_AA,

'aa Aa'= -a*p_AA_aa_Aa + 0*p_Aa_aa_Aa + a*p_aa_aa_Aa,

'aa aa'= -a*p_AA_aa_aa + 0*p_Aa_aa_aa + a*p_aa_aa_aa)

}

#Calculate conditional probability of father's genotypes given sib's genotype

p_dA_AA_AA <- 1	#P(Dad = A | Sibx = AA and Sibx = AA)

p_da_AA_AA <- 0	#P(Dad = a | Sibx = AA and Sibx = AA)

p_dA_AA_Aa <- 1	#P(Dad = A | Sibx = AA and Sibx = Aa)

p_da_AA_Aa <- 0	#P(Dad = a | Sibx = AA and Sibx = Aa)

p_dA_Aa_AA <- 1	#P(Dad = A | Sibx = Aa and Sibx = AA)

p_da_Aa_AA <- 0	#P(Dad = a | Sibx = Aa and Sibx = AA)

p_dA_Aa_Aa <- 2/3-1/3*p_obs	#P(Dad = A | Sibx = Aa and Sibx = Aa)

p_da_Aa_Aa <- 2/3-1/3*q_obs	#P(Dad = a | Sibx = Aa and Sibx = Aa)

p_dA_Aa_aa <- 0	#P(Dad = A | Sibx = Aa and Sibx = aa)

p_da_Aa_aa <- 1	#P(Dad = a | Sibx = Aa and Sibx = aa)

p_dA_aa_Aa <- 0	#P(Dad = A | Sibx = aa and Sibx = Aa)

p_da_aa_Aa <- 1	#P(Dad = a | Sibx = aa and Sibx = Aa)

p_dA_aa_aa <- 0	#P(Dad = A | Sibx = aa and Sibx = aa)

p_da_aa_aa <- 1	#P(Dad = a | Sibx = aa and Sibx = aa)

for (i in 1:N) {

Zpxx_imp[i] <- switch(paste(Z_sib1x[i], Z_sib2x[i]),

'AA AA'= -a*p_dA_AA_AA + a*p_da_AA_AA,

'Aa AA'= -a*p_dA_Aa_AA + a*p_da_Aa_AA,

'AA Aa'= -a*p_dA_AA_Aa + a*p_da_AA_Aa,

'Aa Aa'= -a*p_dA_Aa_Aa + a*p_da_Aa_Aa,

'aa Aa'= -a*p_dA_aa_Aa + a*p_da_aa_Aa,

'Aa aa'= -a*p_dA_Aa_aa + a*p_da_Aa_aa,

'aa aa'= -a*p_dA_aa_aa + a*p_da_aa_aa)

}

#Convert AA/Aa/aa to genetic value

Zm <- ifelse(Zm=='AA', -a, ifelse(Zm=='Aa', 0, a))

Zp <- ifelse(Zp=='A', -a, a)

Z_sib1x <- ifelse(Z_sib1x=='AA', -a, ifelse(Z_sib1x=='Aa', 0, a))

Z_sib2x <- ifelse(Z_sib2x=='AA', -a, ifelse(Z_sib2x=='Aa', 0, a))

#Create correlated error variables for sib 1 and sib 2 in the model

Sigma_xx <- matrix(c(Ve_x, rho, rho, Ve_x),2,2)

e_xx <- mvrnorm(n = N, mu = c(0, 0), Sigma_xx)

#Simulate offspring outcome

Y_sib1x_xx <- BetaM*Zm + BetaP*Zp + BetaF*Z_sib1x + e_xx[,1]

Y_sib2x_xx <- BetaM*Zm + BetaP*Zp + BetaF*Z_sib2x + e_xx[,2]

test <- data.frame(rbind(cbind(1:N, Y_sib1x_xx, Zm, Zp, Zmxx_imp, Zpxx_imp, Z_sib1x),

cbind(1:N, Y_sib2x_xx, Zm, Zp, Zmxx_imp, Zpxx_imp, Z_sib2x)))

colnames(test) <- c("fam", "Y_xx", "Zm", "Zp", "Zmxx_imp", "Zpxx_imp", "Z_xx")

#Run analyses

lmer_genotyped <- lmer(Y_xx ~ Zm + Zp + Z_xx + (1|fam), data = test, REML = FALSE); results_genotyped <- summary(lmer_genotyped)

lmer_imputed <- lmer(Y_xx ~ Zmxx_imp + Zpxx_imp + Z_xx + (1|fam), data = test, REML = FALSE); results_imputed <- summary(lmer_imputed)

lmer_sib <- lmer(Y_xx ~ Z_xx + (1|fam), data = test, REML = FALSE); results_sib <- summary(lmer_sib)

lmer_null <- lmer(Y_xx ~ (1|fam), data = test, REML = FALSE)

beta_genotyped_mat[j] <- results_genotyped$coefficient[2,1]

se_genotyped_mat[j] <- results_genotyped$coefficient[2,2]

pval_genotyped_mat[j] <- results_genotyped$coefficient[2,5]

beta_genotyped_pat[j] <- results_genotyped$coefficient[3,1]

se_genotyped_pat[j] <- results_genotyped$coefficient[3,2]

pval_genotyped_pat[j] <- results_genotyped$coefficient[3,5]

beta_genotyped_fet[j] <- results_genotyped$coefficient[4,1]

se_genotyped_fet[j] <- results_genotyped$coefficient[4,2]

pval_genotyped_fet[j] <- results_genotyped$coefficient[4,5]

pval_genotyped_omni[j] <- anova(lmer_genotyped,lmer_null)[2,8]

beta_imputed_mat[j] <- results_imputed$coefficient[2,1]

se_imputed_mat[j] <- results_imputed$coefficient[2,2]

pval_imputed_mat[j] <- results_imputed$coefficient[2,5]

beta_imputed_pat[j] <- results_imputed$coefficient[3,1]

se_imputed_pat[j] <- results_imputed$coefficient[3,2]

pval_imputed_pat[j] <- results_imputed$coefficient[3,5]

beta_imputed_fet[j] <- results_imputed$coefficient[4,1]

se_imputed_fet[j] <- results_imputed$coefficient[4,2]

pval_imputed_fet[j] <- results_imputed$coefficient[4,5]

pval_imputed_omni[j] <- anova(lmer_imputed,lmer_null)[2,8]

beta_sib[j] <- results_sib$coefficient[2,1]

se_sib[j] <- results_sib$coefficient[2,2]

pval_sib[j] <- results_sib$coefficient[2,5]

}

**Opposite sex sibling pairs at X chromosomal loci**

library(MASS)

library(lmerTest)

rm(list=ls())

Nrep = 1000	#Number of simulation replicates

N = 2000	#Sample size

Vm = 0.001	#Variance explained by maternal effect

Vp = 0.001	#Variance explained by paternal effect

Vf = 0.001	#Variance explained by fetal effect

p = 0.5	#Increaser allele frequency

Opp = FALSE #Boolean variable to denote whether maternal and fetal effects in opposite directions (TRUE = opposite directions)

rho = 0.2	#Covariance between sibling residual variances

q <- 1-p	#Decreaser allele frequency

#Create vectors to save results from regression analyses

#For analysis using simulated genotypes:

beta_genotyped_mat <- vector(length = Nrep) #Fitting regression with maternal and paternal genotypes known (maternal coefficient)

beta_genotyped_pat <- vector(length = Nrep) #Fitting regression with maternal and paternal genotypes known (paternal coefficient)

beta_genotyped_fet <- vector(length = Nrep) #Fitting regression with maternal and paternal genotypes known (fetal coefficient)

se_genotyped_mat <- vector(length = Nrep)

se_genotyped_pat <- vector(length = Nrep)

se_genotyped_fet <- vector(length = Nrep)

pval_genotyped_mat <- vector(length = Nrep)

pval_genotyped_pat <- vector(length = Nrep)

pval_genotyped_fet <- vector(length = Nrep)

#Analysis using imputed maternal and paternal genotypes:

beta_imputed_mat <- vector(length = Nrep) #Fitting regression with genotypes imputed (maternal coefficient)

beta_imputed_pat <- vector(length = Nrep) #Fitting regression with genotypes imputed (paternal coefficient)

beta_imputed_fet <- vector(length = Nrep) #Fitting regression with genotypes imputed (fetal coefficient)

se_imputed_mat <- vector(length = Nrep)

se_imputed_pat <- vector(length = Nrep)

se_imputed_fet <- vector(length = Nrep)

pval_imputed_mat <- vector(length = Nrep)

pval_imputed_pat <- vector(length = Nrep)

pval_imputed_fet <- vector(length = Nrep)

#Analysis using only half-sib genotypes:

beta_fet <- vector(length = Nrep) #Fitting incorrect model to just sibs

se_fet <- vector(length = Nrep)

pval_fet <- vector(length = Nrep)

#ANOVA for full model against null model

pval_imputed_omni <- vector(length = Nrep)

pval_genotyped_omni <- vector(length = Nrep)

a <- sqrt(1/(2*p*q))	#Create genetic variable of variance one for maternal SNPs on X chromosome. Assume no dominance.

BetaM <- sqrt(Vm)	#Path coefficient for maternal effect

BetaP <- sqrt(Vp/2)	#Path coefficient for paternal effect

BetaF <- sqrt(Vf/2)	#Path coefficient for fetal effect. Assume **male** genotype

if(Opp == TRUE) {	#If Opp is true then maternal and fetal effects have opposite directions of effect

BetaF = -BetaF

}

Ve_x <- (1 - BetaM^2 - BetaF^2 - BetaP^2*2 - 2*0.5*BetaM*BetaF - 2*0.5*BetaP*BetaF*2) #Residual variance in trait for female sib

Ve_y <- (1 - BetaM^2 - BetaF^2*2 - BetaP^2*2 - 2*1*BetaM*BetaF) #Residual variance in trait for male sib

for(j in 1:Nrep) {

#Sample mothers' genotypes

Zm <- sample(x = c('AA','Aa','aa'), size = N, replace = TRUE, prob = c(p^2, 2*p*q, q^2))

#Sample fathers' genotypes

Zp <- sample(x = c('A','a'), size = N, replace = TRUE, prob = c(p, q))

Z_sibx <- vector(length = N)

Z_siby <- vector(length = N)

Zmxy_imp <- vector(length = N) #Imputed vector of genotypes at mum's locus

Zpxy_imp <- vector(length = N) #Imputed vector of genotypes at dad's locus

#Simulate female sib genotype

r <- runif(N)

for (i in 1:N) {

Z_sibx[i] <- switch(paste(Zm[i],Zp[i]),

'AA A'='AA',

'AA a'='Aa',

'Aa A'=ifelse(r[i] <= 0.5, 'AA', 'Aa'),

'Aa a'=ifelse(r[i] <= 0.5, 'Aa', 'aa'),

'aa A'='Aa',

'aa a'='aa')

}

#Simulate male sib genotype

r <- runif(N)

for (i in 1:N) {

Z_siby[i] <- switch(Zm[i],

'AA'='A',

'Aa'=ifelse(r[i] <= 0.5, 'A', 'a'),

'aa'='a')

}

#Change coding to 0, 1 and 2 in order to calculate allele frequencies

Z_sibx_012 <- ifelse(Z_sibx=='AA', 0, ifelse(Z_sibx=='Aa', 1, 2))

Z_siby_01 <- ifelse(Z_siby=='A', 0, 1)

#Calculate observed allele frequency based on female sibling one's genotypes

q_obs <- sum(Z_sibx_012)/(2*N)

p_obs <- 1 - q_obs

#Impute parental genotypes from male-female siblings

#Calculate conditional probability of mother's genotypes given sibling genotypes

p_AA_A_AA = (2*p_obs)/(p_obs+1)	#P(Mum = AA | Siby = A and Sibx = AA)

p_Aa_A_AA = (1-p_obs)/(p_obs+1)	#P(Mum = Aa | Siby = A and Sibx = AA)

p_aa_A_AA = 0	#P(Mum = aa | Siby = A and Sibx = AA)

p_AA_A_Aa = (2*p_obs)/(2*p_obs+1)	#P(Mum = AA | Siby = A and Sibx = Aa)

p_Aa_A_Aa = 1/(2*p_obs+1)	#P(Mum = Aa | Siby = A and Sibx = Aa)

p_aa_A_Aa = 0	#P(Mum = aa | Siby = A and Sibx = Aa)

p_AA_A_aa = 0	#P(Mum = AA | Siby = A and Sibx = aa)

p_Aa_A_aa = 1	#P(Mum = Aa | Siby = A and Sibx = aa)

p_aa_A_aa = 0	#P(Mum = aa | Siby = A and Sibx = aa)

p_AA_a_AA = 0	#P(Mum = AA | Siby = a and Sibx = AA)

p_Aa_a_AA = 1	#P(Mum = Aa | Siby = a and Sibx = AA)

p_aa_a_AA = 0	#P(Mum = aa | Siby = a and Sibx = AA)

p_AA_a_Aa = 0	#P(Mum = AA | Siby = a and Sibx = Aa)

p_Aa_a_Aa = 1/(2*q_obs+1)	#P(Mum = Aa | Siby = a and Sibx = Aa)

p_aa_a_Aa = (2*q_obs)/(2*q_obs+1)	#P(Mum = aa | Siby = a and Sibx = Aa)

p_AA_a_aa = 0	#P(Mum = AA | Siby = a and Sibx = aa)

p_Aa_a_aa = (1-q_obs)/(q_obs+1)	#P(Mum = Aa | Siby = a and Sibx = aa)

p_aa_a_aa = (2*q_obs)/(q_obs+1)	#P(Mum = aa | Siby = a and Sibx = aa)

p_AA_AA_A <- p_AA_A_AA

p_Aa_AA_A <- p_Aa_A_AA

p_aa_AA_A <- p_aa_A_AA

p_AA_Aa_A <- p_AA_A_Aa

p_Aa_Aa_A <- p_Aa_A_Aa

p_aa_Aa_A <- p_aa_A_Aa

p_AA_aa_A <- p_AA_A_aa

p_Aa_aa_A <- p_Aa_A_aa

p_aa_aa_A <- p_aa_A_aa

p_AA_AA_a <- p_AA_a_AA

p_Aa_AA_a <- p_Aa_a_AA

p_aa_AA_a <- p_aa_a_AA

p_AA_Aa_a <- p_AA_a_Aa

p_Aa_Aa_a <- p_Aa_a_Aa

p_aa_Aa_a <- p_aa_a_Aa

p_AA_aa_a <- p_AA_a_aa

p_Aa_aa_a <- p_Aa_a_aa

p_aa_aa_a <- p_aa_a_aa

for (i in 1:N) {

Zmxy_imp[i] <- switch(paste(Z_siby[i], Z_sibx[i]),

'A AA'= -a*p_AA_A_AA + 0*p_Aa_A_AA + a*p_aa_A_AA,

'A Aa'= -a*p_AA_A_Aa + 0*p_Aa_A_Aa + a*p_aa_A_Aa,

'A aa'= -a*p_AA_A_aa + 0*p_Aa_A_aa + a*p_aa_A_aa,

'a AA'= -a*p_AA_a_AA + 0*p_Aa_a_AA + a*p_aa_a_AA,

'a Aa'= -a*p_AA_a_Aa + 0*p_Aa_a_Aa + a*p_aa_a_Aa,

'a aa'= -a*p_AA_a_aa + 0*p_Aa_a_aa + a*p_aa_a_aa)

}

#Calculate conditional probability of father's genotypes given sib's genotype

p_A_A_AA <- 1	#P(Dad = A | Siby = A and Sibx = AA)

p_a_A_AA <- 0	#P(Dad = a | Siby = A and Sibx = AA)

p_A_A_Aa <- p_obs/(2*p_obs+1)	#P(Dad = A | Siby = A and Sibx = Aa)

p_a_A_Aa <- (p_obs+1)/(2*p_obs+1)	#P(Dad = a | Siby = A and Sibx = Aa)

p_A_A_aa <- 0	#P(Dad = A | Siby = A and Sibx = aa)

p_a_A_aa <- 1	#P(Dad = a | Siby = A and Sibx = aa)

p_A_a_AA <- 1	#P(Dad = A | Siby = a and Sibx = AA)

p_a_a_AA <- 0	#P(Dad = a | Siby = a and Sibx = AA)

p_A_a_Aa <- (q_obs+1)/(2*q_obs+1)	#P(Dad = A | Siby = a and Sibx = Aa)

p_a_a_Aa <- q_obs/(2*q_obs+1)	#P(Dad = a | Siby = a and Sibx = Aa)

p_A_a_aa <- 0	#P(Dad = A | Siby = a and Sibx = aa)

p_a_a_aa <- 1	#P(Dad = a | Siby = a and Sibx = aa)

for (i in 1:N) {

Zpxy_imp[i] <- switch(paste(Z_siby[i], Z_sibx[i]),

'A AA'= -a*p_A_A_AA + a*p_a_A_AA,

'A Aa'= -a*p_A_A_Aa + a*p_a_A_Aa,

'A aa'= -a*p_A_A_aa + a*p_a_A_aa,

'a AA'= -a*p_A_a_AA + a*p_a_a_AA,

'a Aa'= -a*p_A_a_Aa + a*p_a_a_Aa,

'a aa'= -a*p_A_a_aa + a*p_a_a_aa)

}

#Convert AA/Aa/aa to genetic value -a/0/a

Zm <- ifelse(Zm=='AA', -a, ifelse(Zm=='Aa', 0, a))

Zp <- ifelse(Zp=='A', -a, a)

Z_sibx <- ifelse(Z_sibx=='AA', -a, ifelse(Z_sibx=='Aa', 0, a))

Z_siby <- ifelse(Z_siby=='A', -a, a)

#Create correlated error variables for sib 1 and sib 2 in the model

Sigma_xy <- matrix(c(Ve_x, rho, rho, Ve_y),2,2) #Female-male sibs

e_xy <- mvrnorm(n = N, mu = c(0, 0), Sigma_xy)

#Simulate offspring outcome

Y_sibx_xy <- BetaM*Zm + BetaP*Zp + BetaF*Z_sibx + e_xy[,1]

Y_siby_xy <- BetaM*Zm + BetaP*Zp + BetaF*Z_siby + e_xy[,2]

testxy <- data.frame(rbind(cbind(1:N, Y_sibx_xy, Zm, Zp, Zmxy_imp, Zpxy_imp, Z_sibx),

cbind(1:N, Y_siby_xy, Zm, Zp, Zmxy_imp, Zpxy_imp, Z_siby)))

colnames(testxy) <- c("fam", "Y_xy", "Zm", "Zp", "Zmxy_imp", "Zpxy_imp", "Z_xy")

#Run analyses

lmer_genotyped <- lmer(Y_xy ~ Zm + Zp + Z_xy + (1|fam), data = testxy, REML = FALSE); results_genotyped <- summary(lmer_genotyped)

lmer_imputed <- lmer(Y_xy ~ Zmxy_imp + Zpxy_imp + Z_xy + (1|fam), data = testxy, REML = FALSE); results_imputed <- summary(lmer_imputed)

lmer_sib <- lmer(Y_xy ~ Z_xy + (1|fam), data = testxy, REML = FALSE); results_sib <- summary(lmer_sib)

lmer_null <- lmer(Y_xy ~ (1|fam), data = testxy, REML = FALSE)

beta_genotyped_mat[j] <- results_genotyped$coefficient[2,1]

se_genotyped_mat[j] <- results_genotyped$coefficient[2,2]

pval_genotyped_mat[j] <- results_genotyped$coefficient[2,5]

beta_genotyped_pat[j] <- results_genotyped$coefficient[3,1]

se_genotyped_pat[j] <- results_genotyped$coefficient[3,2]

pval_genotyped_pat[j] <- results_genotyped$coefficient[3,5]

beta_genotyped_fet[j] <- results_genotyped$coefficient[4,1]

se_genotyped_fet[j] <- results_genotyped$coefficient[4,2]

pval_genotyped_fet[j] <- results_genotyped$coefficient[4,5]

pval_genotyped_omni[j] <- anova(lmer_genotyped,lmer_null)[2,8]

beta_imputed_mat[j] <- results_imputed$coefficient[2,1]

se_imputed_mat[j] <- results_imputed$coefficient[2,2]

pval_imputed_mat[j] <- results_imputed$coefficient[2,5]

beta_imputed_pat[j] <- results_imputed$coefficient[3,1]

se_imputed_pat[j] <- results_imputed$coefficient[3,2]

pval_imputed_pat[j] <- results_imputed$coefficient[3,5]

beta_imputed_fet[j] <- results_imputed$coefficient[4,1]

se_imputed_fet[j] <- results_imputed$coefficient[4,2]

pval_imputed_fet[j] <- results_imputed$coefficient[4,5]

pval_imputed_omni[j] <- anova(lmer_imputed,lmer_null)[2,8]

beta_fet[j] <- results_sib$coefficient[2,1]

se_fet[j] <- results_sib$coefficient[2,2]

pval_fet[j] <- results_sib$coefficient[2,5]

}
